# Employing RNA editing to engineer personalized tumor-specific neoantigens (editopes)

**DOI:** 10.1101/2023.03.16.532918

**Authors:** Riccardo Pecori, Beatrice Casati, Rona Merdler-Rabinowicz, Netanel Landesman, Khwab Sanghvi, Stefan Zens, Kai Kipfstuhl, Veronica Pinamonti, Annette Arnold, John M. Lindner, Michael Platten, Rienk Offringa, Rafael Carretero, Eytan Ruppin, Erez Y. Levanon, Fotini Nina Papavasiliou

**Author notes:** These authors contributed equally to this work (Riccardo Pecori, Beatrice Casati, Rona Merdler-Rabinowicz).

## Abstract

Increasing the quantity and immunogenicity of neoantigens in tumors is essential for advancing immunotherapy. However, engineering neoantigens remains challenging due to the need for precise, tumor-specific antigen modification without affecting normal cells. To tackle this challenge, we developed *Short Precise-Encodable ADAR Recruiting* (SPEAR) ADAR-engagers, an approach that uses short guide RNAs to engage the endogenous RNA editor ADAR1 and direct it to regions of mRNA targets known to encode MHC-presented peptides. By precisely editing adenosine-to-inosine (A-to-I) in these contexts, we effectively mutate specific epitopes into neoepitopes (which we now term “editopes”). As proof of concept, we targeted the known antigen MART-1 (Melanoma-Associated Antigen Recognized by T cells-1), and demonstrated that guided ADAR1 editing can generate immunogenic epitopes that activate T cells and promote tumor cell elimination. Building on this concept, we developed a computational pipeline to identify tumor-specific somatic mutations suitable for SPEAR-mediated editing. This strategy enables selective neoantigen generation in cancer cells, effectively increasing their apparent tumor mutational burden and potentially enhancing their susceptibility to immunotherapy.

## Introduction

In recent years, immunotherapy has emerged as a revolutionary approach in cancer therapy, demonstrating remarkable clinical response in numerous cases with acceptable patient risk^1^. This success depends on the ability to trigger a robust immune response against the tumor, and requires both the presence of cancer-specific antigens on tumor cells and T lymphocytes bearing antigen-specific T cell receptors (TCR) that recognize and kill them. Responsiveness to immunotherapy is positively correlated with a tumor’s mutational burden (TMB)^2–4^. Since only a subset of somatic mutations give rise to neoepitopes that are processed and presented as non-self by the major histocompatibility complex (MHC), tumors with a high TMB are more likely to produce immunogenic neoantigens and to respond favorably to immunotherapy^5^.

Neoantigens arise from non-synonymous DNA mutations, but also from aberrant post-transcriptional and post-translational modifications, including aberrant mRNA translation, alternative splicing, and RNA editing^6–8^. However, many cancers harbor an insufficient repertoire of neoantigens capable of eliciting a strong immune reaction^9^ and are therefore poorly responsive to immunotherapy. To address this challenge, strategies aimed at artificially enhancing neoantigen production have emerged, such as modulation of alternative splicing^10^. Yet, these approaches are not tumor-specific and can often also affect healthy tissues, potentially leading to adverse side effects, such as autoimmunity. Effective methods to generate specific neoantigens exclusively within tumor cells are still lacking.

Here, we engineer tumor-specific neoantigens by employing ADAR-recruiting gRNAs (reviewed in *REFs*^11–13^) to introduce precise and controlled RNA edits specifically in transcripts exclusively present in cancer cells, thereby generating immunogenic peptides selectively in malignant tissues. Physiologically, Adenosine Deaminase Acting on RNA 1 (ADAR1) is an enzyme that recognizes long double-stranded RNA (dsRNA) structures and converts adenosines (A) to inosines (I), which are interpreted as guanosines (G) by the cellular translation machineries^14,15^ (an attribute that has enabled the development of guided editing through the engagement of the endogenous enzyme by provision of an antisense guide RNA - an “ADAR engager”). ADAR1 activity, which is already prolific across the mammalian transcriptome^14^, increases in almost all cancers^16^, contributing to transcriptomic diversity and neoantigen generation^7^.

While ADAR engagers have been explored in the clinic for the correction of disease-causing mutations such as premature stop codons, we hypothesized that this same mechanism could be repurposed to deliberately introduce precise nucleotide substitutions into cancer cell mRNAs to generate tumor-specific neoantigens. To this end, we set out to develop Short Precise-Encodable ADAR Recruiting (SPEAR) ADAR-engagers; compact, genetically encodable guide RNAs specifically designed to recruit endogenous ADAR1 and drive A-to-I RNA editing in a highly controlled manner. To evaluate the feasibility of this approach, we selected a well-characterized tumor antigen, MART-1 (Melanoma-Associated Antigen Recognized by T cells-1). We engineered a non-immunogenic variant of it as a model system to assess whether targeted A-to-I editing could restore T cell recognition and cytotoxicity, thereby demonstrating the potential of RNA editing to induce functional neoantigen formation directly from endogenous transcripts.

Building on this conceptual framework, we also designed a computational pipeline to systematically identify tumor-specific somatic mutations that could serve as substrates for SPEAR-mediated editing. Because these editing events are designed to target only transcripts harboring tumor-specific mutations, the strategy offers the potential for tumor-selective neoantigen formation with minimal risk to healthy tissues. This approach provides a personalized and safe avenue for enhancing tumor immunogenicity and expanding the repertoire of actionable neoantigens for cancer immunotherapy.

## Results

### Optimization of Short Precise-Encodable ADAR1 Recruiting (SPEAR) gRNAs

Several methods have enabled precise site-directed RNA editing using engineered or endogenous editing enzymes. A particularly useful approach relies on ADAR-engagers that base-pair with target transcripts to form dsRNA structures capable of recruiting endogenous ADAR1 to specific RNA sites^17^. This method offers several therapeutic advantages: it eliminates the need for exogenous protein (these programmed oligonucleotides can be delivered directly into cells), reduces the molecular payload (ADAR-engagers consisting of merely a few dozen nucleotides (nt) prove effective variants^18–21^, and minimizes the risk of an anti-drug immune response (allowing for repeat dosing).

ADAR-engagers can be delivered as chemically modified antisense oligonucleotides (ASOs), as genetically encoded constructs via plasmids or viral particles, or as in vitro transcribed RNA moieties. Upon hybridization with target mRNAs, gRNAs form duplexes that recruit endogenous ADAR enzymes to catalyze A-to-I editing^19–27^. These gRNAs generally fall into two main categories: (1) long gRNAs (>100 nt) with extended antisense “specificity domains” for target pairing^20,22,24^, and (2) shorter, structured gRNAs that include a compact specificity domain along with a “recruitment domain” RNA motif to enhance ADAR recruitment^19,21,23,28^. Long gRNAs are prone to extensive bystander off-target editing, requiring time-consuming, target-specific optimization^20,22,24^. In contrast, short, structured gRNAs exhibit high precision and editing efficiency when chemically modified and delivered as ASOs^18,21^, but show poor performance when genetically encoded^29^. More complex designs using clustered ADAR recruitment motifs offer improved specificity, though they involve intricate design rules and require non-intuitive computational tools^19,23^. To overcome these challenges, we sought to optimize a genetically encodable gRNA that could efficiently and precisely recruit endogenous ADAR1. Our goal was to minimize the gRNA structure required for efficient ADAR1 recruitment, thereby developing a simple and flexible platform for therapeutic deployment. Such optimized gRNAs could be delivered as plasmid- or virus-encoded constructs, or alternatively as chemically stabilized RNA molecules for direct administration.

To evaluate the capability of such gRNAs to recruit endogenous ADARs, we generated a dual-reporter HEK293T cell line stably expressing mCherry and a mutated eGFP containing a premature stop codon (W58X) linked by a P2A self-cleaving peptide^30^. In the absence of editing, only mCherry is expressed. Successful recruitment of ADAR to the UAG codon converts this stop codon to UIG, which is interpreted as a tryptophan codon (UGG), thereby restoring eGFP fluorescence (**Figure 1A** and *REF*^31^). This system enabled rapid, quantitative assessment of gRNA activity via flow cytometry. Additionally, HEK293T cells express ADAR1 but not ADAR2^20^, allowing us to specifically assess ADAR1-dependent editing. Notably, the use of a stably integrated reporter mimics the behavior of an endogenous transcript more accurately than co-transfection of guide and target RNAs, thereby offering a more realistic and relevant testing environment.

**Figure 1.**
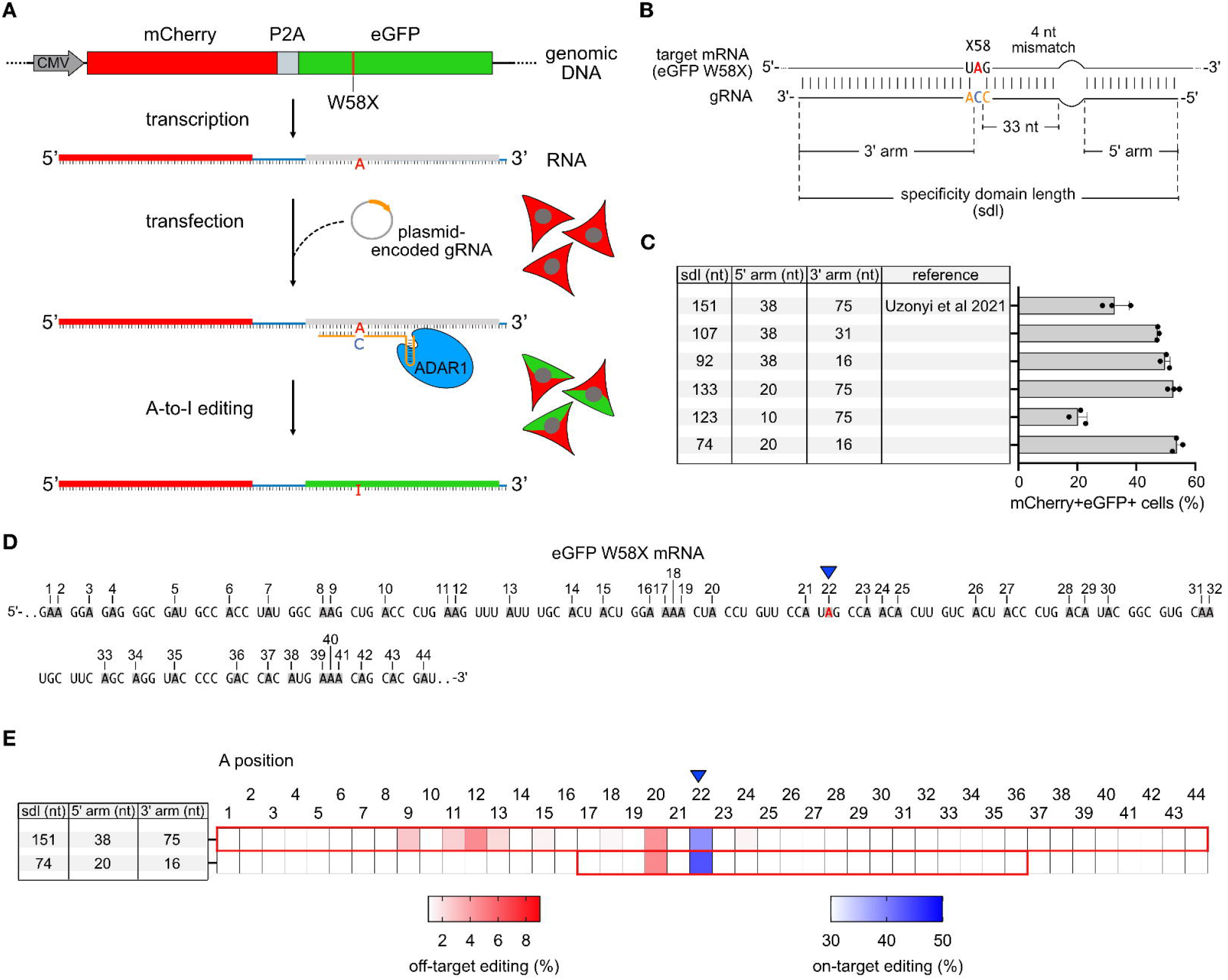
Short Precise Encodable ADAR Recruiting (SPEAR) gRNAs enable efficient and precise site-directed RNA editing. **(A)** Schematic of the dual reporter cell line and the experimental setup used to screen different genetically encoded gRNAs. **(B)** Diagram of the eGFP W58X target mRNA and the corresponding targeting gRNA used in the experiment shown in panel **C.** (**C**) Flow cytometry analysis of mCherry and eGFP double-positive cells, presented as a bar plot for the tested gRNAs. (**D**) Schematic of the sequencing window where A-to-I edits were assessed, as shown in panel **E**. (**E**) Heatmap of A-to-I editing from NGS amplicon sequencing, showing on-target editing (in blue) and off-target editing (in red). The region targeted by each gRNA is indicated with a red rectangle. For panel C, bars represent the mean for three biological replicates; error bars indicate the standard deviation.

We began by examining the impact of three structural features of encoded gRNAs: (1) specificity domain length, (2) the distance between the target adenosine and the recruitment domain, and (3) the recruitment domain itself (**Figure S1A**). Consistent with prior work, we confirmed that long, unstructured gRNAs were more effective than structured gRNAs when expressed from plasmids (**Figure S1B**), and that editing efficiency improved upon inclusion of a 4-nt mismatch placed 35 nt downstream of the target adenosine^32^ (**Figure S1B,C**). However, amplicon sequencing revealed that longer gRNAs incurred greater off-target editing across adjacent adenosines, validating concerns about precision and bystander activity (**Figure S1D-E**). This analysis also confirmed the flow cytometry results for on-target editing, validating the robustness of our dual-reporter cell line for gRNA screening (**Figure S1D**).

To systematically optimize gRNA architecture, we designed a series of truncated gRNAs based on the long LEAPER construct with a 4-nt mismatch. We varied the length of the 5⍰ and 3⍰ arms independently (**Figure 1B,C**). Reducing the 3⍰ arm from 75 to 16 nt resulted in a 1.5-fold increase in mCherry^+^eGFP^+^ cells. Truncating the 5⍰ arm (externally of the 4-nt mismatch) from 38 to 20 nt resulted in a modest gain in editing, but further shortening to 10 nt drastically reduced activity (**Figure 1C**). These findings suggest that the duplex formed by the 5’ arm of the gRNA is essential for ADAR1 recruitment and positioning.

Combining both optimizations, we generated a compact 74-nt-long gRNA with reduced 5⍰ and 3⍰ arms, half the size of the original 151-nt LEAPER. This shortened design yielded the highest editing efficiency, as measured by fluorescence restoration (**Figure 1C**), and exhibited reduced off-target editing, as determined by amplicon sequencing (**Figure 1D,E**). Collectively, these results demonstrate that the efficiency and precision of plasmid-encoded ADAR-engagers can be significantly improved through rational design of duplex geometry. Specifically, we identify the length of the duplex region downstream of the 4 nt-mismatch as a key determinant of ADAR1 recruitment. We refer to this optimized design as *Short Precise-Encodable ADAR Recruiting (SPEAR)* gRNAs.

### SPEAR-mediated RNA editing restores T cell activation

To test whether endogenous ADAR1 RNA editing could be leveraged with SPEAR gRNAs to generate functional *editopes* capable of activating antigen-specific T cells, we designed a proof-of-concept experiment based on the well-characterized melanoma antigen MART-1 (Melanoma Antigen Recognized by T cells-1)^33,34^. MART-1 is encoded by the *MLANA* gene and is a clinically relevant immunogenic target in melanoma. Critically for our purposes, the specific T cell receptor (TCR) recognizing the MART-1 epitope (EAAGIGLTV) is well characterized. For this, we used the DMF5 TCR, known for its high avidity against MART-1-expressing cells^35^.

To increase T cell sensitivity and stability of the peptide-MHC complex, we used a MART-1 peptide variant (E**L**AGIGILTV)^36^ (**Figure 2A, S2A**). We then introduced a point mutation in the MART-1 coding sequence to generate the G31S mutant (MART-1^G31S^; ELAGI**<u>S</u>**ILTV), substituting glycine (**<u>G</u>**GC) with serine (**<u>A</u>**GC) at position 31. This mutation disrupted the epitope, rendering it incapable of activating DMF5 T cells (**Figure 2A, B, S2A**). To reverse this effect via RNA editing, we designed a SPEAR gRNA targeting the mutant *MART-1*^*G31S*^ transcript to recruit endogenous ADAR1 and direct A-to-I editing at the first base of the serine codon (**<u>A</u>**GC), converting it to IGC (interpreted by the ribosome as GGC) at the RNA level and restoring the original epitope (Figure 2A, S2A). This system models our broader goal of generating immunogenic *editopes* from non-immunogenic epitopes through precise RNA editing.

**Figure 2.**
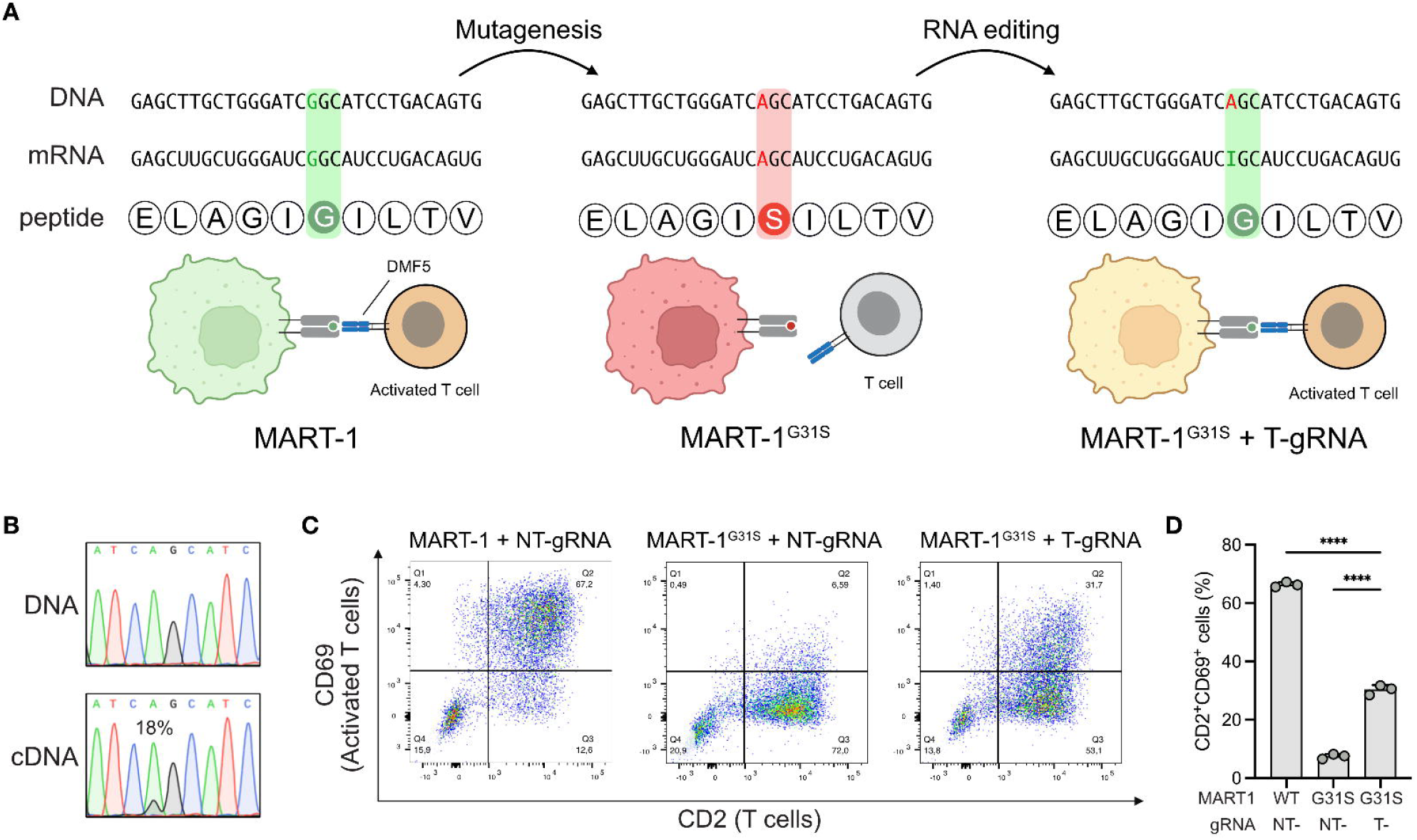
SPEAR gRNAs can generate functional epitopes and activate specific T cells. **(A)** Schematic of the proof-of-principle experiment: HEK293T cells expressing either wild-type (WT) or G31S mutant MART-1 are co-cultured with DMF5-expressing T cells, followed by assessment of T cell activation via flow cytometry. SPEAR gRNAs are used to restore the original MART-1 coding sequence at the RNA level, thereby rescuing antigen-specific T cell activation. (**B**) Representative Sanger sequencing traces at the G31S codon from both genomic DNA and cDNA of HEK293T MART-1 G31S cells expressing the SPEAR gRNA. (C) Representative flow cytometry dot plots showing CD69 expression as a marker of T cell activation under the three experimental conditions. (D) Bar plot showing average T cell activation, measured as the percentage of CD69^+^ CD2^+^ double-positive cells, across three biological replicates for each condition (adjusted p-values: **** < 0.0001, one-way ANOVA with multiple comparisons). Bars represent the mean for three biological replicates; error bars indicate the standard deviation.

To evaluate whether SPEAR-driven ADAR1 editing could restore antigenicity and T cell activation, we engineered HEK293T and TCR-deficient Jurkat cell lines to stably express either *MART-1* ^*WT*^ or *MART-1*^*G31S*^, and the DMF5 TCR, respectively. HEK293T cells were also expressing either a non-targeting (NT) or targeting (T) SPEAR gRNA (Figure S2B, C). Sanger sequencing of

PCR amplicons from genomic DNA and cDNA in *MART-1*^*G31S*^ cells expressing the T-gRNA confirmed A-to-I editing at the intended site, with an editing efficiency of approximately 20% (**Figure 2B**). We then co-cultured the engineered HEK293T and Jurkat cells and assessed T cell activation via flow cytometry using CD2 as a T cell marker and CD69 as an activation marker. As expected, cells expressing *MART-1*^*WT*^ with a NT-gRNA elicited robust T cell activation (∼65% CD2^+^CD69^+^ cells), while *MART-1*^*G31S*^ cells with NT-gRNA showed minimal activation (∼5%). Importantly, co-cultures with *MART-1*^*G31S*^ cells expressing the T-gRNA led to a significant restoration of T cell activation (∼30% CD2^+^CD69^+^ cells) (**Figure 2C, D**).

Despite the “moderate” level of editing, recovery of T cell activation was substantial (∼50%) relative to the wild-type epitope, suggesting that even modest RNA editing rates (<30%) may be sufficient to elicit strong T cell responses and potentially therapeutic immune activation. These results demonstrate that SPEAR gRNAs can efficiently harness endogenous ADAR1 to generate immunogenic peptides capable of activating a specific T cell response. Our findings establish a mechanistic foundation for using programmable RNA editing to engineer *editope* formation in tumor cells.

### Programmable RNA editing drives T cell-mediated tumor rejection in an in vivo preclinical model

Having demonstrated that SPEAR-mediated RNA editing can restore cognate T cell activation *in vitro*, we next evaluated its therapeutic potential *in vivo* using a xenograft mouse model. For this, we employed the human melanoma cell line 624-mel, which endogenously expresses MART-1. We first introduced the same G-to-A mutation described above into the endogenous *MLANA* gene using DNA base editing, resulting in expression of the mutant MART-1^G31S^ from the endogenous locus (**Figure S3A**). These cells were further engineered to stably express either a targeting (T) or a non-targeting (NT) SPEAR gRNA designed to recruit endogenous ADAR1 to MART-1^G31S^ (**Figure S3B-D**). We initially assessed editing efficiency in these modified cell lines and observed ∼10% A-to-I editing at the endogenous target site in the cell line expressing MART-1^G31S^ together with the T-gRNA, while no editing was detected in cells expressing the NT-gRNA (**Figure 3A, B**). To test whether this level of editing was sufficient to induce T cell activation, we performed an *in vitro* T cell-mediated cytotoxicity assay using the xCELLigence system. As expected, 624-mel MART-1^WT^ cells grew normally in the absence of T cells, but the addition of MART-1-specific T cells resulted in a strong antitumoral effect (**Figure 3C**, green lines). In contrast, MART-1^G31S^ cells with no or the control NT-gRNA were unaffected by T cell addition, indicating a lack of recognition by the latter (red and gray lines). Notably, MART-1^G31S^ cells expressing the T-gRNA showed robust killing by T cells, with a cytotoxicity profile similar to MART-1^WT^ cells (**Figure 3C**, orange dotted line). Despite only ∼10% RNA editing at the G31S site, the resulting partial correction was sufficient to generate a functional immunogenic *editope*, again underscoring that even modest editing can restore effective T cell recognition (**Figure 3A-C**).

**Figure 3.**
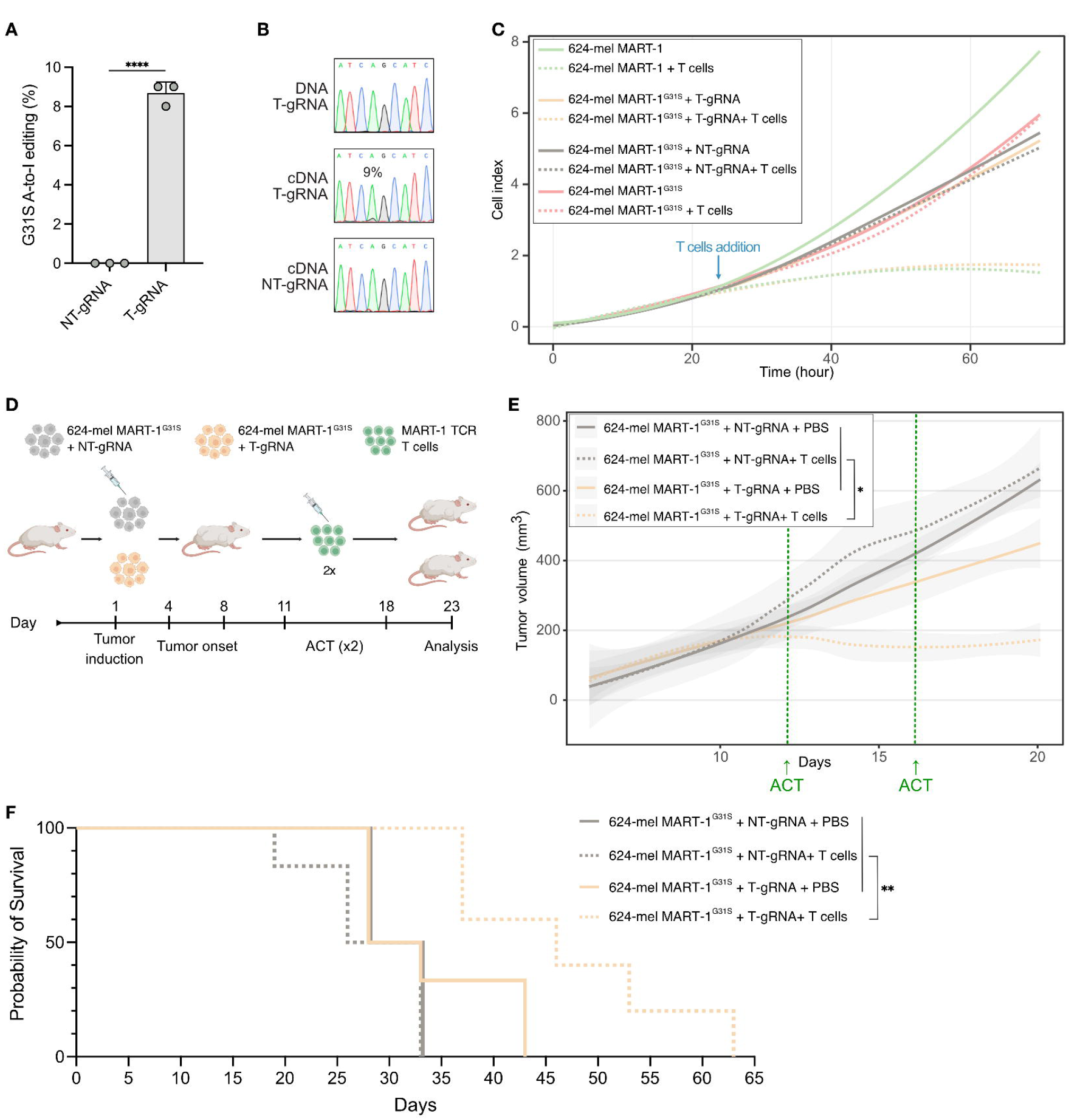
SPEAR gRNA editing induces T cell activation and tumor cell elimination *in vivo*. (**A)** Bar plot showing average A-to-I editing percentages at the MART-1 G31S site, quantified from Sanger sequencing traces using MultiEditR^42^, in 624-mel MART-1^G31S^ cell lines expressing either a non-targeting (NT) or targeting (T) gRNA (p-value: **** < 0.0001, two-tailed unpaired t-test). (**B**) Representative Sanger sequencing traces at the G31S codon from both genomic DNA and cDNA of 624-mel MART-1^G31S^ cells expressing NT- or T-gRNAs. Bars represent the mean for three biological replicates; error bars indicate the standard deviation. (**C**) xCELLigence real-time analysis of the cytotoxicity of MART-1-specific human T cells co-cultured with 624-mel cells expressing MART-1, MART-1^G31S^, or mutant MART-1^G31S^ with NT- or T-gRNAs. For MART-1 or MART-1^G31S^ cells, each curve represents the mean of three technical replicates. For cells expressing NT- or T-gRNAs, each curve represents the mean from three biological replicates, each of those with three technical replicates. T cells were added after 24 hours; the cell index is normalized to the time point of T cell addition. (**D**) Schematic of the *in vivo* experimental setup. (**E**) *In vivo* tumor growth analysis of MART-1^G31S^ cells expressing the T- or NT-gRNA with or without adoptive cell therapy (ACT) using MART-1-specific human-derived T cells. Each curve represents the mean of 6 biological replicates. Ordinary one-way ANOVA was applied: * (P value: 0.0493). (**F**) Kaplan-Meier survival analysis curves of mice injected with MART-1^G31S^ cells expressing the T- or NT-gRNA with or without adoptive cell therapy (ACT) using MART-1-specific human-derived T cells. n = 6, Logrank (Mantel-Cox) test: ** (P value: 0.0055).

We next evaluated this approach *in vivo* using a melanoma xenograft mouse model. Immunodeficient mice were injected subcutaneously with either MART-1^WT^ or MART-1^G31S^ cells to induce tumor formation. After tumors were established, mice received adoptive T cell therapy (ACT) with MART-1 specific T cells (**Figure S3E**). As expected, tumors derived from MART-1^WT^ cells were completely eliminated following two rounds of ACT, while tumors in untreated controls progressed to the experimental endpoint (**Figure S3F**), validating the working parameters of the system. To assess the effect of RNA editing in this context, we repeated the experiment, but now injecting MART-1^G31S^ cells expressing either the NT- or T-SPEAR gRNA into immunodeficient mice (**Figure 3D**). Mice injected with MART-1^G31S^ cells expressing the NT-gRNA failed to respond to ACT, and tumors continued to grow. In contrast, tumors derived from MART-1^G31S^ cells expressing the T-gRNA showed marked regression following ACT (**Figure 3E, S3G**). Furthermore, we also observed prolonged survival for mice carrying tumors derived from MART-1^G31S^ cells expressing the T-gRNA relative to those expressing the NT-gRNA (**Figure 3F**).

Together, these findings demonstrate that SPEAR gRNAs can successfully reprogram tumor cells by recruiting endogenous ADAR1 to generate immunogenic neoepitopes, or *editopes*, enabling precise T cell targeting and tumor rejection.

### A pipeline for the generation of patient-specific editopes across seven cancer types

To extend our proof of concept findings beyond single antigen models and into a broader oncologic context, we developed and applied our neoADARgen algorithm (explained in detail in Methods) to genomic and transcriptomic data from The Cancer Genome Atlas (TCGA). We focused on seven major cancer types: breast cancer (BRCA), skin melanoma (SKCM), lung adenocarcinoma (LUAD), lung squamous cell carcinoma (LUSC), glioblastoma (GBM), ovarian cancer (OV) and liver hepatocellular carcinoma (LIHC). For each of the 3,224 patients in these cohorts, we integrated somatic mutation profiles, HLA haplotype and gene expression data to predict the likelihood that individual mutations could result in neoantigen formation. In total, 808,297 somatic mutations were analyzed, resulting in 19,311 predicted novel neoantigens across all HLA haplotypes. Prior to applying NeoADARgen, patients harbored on average of 4.3 predicted epitopes. Following the algorithm’s application, this number increased to an average of 10.3 epitopes per patient, demonstrating a consistent enhancement in predicted immunogenicity across all cancer types (**Table 1**).

**Table 1.** Summary of NeoADARgen-predicted neoantigens across seven TCGA cancer cohorts. Application of the NeoADARgen algorithm to 3,224 TCGA patients revealed a consistent increase in predicted neoantigens across all analyzed cancer types. The table summarizes cohort sizes, baseline epitope counts, and the number of novel epitopes identified following RNA-editing–based prediction.

| TCGA project | Total number of patients included | Epitopes prior to NeoADARgen algorithm (total / avg. per patient) | Patients with novel epitopes identified | Novel epitopes predicted (total / avg. per patient) | Shared novel epitopes |
| --- | --- | --- | --- | --- | --- |
| BRCA | 965 | 2431/2.5 | 801 | 3675/4.6 | 39 |
| SKCM | 467 | 3089/6.6 | 434 | 4642/10.7 | 41 |
| OV | 289 | 3323/11.5 | 273 | 3190/11.7 | 6 |
| GBM | 147 | 188/1.3 | 107 | 289/2.7 | 4 |
| LIHC | 362 | 702/1.9 | 267 | 1029/3.9 | 4 |
| LUAD | 507 | 1999/3.9 | 474 | 3185/6.7 | 27 |
| LUSC | 487 | 2029/4.2 | 466 | 3301/7.1 | 15 |

For instance, in breast cancer (BRCA), patients initially exhibited an average of 2.5 predicted neoantigens. After applying our algorithm, 801 patients were predicted to generate a total of 3,675 *editopes*, corresponding to an average of about 6.8 neoantigens per patient). These *editopes* could, in principle, be generated by the delivery of multiple SPEAR ADAR-engagers, thereby increasing the number of recognizable antigens and enhancing the likelihood of effective immune targeting in low-TMB tumors.

### Engineering shared neoepitopes between individuals

ADAR-engager based therapies, whether aimed at correcting pathogenic variants or generating personalized *editopes*, are inherently limited in scalability. To broaden the applicability of our editope-generation approach to a wider patient population, we sought to identify a limited set of shared *editopes* (and their corresponding SPEAR-gRNAs) that could cover a large fraction of cancer cases. To this end, we analyzed the 50 most frequent somatic mutations documented in the NCI Genomic Data Commons (GDC) database^37^ across all cancer types, along with the 50 most common HLA alleles represented in the same database. We then applied our algorithm to evaluate each possible mutation-HLA combination and identify potential *editopes* predicted to be strong neoantigens.

For each combination, we first used a peptide-HLA binding prediction tool to calculate the best possible rank before editing^38^. If no high-affinity neoantigen was predicted at a particular site, we ran our algorithm to search for nearby adenosines that could be targeted for editing to generate a neoantigenic peptide. Only combinations predicted to yield a potent neoantigen after editing were retained as viable (**Figure 4**). As an illustrative example, KRAS, one of the most frequently mutated oncogenes, harbors mutations at codon 12, where substitution of glycine is a common driver event in pancreatic, colorectal, and lung cancers. These mutations lead to the constitutive activation of the KRAS protein, promoting oncogenic signaling and uncontrolled proliferation^21^. According to the GDC database, KRAS^G12^ mutations occur in approximately 7% of all cancers.

**Figure 4.**
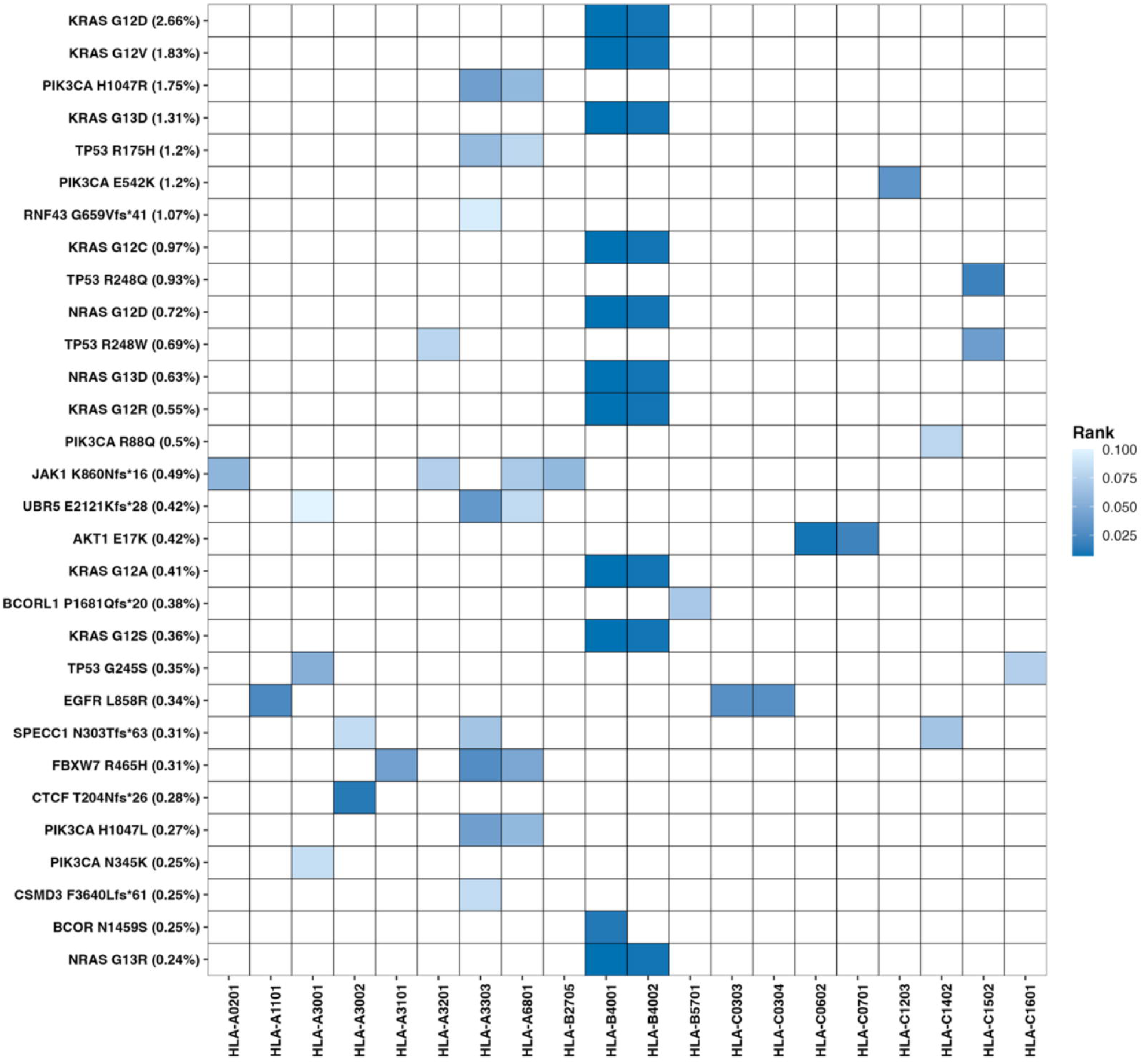
Neoantigens that can be generated next to prevalent hot-spot somatic mutations. A heatmap of the 50 most common somatic mutations across all cancer types and the 50 most prevalent HLA types, illustrating all possible combinations that can generate novel potent neoantigens. Only rows and columns in which a result was detected are shown.

Focusing on this hotspot, our algorithm identified a candidate *editope* for four common HLA types by targeting an adenosine located eleven bases downstream of the mutation site.

The corresponding mRNA sequence is: CUUGUGGUAGUUGGAGCUG**X**UGGCGUAGGC*AGAGUGCCU, where **X** represents the mutation site (as multiple SNVs at this hotspot are known KRAS mutations), and the asterisk (*) denotes the adenosine predicted for editing. For this target, the SPEAR gRNA sequence would be UUGGAUCAUAUUCGUCCACA<u>UUU</u>AGAUUCUGAAUUAGCUGUA

UCGUCAAGGCACUCU<u>C</u>GCCUACGCCAYCAGCU, where the underlined bases are mismatched to the mRNA sequence for editing purposes and **Y** indicates the base complementary to **X**. The resulting neoantigen achieved a predicted rank of 0.01% and a binding affinity of 6.35 nM. Notably, this same *editope* can also be generated for tumors carrying mutations in KRAS^G13^, as well as NRAS^G12^ or NRAS^G13^ due to gene homology, requiring only minor adjustments to the designed gRNA, potentially extending applicability to an additional 2% of cancers. This example highlights how our approach can exploit recurrent hotspot mutations to generate *editopes* across multiple patients. Such shared targets could enable the development of off-the-shelf SPEAR gRNAs for tumors with common mutational profiles, substantially increasing the clinical and pharmaceutical utility of the editope generation strategy.

## Discussion

Enhancing the number and immunogenicity of neoantigens within tumors is a critical step for improving the efficacy of immunotherapy. This is especially relevant for cancer vaccines, where the ability to design potent, tumor-specific neoantigens enables the patient to direct an immune response against these engineered neoantigens. However, the selective generation of neoantigens remains challenging.

Here, we introduce a novel approach for neoantigen engineering based on the recruitment of endogenous ADAR1 via SPEAR gRNAs. This technology enables programmable RNA editing through gRNA delivery and without the need for exogenous enzymes. Furthermore, base editing of RNA rather than DNA enhances safety by reducing the risk of permanent genomic alterations. In our proof of concept, we demonstrated that ADAR1-mediated RNA editing can modulate the immunogenicity of a tumor epitope by altering its RNA sequence while maintaining its translation, processing, and presentation by MHC-I molecules. This RNA-edited epitope, termed *editope*, can be recognized by and activate antigen-specific T cells, inducing T cell-mediated cytotoxicity.

After establishing and demonstrating the feasibility of *editope* generation both *in vitro* and *in vivo*, we developed a computational pipeline capable of designing SPEAR gRNAs tailored to each patient’s HLA genotype but derived from the mutational landscape of a population of tumors. This personalization ensures that edits are introduced exclusively in tumor cells— anchored to somatic mutations—and that the resulting peptides are highly likely to bind MHC-I and be presented as neoantigens.

Applying our computational algorithm to thousands of tumor samples across seven cancer types, we demonstrated that our approach could nearly triple the number of predicted potent neoantigens for the majority of patients. We also showcased several editing candidates derived from prevalent somatic mutations that can serve as entry points for broad clinical applications of oncologic patients.

While our *in silico* predictions provide a compelling roadmap, we acknowledge the complexity of immune recognition. We relied on an established MHC-binding prediction tool that estimates the likelihood of a peptide binding to a specific MHC molecule based on the top percentile scores obtained from random natural peptides^38^. Additionally, we incorporated gene expression levels into our scoring. However, accurate modeling of neoantigen presentation remains challenging due to the involvement of multiple cellular processes, including antigen processing, peptide trimming, proteasomal cleavage, and peptide loading. Currently, modeling these processes computationally remains highly complex^39^. Of course, there are general mechanisms that enable tumor cells to escape immune surveillance, notably the somatic loss of HLA-I heterozygosity, which is a known factor in immune evasion^40^. This is a risk that our platform shares with neoepitope vaccines; however, recent data support substantial efficacy for such vaccines^41^, which theoretically our platform should also share.

A general limitation of ADAR engagers is editing efficiency. Previous studies have reported variable editing levels across targets and delivery formats^19–27^. In our context, high editing efficiency is not necessary: the generation of even a small number of edited transcripts appears to suffice to trigger T cell recognition, as we observed both *in vitro* and *in vivo*. Unlike conventional base editing applications that seek full protein restoration, our goal is to create peptides that are processed and presented as *non-self*, enabling immune activation by a relatively small fraction of edited mRNAs and for this, even low editing efficiency is sufficient.

A final general limitation of targeted ADAR1 recruitment lies in the delivery of the gRNAs to the tissues of interest. This is less applicable to our technology: our focus on *editopes* that are translated from tissue/HLA specific tumour exclusive transcripts, already supports a level of selectivity even to systemic delivery. However as delivery technologies continue to advance, we anticipate that the innovative approach presented here could serve as a valuable therapeutic adjunct for immunotherapy. In all, the approach we present, harnessing programmable RNA editing via SPEAR gRNAs and endogenous ADAR1, offers a safer, precise, and adaptable platform for generating tumor-specific neoantigens or *editopes*. This technology has the potential to strongly enhance immunogenicity in poorly immunogenic (“cold”) tumors and to serve as a powerful complement to current immunotherapeutic strategies, including checkpoint inhibitors and personalized cancer vaccines.

## Acknowledgements and funding

We thank the Flow Cytometry unit of the Imaging and Cytometry Core Facility, German Cancer Research Center (DKFZ), for providing excellent sorting services. We also acknowledge Dr. Florian Grünschläger for his support and guidance in data visualization using R. This work was supported by the Israel Science Foundation grants 2039/20 and 231/21 to E.Y.L., the HI-TRON Kick-Start Seed Funding Program 2021 awarded to RP, and by the Deutsche Forschungsgemeinschaft (DFG, German Research Foundation; SVB TRR319-RMaP) to F.N.P. as well as core funds to F.N.P. by the Helmholtz Association.

## Author contributions

R.P. and F.N.P. conceived the project and supervised the research. R.M.R., N.L., E.R., and E.Y.L. conceived and developed the computational algorithm. R.P. and A.A. conceived, performed, and analyzed targeted RNA editing experiments. V.P. and J.M.L. contributed to the co-culture assays. R.P., B.C., K.S., S.Z., K.K., R.C., M.P., and R.O. conceived, performed, and analyzed experiments in mice. R.P., B.C., and F.N.P. interpreted the overall results of the study. R.P., B.C., and R.N.R. prepared the data visualizations. R.P., B.C., and R.N.R. wrote the manuscript with input from all authors. All authors reviewed and approved the final version of the manuscript.

## Data Availability

The source code used to produce the results and analyses presented in this manuscript are available on a GitHub repository at: https://github.com/landsboy/neoADARgen.git

## Competing interests

R.P. and A.A. are inventors on a patent application related to the design of ADAR-recruiting guide RNAs or ASOs (PCT/EP2024/051033) and R.P., B.C., A.A. and F.N.P. on a separate application covering the use of *editopes*. All other authors declare no competing interests.

## Materials and Methods

### Plasmids

#### pcDNA3.1(-) mCherry-P2A-eGFP^W58X^ puro

This plasmid was generated by inserting a puromycin resistance cassette into the pcDNA3.1(-) mCherry-2A-FLAG-eGFP^W58X^ backbone, which was linearized with BglII. The parental plasmid was a kind gift from Dr. Joshua Rosenthal (University of Chicago) and previously described by *REF*^31^. The puromycin selection cassette was excised from plasmid pLoxPuro^43^ via BamHI digestion and ligated into the linearized vector using the Rapid DNA Ligation kit (Merck, Cat# 11635379001).

#### Plasmids encoding ADAR1-recruiting gRNAs for eGFP ^W58X^

Guide RNA-encoding DNA fragments were generated by inserting single-stranded DNA (ssDNA) oligonucleotides into a pENTR-U6 backbone (a kind gift of Dr. Joshua Rosenthal, University of Chicago), which was linearized via PCR using Q5® High-Fidelity DNA Polymerase (NEB, Cat# M0491) and primers #1 and #2 (see Supplementary Table 1). One or two ssDNA oligos or a dsDNA fragment were inserted into the PCR-amplified vector using NEBuilder® HiFi DNA Assembly Master Mix (NEB, Cat# E2621). All ssDNA oligonucleotides were ordered by Sigma-Aldrich, and the dsDNA fragments by Integrated DNA Technologies (IDT). Nucleotide sequences are provided below, with NEBuilder-compatible overlapping regions shown in lowercase. All plasmids were verified by Sanger sequencing. Detailed plasmid maps are available in the Supplementary Information as text files.

The following gRNA-expressing plasmids were generated for eGFP^W58X^ (**Figure S1B and 1C**):

- **pENTR-U6 GluRv1-16** (eGFP^W58X^) – inserted ssDNA oligo: tatatcttgtggaaaggacgaaacaccGGTGAATAGTATAACAATATGCTAAATGTTGTTATAGTATCCACCGTTGGCCATGGAACAGttttttctagacccagctttcttgtac
- **pENTR-U6 GluRv4-16** (eGFP^W58X^) – inserted ssDNA oligo: tatatcttgtggaaaggacgaaacaccGGTGAAGAGGAGAACAATATGCTAAATGTTGTTCTCGTCTCCACCGTTGGCCATGGAACAGttttttctagacccagctttcttgtac
- pENTR-U6 GluRv9.4-16 (eGFP^W58X^) – inserted ssDNA oligo: tcttgtggaaaggacgaaacacc GGTGTCGAGAAGAGGAGAACAATATGCTAAATGTTGTTCTCGTCTCCTCGACACCGTTGGCCATGGAACAGttttttctagacccagctttctt
- **pENTR-U6 GluRv9.4-38** (eGFP^W58X^) – inserted ssDNA oligos: (1) tcttgtggaaaggacgaaacacc GGTGTCGAGAAGAGGAGAACAATATGCTAAATGTTGTTCTCGTCTCCTCGACACCGTATGTCAGGGTAGTGACAA; (2) acaagaaagctgggtctagaaaaaaCTGTTCCATGGCCAACACTTGTCACTACCCTGACATACGGTGTCGAGGAGACGAGAACAACAT
- **pENTR-U6 LEAPER-70** (eGFP^W58X^) – inserted ssDNA oligo: tcttgtggaaaggacgaaacaccGGATTGCACGCCGTATGTCAGGGTAGTGACAAGTGTTGGCCATGGAACAGGTAGTTTTCCAGTAGTGAAATttttttctagacccagctttctt
- **pENTR-U6 LEAPER-151** (Qu et al 2019) (eGFP^W58X^) – inserted ssDNA oligos: (1) tatcttgtgga aaggacgaaacaccGGTCGTGCTGTTTCATGTGGTCGGGGTACCTGCTGAAGCATTGCACGCCGTATGTCAGGGTAGTGACAAGTGTTGGCCATGGAACAGGTAGTTTT; (2) acaagaaagctgggtctagaaa aaaAAGGAGAGGGCGATGCCACCTATGGCAAGCTGACCCTGAAGTTTATTTGCACTACTGGAAAACTACCTGTTCCATGGCCAACACTTGTCACTACC
- **pENTR-U6 LEAPER-151** (38-4ntMM-33-C-75; Uzonyi et al 2021) (eGFP^W58X^) – inserted ssDNA oligos: (1) tatcttgtggaaaggacgaaacaccGGTCGTGCTGTTTCATGTGGTCGGGGTACCTGCTGAAGCAAACGACGCCGTATGTCAGGGTAGTGACAAGTGTTGGCCATGGAACAGGTAGTTTT; (2) acaagaaagctgggtctagaaaaaaAAGGAGAGGGCGATGCCACCTATGGCAAGCTGACCCTGAAGTTTATTTGCACTACTGGAAAACTACCTGTTCCATGGCCAACACTTGTCACTACC
- **pENTR-U6 38-4ntMM-33-C-31** (sdl 107) (eGFP^W58X^) – inserted ssDNA oligos: (1) tatcttgtgg aaaggacgaaacaccGGTCGTGCTGTTTCATGTGGTCGGGGTACCTGCTGAAGCAAACGACGCCGTATGTCAGGGTAGTGACAAGTGTTGGCCATGGAACAGGTAGTTTT; (2) acaagaaagctgggtctagaa aaaaATTTGCACTACTGGAAAACTACCTGTTCCATGGCCAACACTTGTCACTACC
- **pENTR-U6 38-4ntMM-33-C-16** (sdl 92) (eGFP^W58X^) – inserted ssDNA oligos: (1) tatcttgtgga aaggacgaaacaccGGTCGTGCTGTTTCATGTGGTCGGGGTACCTGCTGAAGCAAACGACGCCGTATGTCAGGGTAGTGACAAGTGTTGGCCATGGAACAGGTAGTTTT; (2) acaagaaagctgggtctagaaa aaaAAACTACCTGTTCCATGGCCAACACTTGTCACTACC
- **pENTR-U6 20-4ntMM-33-C-75** (sdl 133) (eGFP^W58X^) – inserted ssDNA oligos: (1) tatcttgtgg aaaggacgaaacaccGGTCGGGGTACCTGCTGAAGCAAACGACGCCGTATGTCAGGGTAGTGACAAGTGTTGGCCATGGAACAGGTAGTTTT; (2) acaagaaagctgggtctagaaaaaaAAGGAGAGGGCGATGCCACCTATGGCAAGCTGACCCTGAAGTTTATTTGCACTACTGGAAAACTACCTGTTCCATGGCCAACACTTGTCACTACC
- **pENTR-U6 10-4ntMM-33-C-75** (sdl 133) (eGFP^W58X^) – inserted ssDNA oligos: (1) tatcttgtggaaaggacgaaacaccGGTGCTGAAGCAAACGACGCCGTATGTCAGGGTAGTGACAAGTGTTGGCC ATGGAACAGGTAGTTTT; (2) acaagaaagctgggtctagaaaaaaAAGGAGAGGGCGATGCCACCTATGGCAAGCTGACCCTGAAGTTTATTTGCACTACTGGAAAACTACCTGTTCCATGGCCAACACTTGTCACTACC
- **pENTR-U6 20-4ntMM-33-C-16** (sdl 74, SPEAR) (eGFP^W58X^) – inserted ssDNA oligo: cttgtgga aaggacgaaacaccGGTCGGGGTACCTGCTGAAGCAAACGACGCCGTATGTCAGGGTAGTGACAAGTGTTGGCCATGGAACAGGTAGTTTttttttctagacccagctttct

#### Plasmids for *in vitro* T cell activation assay

The plasmids used for the proof-of-concept T cell activation assay (**Figure 2, S2**) contain a dual-expression cassette structure: a hU6-driven gRNA expression cassette, followed by a hPGK1-driven cassette encoding a codon optimized MART-1 (coMART-1), eBFP, and the blasticidin resistance gene (BSD), separated by two different self-cleaving 2A peptides^30^ (**Figure S2B**). This design enables both antibiotic selection and monitoring of MART-1 expression via eBFP fluorescence. We began by ordering a custom plasmid backbone from VectorBuilder containing the hU6-gRNA and hPGK1-BSD expression cassettes (VectorBuilder ID: VB210525-1302qyy; this ID can be used to retrieve detailed information about the vector on vectorbuilder.com). The vector was linearized using SpeI-HF (NEB, Cat# R3133) and BsrGI-HF (NEB, Cat# R3575). The individual components—hU6-gRNA cassette, hPGK-MART-1^WT^- or MART-1^G31S^-T2A, and eBFP-P2A-BSD—were amplified separately by PCR using Q5® High-Fidelity DNA Polymerase (NEB, Cat# M0491) from existing plasmids, and assembled into the linearized backbone using NEBuilder® HiFi DNA Assembly Master Mix (NEB, Cat# E2621). The resulting plasmids express either a non-targeting (NT) or targeting (T) SPEAR gRNA as follows:

- **NT-SPEAR gRNA** (control): TCGGGGTACCTGCTGAAGCAAACGACGCCGTATGTCAGGGTAGTGACAAGTGTTGGCCATGGAACAGGTAGTTT
- **T-SPEAR gRNA** (coMART-1^G31S^): TCCGGCAGTACCAGCAGCCGTAGTGCAGCAGCACGCCTAGGATCACTGTCAGGAGGCCGATCCCAGCAAGCTCC

#### Plasmids for *in vivo* T cell activation and tumor elimination assay

The plasmids used in the *in vivo* experiments (**Figure 3, S3**) were also designed with a dual-expression cassette structure comprising a hU6-driven gRNA expression cassette, followed by an hPGK1-driven cassette encoding eBFP and the blasticidin resistance gene (BSD), separated by a self-cleaving 2A peptide (**Figure S3B**). This configuration enables antibiotic selection and allows monitoring of plasmid integration through eBFP fluorescence. To enable stable genomic integration, the dual-expression cassette was cloned into a pT vector containing the necessary inverted terminal repeats (ITRs) for Sleeping Beauty transposon system-mediated integration^44^. The vector was linearized using NsiI-HF (NEB, Cat# R3127) and XhoI (NEB, Cat# R0146). The individual elements—hU6-gRNA cassette and hPGK-eBFP-P2A-BSD—were amplified separately by PCR using Q5® High-Fidelity DNA Polymerase (NEB, Cat# M0491) from existing plasmids, and assembled into the linearized backbone using NEBuilder® HiFi DNA Assembly Master Mix (NEB, Cat# E2621). The resulting plasmids express either a non-targeting (NT) or targeting (T) SPEAR gRNA as follows:

- **NT-SPEAR gRNA** (control): TCGGGGTACCTGCTGAAGCAAACGACGCCGTATGTCAGGGTAGTGACAAGTGTTGGCCATGGAACAGGTAGTTT
- **T-SPEAR gRNA** (MART-1^G31S^): TTCTACAATACCAACAGCCGTACTGCAGTAAGACTCCCAGGATCACTGTCAGGATGCCGATCCCAGCGGCCTCT

For stable integration, these plasmids were co-transfected with pPGK-SB13, a Sleeping Beauty transposase expression plasmid driven by a PGK1 promoter. This plasmid was a gift from Masashi Narita (Addgene plasmid #236078; http://n2t.net/addgene:236078; RRID: Addgene_236078)^45^.

#### Plasmids for genome base editing and single guide RNA (sgRNA) expression

The plasmid AALN-BE4max was a gift from David Liu (Addgene plasmid #138161; http://n2t.net/addgene:138161; RRID: Addgene_138161) and encodes a modified cytosine base editor consisting of rAPOBEC1(AALN)-nSpCas9-UGI-UGI, in which APOBEC1 has been engineered to reduce off-target activity^46^. To facilitate the identification of transfected cells, we inserted an eGFP reporter into this plasmid. Specifically, plasmid AALN-BE4max was digested with AgeI-HF (NEB, Cat# R3552) to linearize it between the UGI-SV40_NLS and 6xHis tag regions. The insert, encoding P2A-EGFP-SV40_NLS-6xHis, was PCR-amplified from pCMV-T7-ABEmax(7.10)-SpCas9-NG-P2A-EGFP (Addgene plasmid #140005; gift from Benjamin Kleinstiver; http://n2t.net/addgene:140005; RRID: Addgene_140005)^47^ using Q5® High-Fidelity DNA Polymerase (NEB, Cat# M0491) and primers #3 and #4 (see Supplementary Table 1). The insert was assembled into the linearized backbone using NEBuilder® HiFi DNA Assembly Master Mix (NEB, Cat# E5520).

For sgRNA expression, we used the vector pFYF1320 EGFP Site#1, a gift from Keith Joung (Addgene plasmid #47511; http://n2t.net/addgene:47511; RRID: Addgene_47511)^48^, which originally encoded a sgRNA targeting EGFP. This sequence was replaced with a sgRNA targeting the *MART-1* gene as follows. The backbone was amplified using Q5® PCR to exclude the EGFP-targeting region using primers #5 and #6 (see Supplementary Table 1). A ssDNA oligo containing the protospacer for the *MART-1*-targeting sgRNA (atatatcttgtggaaag gacgaaacaccggGATGCCGATCCCAGCGGCCCgttttagagctagaaatagcaagtt) was assembled into the linearized backbone using NEBuilder® HiFi DNA Assembly Master Mix (NEB, Cat# E5520).

#### DNA oligonucleotides

All primers used in this study can be found in (Supplementary Table S1) and were designed using Primer-BLAST^49^, AmplifX 2.1.1^50^, or ApE^51^.

#### Cell culture and cell lines generation

HEK293T cells (ATCC, Cat# CRL-3216, RRID: CVCL_0063) were cultured at 37°C in a humidified incubator with 5% CO_2_ in high-glucose DMEM (Sigma-Aldrich, Cat# D6429), supplemented with 5% FBS (PAN Biotech, Cat# P40-37100) and 1% penicillin/streptomycin (Sigma-Aldrich, Cat#P4333).

The stable dual-reporter cell line expressing mCherry-P2A-eGFP W58X-puro was generated by transfecting the plasmid into HEK293T cells using Lipofectamine 2000 (ThermoFisher, Cat# 11668019) according to the manufacturer’s protocol. 48 h after transfection, cells were plated by limiting dilution into 96-well plates in medium supplemented with puromycin (1.5 µg/ml). Clonal populations were identified by microscopic inspection, and single clones were screened via flow cytometry analysis to select a single clone stably expressing the dual fluorescence reporter plasmid with no detectable EGFP background.

HEK293T cells stably expressing the constructs for the *in vitro* T cell activation assay (**Figure 2, S2**), which included hU6-gRNA (T- or NT-) and hPGK1-coMART-1^WT or G31S^-2A-eBFP-2A-BSD were generated similarly. Cells were transfected using Lipofectamine 2000, expanded for 48 hours, and then selected with blasticidin (15 µg/ml). As soon as non-transfected cells died, BFP^+^ cells were sorted by flow cytometry. The expression of MART-1^WT or G31S^ and the presence of SPEAR-mediated editing were confirmed by RT-PCR and Sanger sequencing.

Jurkat cells expressing the DMF5 TCR were cultured at 37°C and 5% CO_2_ in RPMI 1640 (Sigma-Aldrich, Cat# R8758) supplemented with 10% FBS (PAN Biotech, Cat# P40-37100), 2 mM L-glutamine (Gibco, Cat# 25030-024) and 1X penicillin/streptomycin (Sigma-Aldrich, Cat#P4333). Jurkat-DMF5 cells were generated as previously described^52^ using a lentivirally-encoded, single- OF TCR sequence separated by a furin-T2A cleavage sequence, followed by 12 days of puromycin selection for the transgenic TCR.

624-mel cells (RRID:CVCL_8054) were cultured at 37°C and 5% CO_2_ in high-glucose DMEM (Sigma-Aldrich, Cat# D6429) supplemented with 5% FBS (PAN Biotech, Cat# P40-37100) and 1% penicillin/streptomycin (Sigma-Aldrich, Cat#P4333).

The 624-mel MART-1^G31S^ cell line was generated via DNA base editing (see below). For *in vivo* experiments (**Figure 3, S3**), 624-mel MART-1^G31S^ cells were transfected with plasmids expressing either a T- or NT-gRNA together with pPGK-SB13 (Sleeping Beauty transposase) in a 2:1 ratio, using Lipofectamine 2000 (ThermoFisher, Cat# 11668019). After 48 hours, cells were expanded, and selected with blasticidin (5 µg/ml). Once selection was complete, BFP^+^ cells were sorted by flow cytometry. Successful expression of MART-1^G31S^ and gRNA-dependent RNA editing were confirmed by RT-PCR and Sanger sequencing.

All cell lines were authenticated using Multiplex Cell Authentication by Multiplexion (Heidelberg, Germany) as described recently^53^. Additionally, the purity of cell lines was validated using the Multiplex cell Contamination Test by Multiplexion (Heidelberg, Germany) as described recently^54^. No Mycoplasma, SMRV or interspecies contamination was detected.

#### Genome base editing with a cytosine base editor (CBE)

10^5^ 624-mel cells were seeded in 500 μl complete growth medium (DMEM, FBS 10%, P/S 1%) in a flat-bottom 24-well plate. After 18-24 hours, the complete growth medium was replaced with 250 μl antibiotic-free reduced serum medium (OptiMEM, gibco, Cat #31985062). Cells were transfected with a mixture of 2 μl lipofectamine 2000, 500 ng sgRNA vector and 1 μg base editor vector diluted in 100 μl OptiMEM following the manufacturer’s instructions. After 6 hours, the transfection medium was replaced with 500 μl complete growth medium. After 72 hours, the medium was removed, and cells were washed with 250 μl PBS. To detach the cells, 100 μl trypsin were added, and cells were incubated for 1 minute at 37°C. Trypsin was inactivated by adding 400 μl pre-warmed complete growth medium. Detached cells were filtered and transferred to a 5 ml round bottom polystyrene test tube with cell strainer snap cap (Corning, Cat #352235). GFP+ cells were single-cell sorted into flat-bottom 96-well plates. Single-cell clones were incubated for at least 2 weeks at 37°C until they were ready to be screened. When ready, medium was removed, and clones were washed once with 100 µl PBS. Cells were detached by adding 20 µl trypsin and resuspended in 200 µl complete growth medium. 100 µl of each clone were transferred into a new flat-bottom 96-well plate already prepared with 100 µl medium per well to keep and further expand the selected clones. The remaining 100 µl were transferred into another flat-bottom 96-well plate already prepared with 100 µl medium per well for screening the next day. DNA was extracted with QuickExtract™ DNA Extraction Kit (Lucigen, Cat #QE0905T). Briefly, medium was removed from the screening plate and cells were washed once with 100 µl PBS. 100 µl Quick-Extract DNA solution were added, cells were detached by thoroughly pipetting up and down and transferred into a 96-well PCR plate (or PCR strips). Cells were incubated at 65°C for 15 min in a thermal cycler, vortexed for 15 sec, and incubated at 98°C for 5 min a thermal cycler. The extracted DNA was used for PCR and then stored at −80°C. PCR was performed on the extracted DNA with Q5 High-Fidelity DNA Polymerase from NEB (#M0491S) according to the manufacturer’s instructions and the following conditions. 5-20 ng of DNA (2.5 μl) were added to 31 μl of RNase-free water, 10 μl of Q5 Buffer, 1 μl of dNTP mix (10 mM each), 2.5 μl of each primer (10 μM) and 0.5 μl of Q5 polymerase in a final reaction volume of 50 μl. *MART-1* amplification was obtained using primers #7 and #8 (Supplementary Table 1). The PCR program was run as follows: denaturation at 98°C for 30 sec, followed by 35 cycles of denaturation at 98°C for 10 sec, annealing at 65°C for 15 sec and elongation at 72°C for 10 sec. In the end, a final elongation step at 72°C was run for 2 min. The correct size of the PCR products (245 bp) was checked on a 1.5% agarose gel. 5 μl of each product diluted with 10 μl nuclease-free water was sent for Sanger sequencing to Eurofins Genomics Germany GmbH.

#### Co-culture and related flow cytometry analysis

In the co-culture experiment using HEK293T overexpressing either MART-1 or MART-1^G31S^ (**Figure 2, S2**), we used the following antibodies: PE anti-human CD69 (BioLegend, Cat# 310906) and APC anti-human CD2 (BioLegend, Cat# 300214). We additionally assessed viability of cells using 7AAD staining (Invitrogen, Cat# A1310). 2.5 x 10^4^ HEK293T stably expressing either MART-1 or MART-1^G31S^ and the SPEAR gRNAs (targeting or not-targeting MART-1^G31S^) were seeded in a 96-wells flat bottom plate in 40 μl of supplemented DMEM. 24 h post seeding, 1 x 10^5^ DMF5^+^ Jurkat cells in 35 μl of supplemented RPMI were added to the same wells for a total volume of 75 μl per well. The cells were maintained in co-culture for 16 h at 37°C and 5% CO_2_. Samples were washed twice in FACS buffer (2 mM EDTA and 2% FBS in PBS) and cells were stained using 50 μl of FACS buffer containing the antibodies diluted 1:200. After 20 minutes of incubation at 4°C, samples were washed twice and resuspended in a final volume of 200 μl of FACS buffer. Flow cytometer acquisition was done using a BD Canto II (BD Biosciences), and data were analyzed with FlowJo.

#### Adenosine-to-inosine (A-to-I) editing sites quantification

For A-to-I editing site quantification, RNA was extracted using the RNeasy Mini Kit (Qiagen, Cat# 74106) and treated with DNase (Invitrogen, Cat# AM1907). One-step RT-PCRs were performed using the One-step RT-PCR kit (Qiagen, Cat# 210212) using primers #9-10 (for coMART-1), #11-12 (for endogenous MART-1) or #13-14 for eGFP^W58X^. Additionally, when needed the same PCR was performed on genomic DNA (extracted using the All Prep DNA/RNA Kit, Qiagen, Cat# 80204) with Q5 High-Fidelity DNA Polymerase (NEB, Cat# M0491) as a no-editing control. All the PCR products were purified (Macherey-Nagel, Cat# 740609) and analyzed by Sanger sequencing. Quantification of editing was performed directly from the Sanger traces using MultiEditR^42^.

#### On-target and bystander off-target editing analysis via amplicon-seq

To evaluate in detail on-target and bystander off-target editing on the eGFP W58X reporter, a second PCR with primers containing adaptor sequences (#15-16, Supplementary Table 1) for MiSeq amplicon sequencing (Eurofins Genomics, GATC services, Germany) was performed using Q5® High-Fidelity DNA Polymerase (NEB, Cat# M0491). Before performing the second PCR, the amplicons from three independent replicates were mixed in a 1:1:1 ratio. Analysis of the amplicon-seq data was performed using CRISPResso2^55^.

#### xCELLigence cytotoxicity assay

MART-1 ELAGIGILTV/A2 TCR (DMF5) was transduced into human PBMC. DMF5 positive cells were expanded in the presence of PBMCs feeder cell loaded with MART-1 peptide (ProImmune) and IL2 (70 IU/ml) for 14 days. >90% of T cells were DMF5 positive after the procedure. Cell-mediated cytotoxicity was analyzed in an impedance-based label-free cytotoxicity assay (xCELLigence) system^56^. 2 x 10^4^ 624-mel cells (each cell line in 3 technical replicates; 3 biological replicates of the MART-1^G31S^ cells expressing either the NT- or T-SPEAR gRNA were compared) were seeded in 50 μl of complete growth medium (DMEM FBS 10% P/S 1%) in a 96-Well E-plate (E-Plate 96 PET, Agilent). The attachment of the cells changes the electrical impedance of these electrodes, which is monitored as an increase of the dimensionless “cell index”. After 24 hours, 4 x 10^4^ MART-1 specific T cells were added in 50 μl of complete growth medium in half of the samples (condition “+ T cells”), while the other half only received 50 μl of fresh complete growth medium. Lysis of the tumor cells by the T cells causes their detachment from the electrodes, which changes the impedance of the wells, resulting in a decrease in the cell index. The tumor cell growth was continuously monitored from the seeding of the tumor cells until 48 hours after addition of the T cells. The co-culture of tumor cells and T cells was incubated for 48 h at 37 °C5% CO_2_ and 95% humidity.

#### Mice

Immunodeficient NXG mice came from the Janvier Laboratory and were bred at the DKFZ animal facility. 10- to 16-week-old mice were assigned to age-matched experimental groups. Mice were housed under Specific and Opportunistic Pathogen Free (SOPF) conditions and a 12-hour day/night cycle. Animal procedures were performed in accordance with all relevant ethical regulations for animal testing and research, and were approved by the governmental authorities (Regional Administrative Authority Karlsruhe, Germany; permit number 35-9185.81/G-11/22).

#### In vivo tumor growth and Adoptive T cell Therapy (ATC)

On day 1, 500.000 melanoma cells in 100 μl were mixed in a 1:1 volume with 100 μl Matrigel Basement Membrane Matrix (Corning, Cat# 356234) and kept strictly on ice till injection in mice. The cell-Matrigel mix was subcutaneously injected on the right flank of NXG mice. Each cell line was injected into 12 mice. Tumor size was measured every other day starting on day 6. On days 12 and 19, 5 x 10^6^ MART-1 specific T cells in 50 μl PBS were administered intratumorally to half of the mice. The other half received PBS as negative control. On day 23, mice were sacrificed, tumors were isolated (if any) and analyzed via flow cytometry and ELISA to search for markers of T cell activation.

#### Statistical analysis and data visualization

Data were analyzed and plotted using GraphPad Prism (version 10.5.0). Specific information about data presentation is provided in each figure caption throughout the manuscript. Multiple illustrations were created with BioRender.

#### Algorithm

Here we present the NeoADARgen – an algorithm that designs a patient-specific gRNA tailored to the individual’s unique tumor genetic characteristics. Owing to its precise design, using this gRNA as the main component of an Endogenous-ADAR will induce A-to-I(G) edits at the tumor RNA level upon delivery to the tissue. The gRNA interacts exclusively with tumor cells, as it is designed to base-pair with a region containing somatic mutations that are absent in normal cells. Our algorithm identifies the optimal points for editing, which in turn alter the translated peptide within the cancer cell, creating a novel potent neoantigen that takes into account the individual’s HLA typing. We utilize a list of somatic mutations sequenced from the tumor and the patient’s HLA typing as basic inputs. For increased accuracy, incorporating tumor gene expression levels is also feasible.

##### Step 1: Ensuring binding to tumor cells while excluding normal cells

To generate neoantigens in tumor-cells but not in normal-cells, the initial step involves targeting a somatic mutation within the cancerous cell. This mutation can manifest in any possible variant type, as long as it modifies the RNA sequence in comparison to the wildtype. The preliminary gRNA is programmed to bind by base pairing to a sequence comprising the mutation along with the 20 nucleotides upstream and the 20 nucleotides downstream, totaling in a sequence of ∼40 nucleotides. Due to the presence of the somatic mutation at the core of the sequence, the gRNA binds to the cancerous cells but not to the normal cells upon approaching the tissue, as normal cells typically lack the tumor somatic variants. This step guarantees binding specificity exclusively to the tumor cells (**Figure S4**).

##### Step2: Identifying the optimal adenosines in the region for editing

The following step involves scanning the gRNA sequence for adenosines which upon conversion to guanosines will modify the peptide in a manner predicted to generate a potent neoantigen. For this purpose, we evaluate each adenosine within the sequence to determine the effect of editing it to guanosine on the resulting protein. Given the natural ADAR’s ability to edit multiple adenosines in clusters, we also explore the impact of editing two adenosines at a time on the resulting protein. Note that adenosines preceded by guanosines are not considered, as they do not meet the criteria for the ADAR motif, which requires the absence of a guanosine 5’ to the edited adenosine. (While this constraint can be overcome by manipulating the gRNA to include an additional mismatch 5’ to the edited adenosine, we prefer not to do so in order to maintain ideal complementarity to the cancerous sequence). Additionally, the examination excludes the first and last nucleotides of the sequence.

To identify the optimal alteration capable of generating a neoantigen, we analyzed all possible 9-mer sub-sequences derived from the gRNA sequence, where one or two adenosines are substituted with guanosines (**Figure S5**). To assess the effect on the resultant peptide, we employed the external tool NetMHCpan4.1^38^ which utilizes artificial neural networks to predict peptide binding for a given sequence by a given HLA. We submitted each combination of altered sub-sequences along with each HLA class I type of the patient to the prediction tool, examining the output parameters for each combination. By default, thresholds of %Rank < 0.5% and %Rank < 2% are used to identify strong MHC binders (SBs) and weak MHC binders (WBs) for class I, respectively^38^. In our analysis, we used even more conservative parameters: %Rank < 0.1% and affinity < 50 as indicators for SB. Additionally, if gene expression data were available, we set a threshold of TPM above 15 as an additional mandatory criterion for robustness. This ensures adequate transcription of the gene for RNA editors to function effectively. If more than one combination met these criteria, we prioritized the sub-sequence with the lowest detected rank.

##### Step3: Presenting the best results

At the end, the algorithm presents all potential gRNA sequences, marking the optimal adenosines to edit with an asterisk (*). These gRNAs are the optimal sequences tailored for a given tumor sample, to be utilized with Endogenous-ADAR base editors. Upon delivery to the tissue, they will bind to a region encompassing a somatic mutation expressed in this tumor to ensure cancer cell specificity and induce single or double A-to-I editing leading to precise peptide alterations. These alterations are predicted to generate neoantigens according to the patient’s HLA typing.

#### Cancer databases

The computational results presented are based on cancer data extracted from the Tumor Cancer Genome Atlas (TCGA). To apply our algorithm to a variety of cancer samples, we utilized the Genomic Data Commons (GDC) data portal [https://portal.gdc.cancer.gov/]. We downloaded the data for seven cancer TCGA projects: BRCA, SKCM, OV, GBM, LIHC, LUAD, and LUSC. The first five were downloaded in September 2023 and the last two in January 2024. For each patient, we used the DNA sample as sequenced from the primary tumor or from the metastatic tumor if the primary sample was not available. In case the DNA file was not available, we used the whole genome sequence file (Supplementary Table 2). For each patient, we also used the gene expression quantification file as sequenced from its tumor. Lastly, we used The Cancer Immunome Atlas (TCIA) data portal [https://tcia.at/home] to extract the HLA typing of each individual as reported by its unique TCGA ID. Only individuals with access to all data were included in our analysis.

To apply our algorithm to the most common cancer hotspots, we extracted a list of the most common somatic mutations and their reported prevalence across all cancer types as presented in GDC. We also downloaded all TCGA samples available in TCIA to aggregate the most common HLA types among them. Both lists were downloaded in September 2023. For this analysis gene expression could not be considered by the algorithm, as the calculation was not done on an individual basis.

## Notes

### Summary of Updates

The manuscript is completely revised. While the general concept is the same, all figures and text are new.

