## Supplementary Information for "Employing RNA editing to engineer personalized tumor-specific neoantigens (editopes)"

Pecori *et al.*

**Inventory:**

1. **Supplementary Information file**
   a. Supplementary Tables	page	2
   b. Supplementary Figures	pages 3-10
   c. Plasmid maps (provided as text)	pages 11-69
   d. References	page	69
a. **Supplementary Tables**

| **#** | **Name** | **Sequence** | **Purpose** | **Figure** |
| --- | --- | --- | --- | --- |
| 1 | pENTR_plasm_F | TTTTTTCTAGACCCAGCTTTCTTGTA | To open via PCR pENTR-U6 plasmid |  |
| 2 | pENTR_plasm_R | GGTGTTTCGTCCTTTCCACA | To open via PCR pENTR-U6 plasmid |  |
| 3 | p2a-eGFP-NLS-6xHis_F | agaagaagaggaaagtctaaccggtGCTACTAACTTCAGCCTG | To amplify p2a-GFP-SV40 NLS-6xHis from Addgene plasmid #140005 |  |
| 4 | p2a-eGFP-NLS-6xHis_R | ATGGTGATGGTGATGATG | To amplify p2a-GFP-SV40 NLS-6xHis from Addgene plasmid #140005 |  |
| 5 | pFYF1320_F | GTTTTAGAGCTAGAAATAGC | To open via PCR pFYF1320 plasmid |  |
| 6 | pFYF1320_R | GGTGTTTCGTCCTTTCCACA | To open via PCR pFYF1320 plasmid |  |
| 7 | gDNA_MART1_F | CTGCTCTGGGTCGTCAAAGT | To screen for MART1^G31S^ positive clones upon DNA base editing (624-mel). |  |
| 8 | gDNA_MART1_R | TGGACTTGGAGGGATTTCAG | To screen for MART1^G31S^ positive clones upon DNA base editing (624-mel). |  |
| 9 | coMART1_F | ATGCCCAGGGAGGACGCCCA | To quantify editing at the G31S site of the codon optimized MART1 (HEK293T). | 2 |
| 10 | coMART1_R | CTCCACCATACAGCCCCGGT | To quantify editing at the G31S site of the codon optimized MART1 (HEK293T). | 2 |
| 11 | RT-PCR_MART1_F | ACGGCCACTCTTACACCACG | To quantify editing at the G31S site of the endogenous MART1 (624-mel). | 3, S3 |
| 12 | RT-PCR_MART1_R | TGGAGCATTGGGAACCACAGG | To quantify editing at the G31S site of the endogenous MART1 (624-mel). | 3, S3 |
| 13 | RT-PCR_eGFP_F | AACTTCAGCCTGCTCAAACAAGCC | To quantify editing at the W58X site of the mCherry-2A-eGFP^W58X^ reporter. | 1, S1 |
| 14 | RT-PCR_eGFP_R | CAGCCCTGGTCTTGTAGTTG | To quantify editing at the W58X site of the mCherry-2A-eGFP^W58X^ reporter. | 1, S1 |
| 15 | MiSeq_eGFP_F | acactctttccctacacgacgctcttccgatctAACTTCAGCCTGCTCAAACAAGCC | MiSeq amplicon sequencing on eGFP W58X. | 1, S1 |
| 16 | MiSeq_eGFP_R | gactggagttcagacgtgtgctcttccgatctCAGCCCTGGTCTTGTAGTTG | MiSeq amplicon sequencing on eGFP W58X. | 1, S1 |

**Supplementary Table 1 |** Primers used in this study. Uppercase letters represent the specific sequence complementary to the specific target to amplify.

| **TCGA project** | **Individuals in this project** | **Individuals included in the analysis** | **DNA files used** | **Whole genome files used** |
| --- | --- | --- | --- | --- |
| BRCA | 969 | 965 | 755 | 211 |
| SKCM | 470 | 467  (primary:102 metastatic:365) | 467 | - |
| OV | 419 | 289 | 26 | 263 |
| GBM | 374 | 147 | 147 | - |
| LIHC | 369 | 362 | 362 | - |
| LUAD | 559 | 507 | 505 | 2 |
| LUSC | 490 | 487 | 481 | 6 |

**Supplementary Table 2 |** The seven TCGA projects utilized for the computational analysis in this study.

b. **Supplementary Figures**

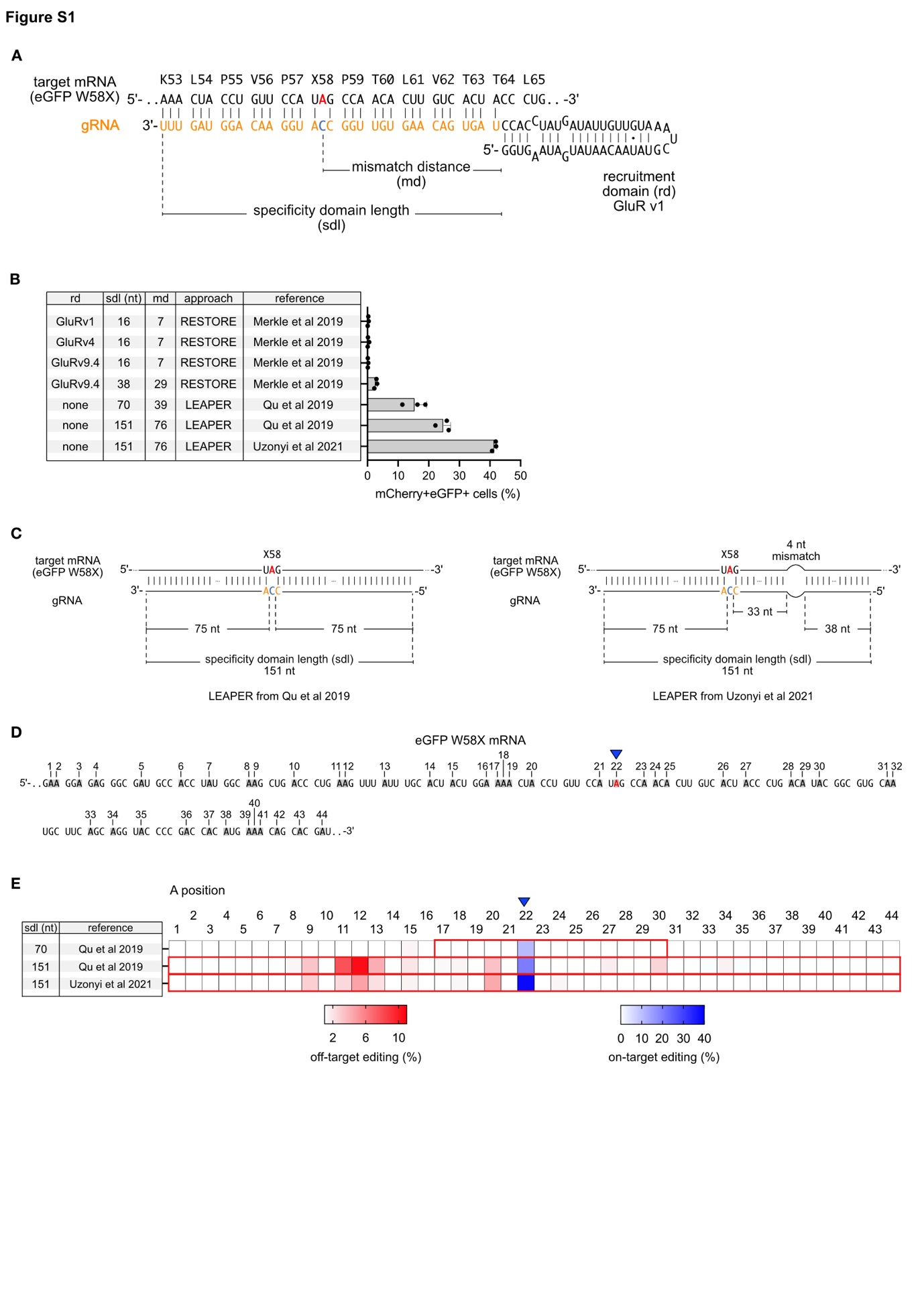

**Figure S1. Optimization of Short Precise Encodable ADAR Recruiting (SPEAR) gRNAs.** (**A**) Schematic of the eGFP W58X target mRNA and the corresponding targeting gRNAs used in the experiment shown in panel **B**. (**B**) Flow cytometry analysis of mCherry and eGFP double-positive cells, presented as a bar plot for the tested gRNAs. Bars represent the mean for three biological replicates; error bars indicate the standard deviation. (**C**) Diagram illustrating the difference between LEAPER gRNAs without (left, *REF*^1^) or with a 4-nucleotide (nt) mismatch (right, *REF*^2^). (**D**) Schematic of the sequencing window where A-to-I edits were assessed, as shown in panel **E**. (**E**) Heatmap of A-to-I editing from NGS amplicon sequencing, showing on-target editing (in blue) and off-target editing (in red) for three different LEAPER gRNAs. The region targeted by each gRNA is indicated with a red rectangle.

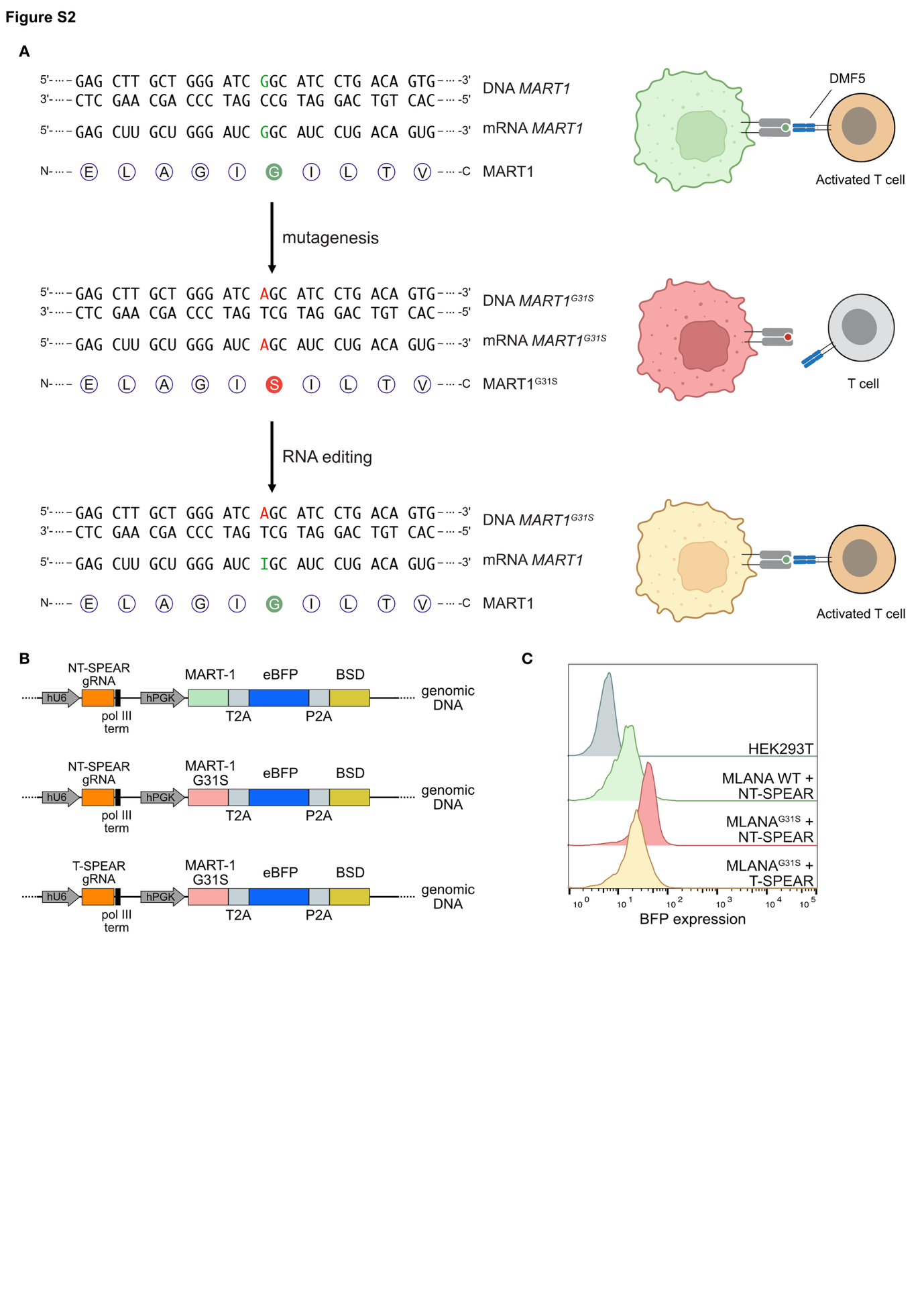

**Figure S2. *In vitro* T cell activation using SPEAR gRNAs.** (**A**) Diagram illustrating the molecular setup (left) and the predicted outcome of the cellular co-culture (right) corresponding to the experiment presented in Figure 2. (**B**) Schematic of the constructs used to generate HEK293T cells expressing either wild-type (WT) or G31S mutant MART-1, along with either a non-targeting (NT) or targeting (T) SPEAR gRNA. eBFP was used as a reporter for MART-1 (WT or G31S) expression, and a blasticidin resistance gene (BSD) was used for selecting transfected cells. (**C**) Flow cytometry analysis of BFP expression for the different cell lines.

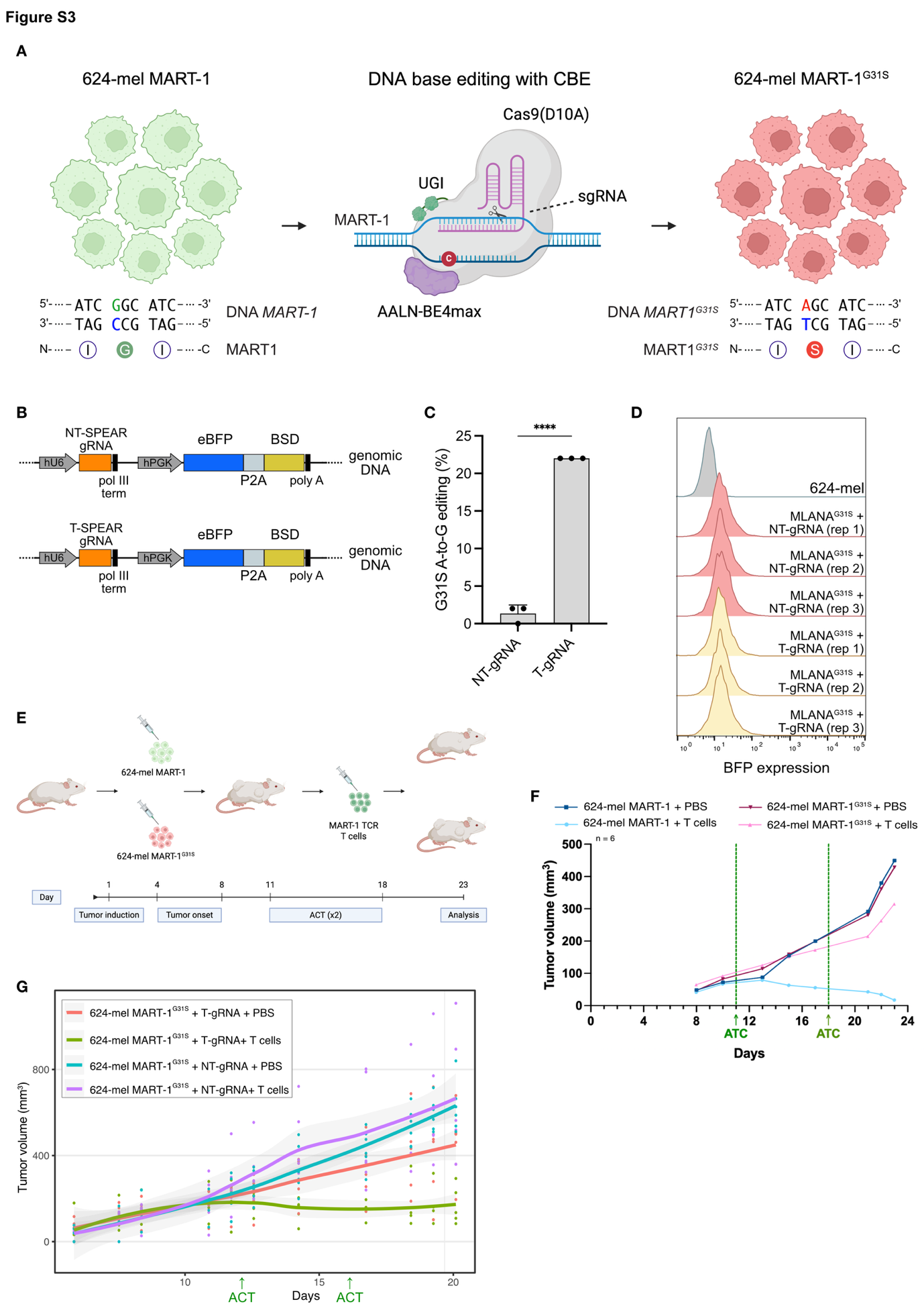

**Figure S3. Generation of 624-mel mutant and edited cell lines for evaluating T cell activation and tumor cell elimination *in vivo*.** (**A**) Schematic of the procedure used to generate 624-mel cell clones endogenously expressing MART-1^G31S^. The cytosine base editor AALN-BE4max^3^ was used to target the cytosine (C highlighted in blue) on the template strand of DNA, opposite the first guanine (G highlighted in green) in the GGC codon encoding glycine (G) at position 31 of the *MART-1* gene. C-to-U base editing converted the codon to AGC, resulting in a glycine-to-serine substitution (G31S). (**B**) Schematic of the constructs used to generate 624-mel MART-1^G31S^ cells expressing either a non-targeting (NT) or targeting (T) SPEAR gRNA. eBFP was used as a reporter for plasmid integration, and a blasticidin resistance gene (BSD) was used for selection. (**C**) Bar plot showing average A-to-I editing percentages at the MART-1 G31S site, quantified from Sanger sequencing traces using MultiEditR^4^ in 624-mel MART-1^G31S^ cells transiently transfected with the plasmids shown in **B** (p-value: **** < 0.0001, two-tailed unpaired t-test). Bars represent the mean for three biological replicates; error bars indicate the standard deviation. (**D**) Flow cytometry analysis of BFP expression in 624-mel MART-1^G31S^ expressing either NT- or T-SPEAR gRNAs, used in the experiments shown in Figure 3. (**E**) Schematic of the *in vivo* experimental setup for results shown in **F**. (**F**) Tumor growth analysis of 624-mel cells expressing MART-1 or MART-1^G31S^ with or without adoptive cell therapy (ACT) using MART-1-specific human-derived T cells. (**G**) Alternative visualization of the *in vivo* tumor growth shown in Figure 3E. Each curve represents the mean of six mice, and each single dot represents each mouse.

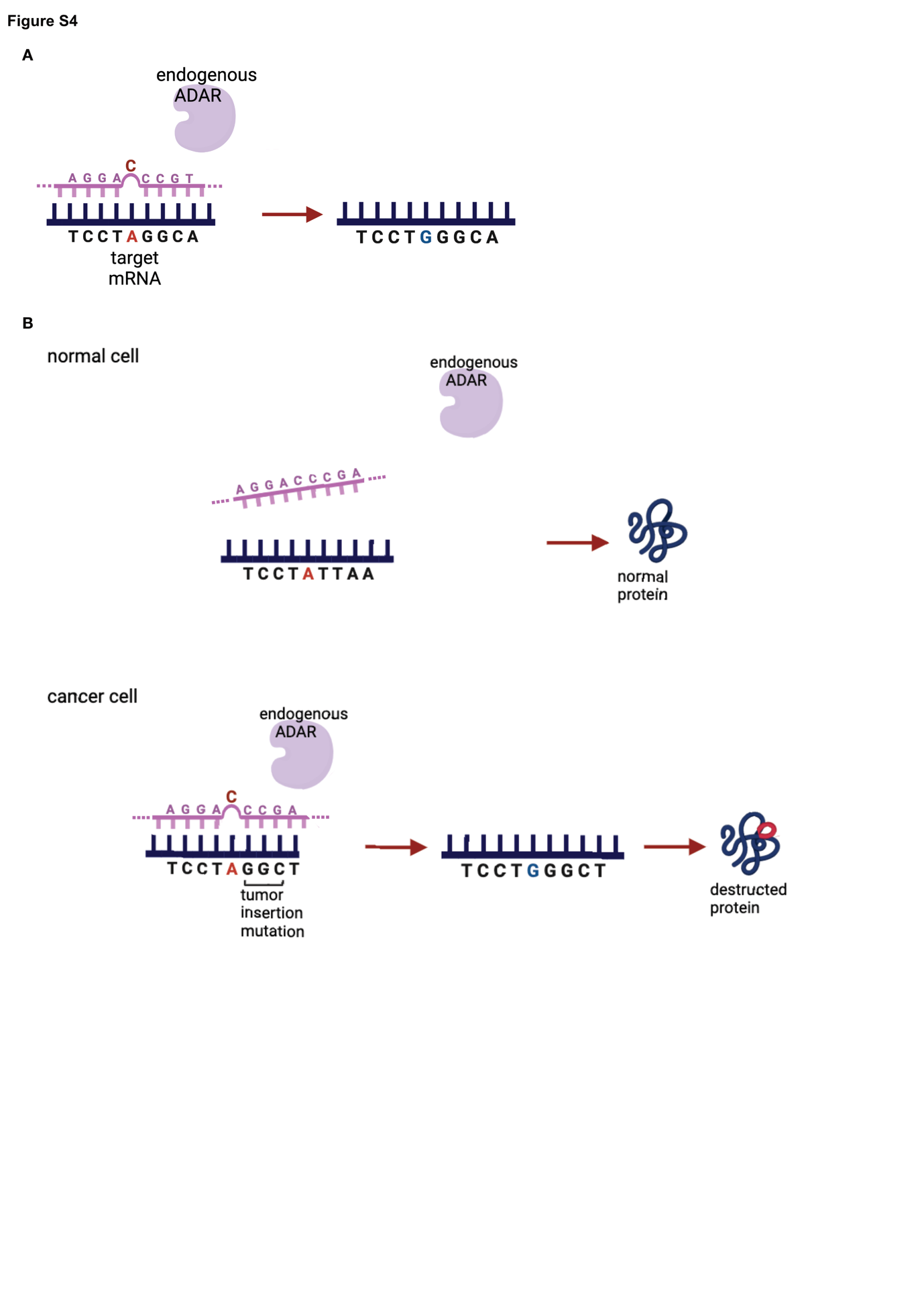

**Figure S4. Schematic presentation of Endogenous-ADAR and of our method.** (**A**) Illustration of the endogenous-ADAR editor that converts adenosine into inosine at the RNA level. The editing depends solely on a gRNA that recruits the endogenous cellular ADAR enzyme. The gRNA ensures target specificity through Watson-Crick base pairing to the desired RNA region. (**B**) A carefully designed gRNA binds to a sequence of a somatic mutation that exists in the tumor but not in healthy cells, ensuring the binding specificity in tumor cells. Then, an adenosine in close proximity is edited to an inosine (read as guanosine from the ribosome), creating a peptide that can serve as a potent neoantigen in the tumor cell.

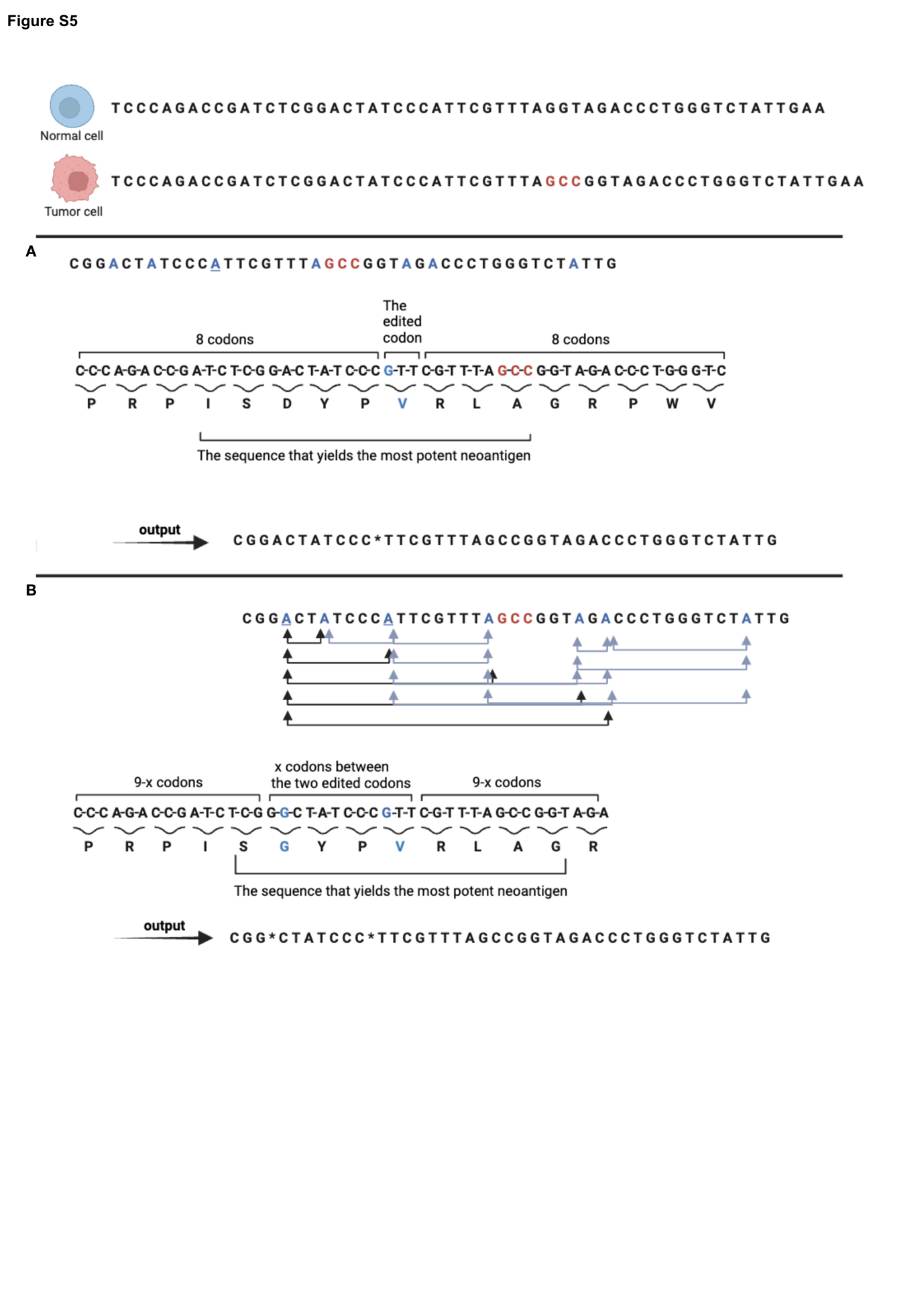

**Figure S5. Constructing the codon sequence for submission to the external prediction tool.** In this hypothetic example, an insertion mutation (GCC, marked in red) exists in the tumor cell and is absent in the normal one, and is considered for the next steps only if no potent neoantigen is currently predicted to be formed by this alteration. The algorithm retrieves the 20 bases upstream and downstream the mutation. This region serves as the locus to which the programmed gRNA binds. Within this span all adenosines are considered as possible targets for editing and the algorithm examines the potential of each to create a potent neoantigen upon conversion to guanosine. (A) Single-editing algorithm: For each candidate adenosine, the algorithm constructs a sequence spanning 17 codons—comprising the codon in which the adenosine is converted to guanosine, along with eight codons upstream and eight codons downstream. From this 17-mer sequence, the external prediction tool identifies the optimal 9-mer peptide according to the patient’s HLA type. Constructing the codon sequence in this manner ensures that the edited adenosine is included within any candidate 9-mer. As an example, we illustrate the analysis of the third adenosine within the region. The algorithm ultimately selects, among all analyzed adenosines, the optimal edit site that yields the strongest predicted neoepitope for the given somatic mutation. The output is the cDNA sequence with the edited adenosine represented by an asterisk (*). B) **Double-editing algorithm:** The same procedure is applied to every possible double-editing combination. For each candidate pair of adenosines, the algorithm constructs a codon sequence that includes all codons between the two edited sites (a total of x codons), supplemented by (9 – x) codons upstream and (9 – x) codons downstream. As a result, the generated sequence can range from a minimum of 9 codons - when the two adenosines are separated by exactly 9 codons - to a maximum of 17 codons when both adenosines reside within the same codon (pairs of adenosines located more than 9 codons apart are excluded from consideration). Constructing the codon sequence in this way ensures that both edited adenosines are encompassed within any 9-mer sequence selected by the prediction tool. As an example, we illustrate the analysis of the first and third adenosines within the region. Ultimately, the algorithm selects the optimal double-editing configuration that yields the highest predicted neoepitope score for the given somatic mutatio

c. **Plasmid maps**

To retrieve the GenBank-formatted plasmid sequence provided below, first open a plain text editor (e.g., TextEdit on macOS or Notepad on Windows). Ensure the editor is set to **Plain Text** mode (in TextEdit: Format → Make Plain Text). Then, copy the GenBank text for the plasmid of interest from below and paste it into the plain text editor. Save the file with a .gb extension to preserve formatting and enable compatibility with standard sequence analysis tools.

**Plasmid pENTR-U6 GluRv1-16 (eGFP^W58X^) sequence (GenBank format, provided as text)**

```
LOCUS       RP321_pENTR_U6_R        2915 bp ds-DNA     circular     14-SEP-2021
DEFINITION  .
ACCESSION
VERSION
SOURCE      .
  ORGANISM  .
COMMENT
COMMENT     ApEinfo:methylated:1
FEATURES             Location/Qualifiers
     misc_feature    1309..2118
                     /vntifkey="21"
                     /locus_tag="KanR"
                     /ApEinfo_fwdcolor="#7eff74"
                     /ApEinfo_revcolor="#7eff74"
                     /ApEinfo_graphicformat="arrow_data {{0 1 2 0 0 -1} {} 0}
                     width 5 offset 0"
     misc_feature    969..1013
                     /locus_tag="GluR loop"
                     /ApEinfo_fwdcolor="cyan"
                     /ApEinfo_revcolor="green"
                     /ApEinfo_graphicformat="arrow_data {{0 1 2 0 0 -1} {} 0}
                     width 5 offset 0"
     promoter        962..968
                     /vntifkey="29"
                     /locus_tag="hU6"
                     /ApEinfo_fwdcolor="#346ee0"
                     /ApEinfo_revcolor="#346ee0"
                     /ApEinfo_graphicformat="arrow_data {{0 1 2 0 0 -1} {} 0}
                     width 5 offset 0"
     promoter        705..961
                     /vntifkey="29"
                     /locus_tag="hU6(1)"
                     /ApEinfo_label="hU6"
                     /ApEinfo_fwdcolor="#346ee0"
                     /ApEinfo_revcolor="#346ee0"
                     /ApEinfo_graphicformat="arrow_data {{0 1 2 0 0 -1} {} 0}
                     width 5 offset 0"
     rep_origin      2249..2868
                     /vntifkey="33"
                     /locus_tag="pBR322ori"
                     /ApEinfo_fwdcolor="#999999"
                     /ApEinfo_revcolor="#999999"
                     /ApEinfo_graphicformat="arrow_data {{0 1 2 0 0 -1} {} 0}
                     width 5 offset 0"
     terminator      1030..1035
                     /locus_tag="PolIII terminator"
                     /ApEinfo_fwdcolor="#9d1b1c"
                     /ApEinfo_revcolor="#9d1b1c"
                     /ApEinfo_graphicformat="arrow_data {{0 1 2 0 0 -1} {} 0}
                     width 5 offset 0"
ORIGIN
        1 ctttcctgcg ttatcccctg attctgtgga taaccgtatt accgcctttg agtgagctga
       61 taccgctcgc cgcagccgaa cgaccgagcg cagcgagtca gtgagcgagg aagcggaaga
      121 gcgcccaata cgcaaaccgc ctctccccgc gcgttggccg attcattaat gcagctggca
      181 cgacaggttt cccgactgga aagcgggcag tgagcgcaac gcaattaata cgcgtaccgc
      241 tagccaggaa gagtttgtag aaacgcaaaa aggccatccg tcaggatggc cttctgctta
      301 gtttgatgcc tggcagttta tggcgggcgt cctgcccgcc accctccggg ccgttgcttc
      361 acaacgttca aatccgctcc cggcggattt gtcctactca ggagagcgtt caccgacaaa
      421 caacagataa aacgaaaggc ccagtcttcc gactgagcct ttcgttttat ttgatgcctg
      481 gcagttccct actctcgcgt taacgctagc atggatgttt tcccagtcac gacgttgtaa
      541 aacgacggcc agtcttaagc tcgggcccca aataatgatt ttattttgac tgatagtgac
      601 ctgttcgttg caacaaattg atgagcaatg cttttttata atgccaactt tgtacaaaaa
      661 agcaggcttt aaaggaacca attcagtcga ctggatccgg taccaaggtc gggcaggaag
      721 agggcctatt tcccatgatt ccttcatatt tgcatatacg atacaaggct gttagagaga
      781 taattagaat taatttgact gtaaacacaa agatattagt acaaaatacg tgacgtagaa
      841 agtaataatt tcttgggtag tttgcagttt taaaattatg ttttaaaatg gactatcata
      901 tgcttaccgt aacttgaaag tatttcgatt tcttggcttt atatatcttg tggaaaggac
      961 gaaacaccgg tgaatagtat aacaatatgc taaatgttgt tatagtatcc accgttggcC
     1021 atggaacagt tttttctaga cccagctttc ttgtacaaag ttggcattat aagaaagcat
     1081 tgcttatcaa tttgttgcaa cgaacaggtc actatcagtc aaaataaaat cattatttgc
     1141 catccagctg atatccccta tagtgagtcg tattacatgg tcatagctgt ttcctggcag
     1201 ctctggcccg tgtctcaaaa tctctgatgt tacattgcac aagataaaaa tatatcatca
     1261 tgaacaataa aactgtctgc ttacataaac agtaatacaa ggggtgttat gagccatatt
     1321 caacgggaaa cgtcgaggcc gcgattaaat tccaacatgg atgctgattt atatgggtat
     1381 aaatgggctc gcgataatgt cgggcaatca ggtgcgacaa tctatcgctt gtatgggaag
     1441 cccgatgcgc cagagttgtt tctgaaacat ggcaaaggta gcgttgccaa tgatgttaca
     1501 gatgagatgg tcagactaaa ctggctgacg gaatttatgc ctcttccgac catcaagcat
     1561 tttatccgta ctcctgatga tgcatggtta ctcaccactg cgatccccgg aaaaacagca
     1621 ttccaggtat tagaagaata tcctgattca ggtgaaaata ttgttgatgc gctggcagtg
     1681 ttcctgcgcc ggttgcattc gattcctgtt tgtaattgtc cttttaacag cgatcgcgta
     1741 tttcgtctcg ctcaggcgca atcacgaatg aataacggtt tggttgatgc gagtgatttt
     1801 gatgacgagc gtaatggctg gcctgttgaa caagtctgga aagaaatgca taaacttttg
     1861 ccattctcac cggattcagt cgtcactcat ggtgatttct cacttgataa ccttattttt
     1921 gacgagggga aattaatagg ttgtattgat gttggacgag tcggaatcgc agaccgatac
     1981 caggatcttg ccatcctatg gaactgcctc ggtgagtttt ctccttcatt acagaaacgg
     2041 ctttttcaaa aatatggtat tgataatcct gatatgaata aattgcagtt tcatttgatg
     2101 ctcgatgagt ttttctaatc agaattggtt aattggttgt aacactggca gagcattacg
     2161 ctgacttgac gggacggcgc aagctcatga ccaaaatccc ttaacgtgag ttacgcgtcg
     2221 ttccactgag cgtcagaccc cgtagaaaag atcaaaggat cttcttgaga tccttttttt
     2281 ctgcgcgtaa tctgctgctt gcaaacaaaa aaaccaccgc taccagcggt ggtttgtttg
     2341 ccggatcaag agctaccaac tctttttccg aaggtaactg gcttcagcag agcgcagata
     2401 ccaaatactg tTcttctagt gtagccgtag ttaggccacc acttcaagaa ctctgtagca
     2461 ccgcctacat acctcgctct gctaatcctg ttaccagtgg ctgctgccag tggcgataag
     2521 tcgtgtctta ccgggttgga ctcaagacga tagttaccgg ataaggcgca gcggtcgggc
     2581 tgaacggggg gttcgtgcac acagcccagc ttggagcgaa cgacctacac cgaactgaga
     2641 tacctacagc gtgagcattg agaaagcgcc acgcttcccg aagggagaaa ggcggacagg
     2701 tatccggtaa gcggcagggt cggaacagga gagcgcacga gggagcttcc agggggaaac
     2761 gcctggtatc tttatagtcc tgtcgggttt cgccacctct gacttgagcg tcgatttttg
     2821 tgatgctcgt caggggggcg gagcctatgg aaaaacgcca gcaacgcggc ctttttacgg
     2881 ttcctggcct tttgctggcc ttttgctcac atgtt
//
```

**Plasmid pENTR-U6 GluRv4-16 (eGFP^W58X^) sequence (GenBank format, provided as text)**

```
LOCUS       RP322_pENTR_U6_R        2915 bp ds-DNA     circular     14-SEP-2021
DEFINITION  .
ACCESSION
VERSION
SOURCE      .
  ORGANISM  .
COMMENT
COMMENT     ApEinfo:methylated:1
FEATURES             Location/Qualifiers
     misc_feature    1309..2118
                     /vntifkey="21"
                     /locus_tag="KanR"
                     /ApEinfo_fwdcolor="#7eff74"
                     /ApEinfo_revcolor="#7eff74"
                     /ApEinfo_graphicformat="arrow_data {{0 1 2 0 0 -1} {} 0}
                     width 5 offset 0"
     misc_feature    969..1013
                     /locus_tag="GluR loop v4"
                     /ApEinfo_fwdcolor="cyan"
                     /ApEinfo_revcolor="green"
                     /ApEinfo_graphicformat="arrow_data {{0 1 2 0 0 -1} {} 0}
                     width 5 offset 0"
     promoter        705..968
                     /vntifkey="29"
                     /locus_tag="hU6"
                     /ApEinfo_fwdcolor="#346ee0"
                     /ApEinfo_revcolor="#346ee0"
                     /ApEinfo_graphicformat="arrow_data {{0 1 2 0 0 -1} {} 0}
                     width 5 offset 0"
     rep_origin      2249..2868
                     /vntifkey="33"
                     /locus_tag="pBR322ori"
                     /ApEinfo_fwdcolor="#999999"
                     /ApEinfo_revcolor="#999999"
                     /ApEinfo_graphicformat="arrow_data {{0 1 2 0 0 -1} {} 0}
                     width 5 offset 0"
     terminator      1030..1035
                     /locus_tag="PolIII terminator"
                     /ApEinfo_fwdcolor="#9d1b1c"
                     /ApEinfo_revcolor="#9d1b1c"
                     /ApEinfo_graphicformat="arrow_data {{0 1 2 0 0 -1} {} 0}
                     width 5 offset 0"
ORIGIN
        1 ctttcctgcg ttatcccctg attctgtgga taaccgtatt accgcctttg agtgagctga
       61 taccgctcgc cgcagccgaa cgaccgagcg cagcgagtca gtgagcgagg aagcggaaga
      121 gcgcccaata cgcaaaccgc ctctccccgc gcgttggccg attcattaat gcagctggca
      181 cgacaggttt cccgactgga aagcgggcag tgagcgcaac gcaattaata cgcgtaccgc
      241 tagccaggaa gagtttgtag aaacgcaaaa aggccatccg tcaggatggc cttctgctta
      301 gtttgatgcc tggcagttta tggcgggcgt cctgcccgcc accctccggg ccgttgcttc
      361 acaacgttca aatccgctcc cggcggattt gtcctactca ggagagcgtt caccgacaaa
      421 caacagataa aacgaaaggc ccagtcttcc gactgagcct ttcgttttat ttgatgcctg
      481 gcagttccct actctcgcgt taacgctagc atggatgttt tcccagtcac gacgttgtaa
      541 aacgacggcc agtcttaagc tcgggcccca aataatgatt ttattttgac tgatagtgac
      601 ctgttcgttg caacaaattg atgagcaatg cttttttata atgccaactt tgtacaaaaa
      661 agcaggcttt aaaggaacca attcagtcga ctggatccgg taccaaggtc gggcaggaag
      721 agggcctatt tcccatgatt ccttcatatt tgcatatacg atacaaggct gttagagaga
      781 taattagaat taatttgact gtaaacacaa agatattagt acaaaatacg tgacgtagaa
      841 agtaataatt tcttgggtag tttgcagttt taaaattatg ttttaaaatg gactatcata
      901 tgcttaccgt aacttgaaag tatttcgatt tcttggcttt atatatcttg tggaaaggac
      961 gaaacaccgg tgaaGagGaG aacaatatgc taaatgttgt tCtCgtCtcc accgttggcC
     1021 atggaacagt tttttctaga cccagctttc ttgtacaaag ttggcattat aagaaagcat
     1081 tgcttatcaa tttgttgcaa cgaacaggtc actatcagtc aaaataaaat cattatttgc
     1141 catccagctg atatccccta tagtgagtcg tattacatgg tcatagctgt ttcctggcag
     1201 ctctggcccg tgtctcaaaa tctctgatgt tacattgcac aagataaaaa tatatcatca
     1261 tgaacaataa aactgtctgc ttacataaac agtaatacaa ggggtgttat gagccatatt
     1321 caacgggaaa cgtcgaggcc gcgattaaat tccaacatgg atgctgattt atatgggtat
     1381 aaatgggctc gcgataatgt cgggcaatca ggtgcgacaa tctatcgctt gtatgggaag
     1441 cccgatgcgc cagagttgtt tctgaaacat ggcaaaggta gcgttgccaa tgatgttaca
     1501 gatgagatgg tcagactaaa ctggctgacg gaatttatgc ctcttccgac catcaagcat
     1561 tttatccgta ctcctgatga tgcatggtta ctcaccactg cgatccccgg aaaaacagca
     1621 ttccaggtat tagaagaata tcctgattca ggtgaaaata ttgttgatgc gctggcagtg
     1681 ttcctgcgcc ggttgcattc gattcctgtt tgtaattgtc cttttaacag cgatcgcgta
     1741 tttcgtctcg ctcaggcgca atcacgaatg aataacggtt tggttgatgc gagtgatttt
     1801 gatgacgagc gtaatggctg gcctgttgaa caagtctgga aagaaatgca taaacttttg
     1861 ccattctcac cggattcagt cgtcactcat ggtgatttct cacttgataa ccttattttt
     1921 gacgagggga aattaatagg ttgtattgat gttggacgag tcggaatcgc agaccgatac
     1981 caggatcttg ccatcctatg gaactgcctc ggtgagtttt ctccttcatt acagaaacgg
     2041 ctttttcaaa aatatggtat tgataatcct gatatgaata aattgcagtt tcatttgatg
     2101 ctcgatgagt ttttctaatc agaattggtt aattggttgt aacactggca gagcattacg
     2161 ctgacttgac gggacggcgc aagctcatga ccaaaatccc ttaacgtgag ttacgcgtcg
     2221 ttccactgag cgtcagaccc cgtagaaaag atcaaaggat cttcttgaga tccttttttt
     2281 ctgcgcgtaa tctgctgctt gcaaacaaaa aaaccaccgc taccagcggt ggtttgtttg
     2341 ccggatcaag agctaccaac tctttttccg aaggtaactg gcttcagcag agcgcagata
     2401 ccaaatactg tTcttctagt gtagccgtag ttaggccacc acttcaagaa ctctgtagca
     2461 ccgcctacat acctcgctct gctaatcctg ttaccagtgg ctgctgccag tggcgataag
     2521 tcgtgtctta ccgggttgga ctcaagacga tagttaccgg ataaggcgca gcggtcgggc
     2581 tgaacggggg gttcgtgcac acagcccagc ttggagcgaa cgacctacac cgaactgaga
     2641 tacctacagc gtgagcattg agaaagcgcc acgcttcccg aagggagaaa ggcggacagg
     2701 tatccggtaa gcggcagggt cggaacagga gagcgcacga gggagcttcc agggggaaac
     2761 gcctggtatc tttatagtcc tgtcgggttt cgccacctct gacttgagcg tcgatttttg
     2821 tgatgctcgt caggggggcg gagcctatgg aaaaacgcca gcaacgcggc ctttttacgg
     2881 ttcctggcct tttgctggcc ttttgctcac atgtt
//
```

**Plasmid pENTR-U6 GluRv9.4-16 (eGFP^W58X^) sequence (GenBank format, provided as text)**

```
LOCUS       323_RP_pENTR_U6_        2925 bp ds-DNA     circular     14-SEP-2021
DEFINITION  .
ACCESSION
VERSION
SOURCE      .
  ORGANISM  .
COMMENT
COMMENT
COMMENT     ApEinfo:methylated:1
FEATURES             Location/Qualifiers
     misc_feature    1319..2128
                     /vntifkey="21"
                     /locus_tag="KanR"
                     /ApEinfo_fwdcolor="#7eff74"
                     /ApEinfo_revcolor="#7eff74"
                     /ApEinfo_graphicformat="arrow_data {{0 1 2 0 0 -1} {} 0}
                     width 5 offset 0"
     promoter        705..968
                     /vntifkey="29"
                     /locus_tag="hU6"
                     /ApEinfo_fwdcolor="#346ee0"
                     /ApEinfo_revcolor="#346ee0"
                     /ApEinfo_graphicformat="arrow_data {{0 1 2 0 0 -1} {} 0}
                     width 5 offset 0"
     rep_origin      2259..2878
                     /vntifkey="33"
                     /locus_tag="pBR322ori"
                     /ApEinfo_fwdcolor="#999999"
                     /ApEinfo_revcolor="#999999"
                     /ApEinfo_graphicformat="arrow_data {{0 1 2 0 0 -1} {} 0}
                     width 5 offset 0"
     terminator      1040..1045
                     /locus_tag="PolIII terminator"
                     /ApEinfo_fwdcolor="#9d1b1c"
                     /ApEinfo_revcolor="#9d1b1c"
                     /ApEinfo_graphicformat="arrow_data {{0 1 2 0 0 -1} {} 0}
                     width 5 offset 0"
     misc_feature    969..1023
                     /locus_tag="GluR loop"
                     /ApEinfo_fwdcolor="cyan"
                     /ApEinfo_revcolor="green"
                     /ApEinfo_graphicformat="arrow_data {{0 1 2 0 0 -1} {} 0}
                     width 5 offset 0"
ORIGIN
        1 ctttcctgcg ttatcccctg attctgtgga taaccgtatt accgcctttg agtgagctga
       61 taccgctcgc cgcagccgaa cgaccgagcg cagcgagtca gtgagcgagg aagcggaaga
      121 gcgcccaata cgcaaaccgc ctctccccgc gcgttggccg attcattaat gcagctggca
      181 cgacaggttt cccgactgga aagcgggcag tgagcgcaac gcaattaata cgcgtaccgc
      241 tagccaggaa gagtttgtag aaacgcaaaa aggccatccg tcaggatggc cttctgctta
      301 gtttgatgcc tggcagttta tggcgggcgt cctgcccgcc accctccggg ccgttgcttc
      361 acaacgttca aatccgctcc cggcggattt gtcctactca ggagagcgtt caccgacaaa
      421 caacagataa aacgaaaggc ccagtcttcc gactgagcct ttcgttttat ttgatgcctg
      481 gcagttccct actctcgcgt taacgctagc atggatgttt tcccagtcac gacgttgtaa
      541 aacgacggcc agtcttaagc tcgggcccca aataatgatt ttattttgac tgatagtgac
      601 ctgttcgttg caacaaattg atgagcaatg cttttttata atgccaactt tgtacaaaaa
      661 agcaggcttt aaaggaacca attcagtcga ctggatccgg taccaaggtc gggcaggaag
      721 agggcctatt tcccatgatt ccttcatatt tgcatatacg atacaaggct gttagagaga
      781 taattagaat taatttgact gtaaacacaa agatattagt acaaaatacg tgacgtagaa
      841 agtaataatt tcttgggtag tttgcagttt taaaattatg ttttaaaatg gactatcata
      901 tgcttaccgt aacttgaaag tatttcgatt tcttggcttt atatatcttg tggaaaggac
      961 gaaacaccgg tgTCGAGaaG agGaGaacaa tatgctaaat gttgttCtCg tCtcCTCGAc
     1021 accgttggcC atggaacagt tttttctaga cccagctttc ttgtacaaag ttggcattat
     1081 aagaaagcat tgcttatcaa tttgttgcaa cgaacaggtc actatcagtc aaaataaaat
     1141 cattatttgc catccagctg atatccccta tagtgagtcg tattacatgg tcatagctgt
     1201 ttcctggcag ctctggcccg tgtctcaaaa tctctgatgt tacattgcac aagataaaaa
     1261 tatatcatca tgaacaataa aactgtctgc ttacataaac agtaatacaa ggggtgttat
     1321 gagccatatt caacgggaaa cgtcgaggcc gcgattaaat tccaacatgg atgctgattt
     1381 atatgggtat aaatgggctc gcgataatgt cgggcaatca ggtgcgacaa tctatcgctt
     1441 gtatgggaag cccgatgcgc cagagttgtt tctgaaacat ggcaaaggta gcgttgccaa
     1501 tgatgttaca gatgagatgg tcagactaaa ctggctgacg gaatttatgc ctcttccgac
     1561 catcaagcat tttatccgta ctcctgatga tgcatggtta ctcaccactg cgatccccgg
     1621 aaaaacagca ttccaggtat tagaagaata tcctgattca ggtgaaaata ttgttgatgc
     1681 gctggcagtg ttcctgcgcc ggttgcattc gattcctgtt tgtaattgtc cttttaacag
     1741 cgatcgcgta tttcgtctcg ctcaggcgca atcacgaatg aataacggtt tggttgatgc
     1801 gagtgatttt gatgacgagc gtaatggctg gcctgttgaa caagtctgga aagaaatgca
     1861 taaacttttg ccattctcac cggattcagt cgtcactcat ggtgatttct cacttgataa
     1921 ccttattttt gacgagggga aattaatagg ttgtattgat gttggacgag tcggaatcgc
     1981 agaccgatac caggatcttg ccatcctatg gaactgcctc ggtgagtttt ctccttcatt
     2041 acagaaacgg ctttttcaaa aatatggtat tgataatcct gatatgaata aattgcagtt
     2101 tcatttgatg ctcgatgagt ttttctaatc agaattggtt aattggttgt aacactggca
     2161 gagcattacg ctgacttgac gggacggcgc aagctcatga ccaaaatccc ttaacgtgag
     2221 ttacgcgtcg ttccactgag cgtcagaccc cgtagaaaag atcaaaggat cttcttgaga
     2281 tccttttttt ctgcgcgtaa tctgctgctt gcaaacaaaa aaaccaccgc taccagcggt
     2341 ggtttgtttg ccggatcaag agctaccaac tctttttccg aaggtaactg gcttcagcag
     2401 agcgcagata ccaaatactg tTcttctagt gtagccgtag ttaggccacc acttcaagaa
     2461 ctctgtagca ccgcctacat acctcgctct gctaatcctg ttaccagtgg ctgctgccag
     2521 tggcgataag tcgtgtctta ccgggttgga ctcaagacga tagttaccgg ataaggcgca
     2581 gcggtcgggc tgaacggggg gttcgtgcac acagcccagc ttggagcgaa cgacctacac
     2641 cgaactgaga tacctacagc gtgagcattg agaaagcgcc acgcttcccg aagggagaaa
     2701 ggcggacagg tatccggtaa gcggcagggt cggaacagga gagcgcacga gggagcttcc
     2761 agggggaaac gcctggtatc tttatagtcc tgtcgggttt cgccacctct gacttgagcg
     2821 tcgatttttg tgatgctcgt caggggggcg gagcctatgg aaaaacgcca gcaacgcggc
     2881 ctttttacgg ttcctggcct tttgctggcc ttttgctcac atgtt
//
```

**Plasmid pENTR-U6 GluRv9.4-38 (eGFP^W58X^) sequence (GenBank format, provided as text)**

```
LOCUS       RP324_pENTR_U6_R        2947 bp ds-DNA     circular     14-SEP-2021
DEFINITION  .
ACCESSION
VERSION
SOURCE      .
  ORGANISM  .
COMMENT
COMMENT
COMMENT     ApEinfo:methylated:1
FEATURES             Location/Qualifiers
     misc_feature    1341..2150
                     /vntifkey="21"
                     /locus_tag="KanR"
                     /ApEinfo_fwdcolor="#7eff74"
                     /ApEinfo_revcolor="#7eff74"
                     /ApEinfo_graphicformat="arrow_data {{0 1 2 0 0 -1} {} 0}
                     width 5 offset 0"
     misc_feature    969..1023
                     /locus_tag="GluR loop"
                     /ApEinfo_fwdcolor="cyan"
                     /ApEinfo_revcolor="green"
                     /ApEinfo_graphicformat="arrow_data {{0 1 2 0 0 -1} {} 0}
                     width 5 offset 0"
     promoter        705..968
                     /vntifkey="29"
                     /locus_tag="hU6"
                     /ApEinfo_fwdcolor="#346ee0"
                     /ApEinfo_revcolor="#346ee0"
                     /ApEinfo_graphicformat="arrow_data {{0 1 2 0 0 -1} {} 0}
                     width 5 offset 0"
     rep_origin      2281..2900
                     /vntifkey="33"
                     /locus_tag="pBR322ori"
                     /ApEinfo_fwdcolor="#999999"
                     /ApEinfo_revcolor="#999999"
                     /ApEinfo_graphicformat="arrow_data {{0 1 2 0 0 -1} {} 0}
                     width 5 offset 0"
     terminator      1062..1067
                     /locus_tag="PolIII terminator"
                     /ApEinfo_fwdcolor="#9d1b1c"
                     /ApEinfo_revcolor="#9d1b1c"
                     /ApEinfo_graphicformat="arrow_data {{0 1 2 0 0 -1} {} 0}
                     width 5 offset 0"
ORIGIN
        1 ctttcctgcg ttatcccctg attctgtgga taaccgtatt accgcctttg agtgagctga
       61 taccgctcgc cgcagccgaa cgaccgagcg cagcgagtca gtgagcgagg aagcggaaga
      121 gcgcccaata cgcaaaccgc ctctccccgc gcgttggccg attcattaat gcagctggca
      181 cgacaggttt cccgactgga aagcgggcag tgagcgcaac gcaattaata cgcgtaccgc
      241 tagccaggaa gagtttgtag aaacgcaaaa aggccatccg tcaggatggc cttctgctta
      301 gtttgatgcc tggcagttta tggcgggcgt cctgcccgcc accctccggg ccgttgcttc
      361 acaacgttca aatccgctcc cggcggattt gtcctactca ggagagcgtt caccgacaaa
      421 caacagataa aacgaaaggc ccagtcttcc gactgagcct ttcgttttat ttgatgcctg
      481 gcagttccct actctcgcgt taacgctagc atggatgttt tcccagtcac gacgttgtaa
      541 aacgacggcc agtcttaagc tcgggcccca aataatgatt ttattttgac tgatagtgac
      601 ctgttcgttg caacaaattg atgagcaatg cttttttata atgccaactt tgtacaaaaa
      661 agcaggcttt aaaggaacca attcagtcga ctggatccgg taccaaggtc gggcaggaag
      721 agggcctatt tcccatgatt ccttcatatt tgcatatacg atacaaggct gttagagaga
      781 taattagaat taatttgact gtaaacacaa agatattagt acaaaatacg tgacgtagaa
      841 agtaataatt tcttgggtag tttgcagttt taaaattatg ttttaaaatg gactatcata
      901 tgcttaccgt aacttgaaag tatttcgatt tcttggcttt atatatcttg tggaaaggac
      961 gaaacaccgg tgTCGAGaaG agGaGaacaa tatgctaaat gttgttCtCg tCtcCTCGAc
     1021 accgtatgtc agggtagtga caagtgttgg cCatggaaca gttttttcta gacccagctt
     1081 tcttgtacaa agttggcatt ataagaaagc attgcttatc aatttgttgc aacgaacagg
     1141 tcactatcag tcaaaataaa atcattattt gccatccagc tgatatcccc tatagtgagt
     1201 cgtattacat ggtcatagct gtttcctggc agctctggcc cgtgtctcaa aatctctgat
     1261 gttacattgc acaagataaa aatatatcat catgaacaat aaaactgtct gcttacataa
     1321 acagtaatac aaggggtgtt atgagccata ttcaacggga aacgtcgagg ccgcgattaa
     1381 attccaacat ggatgctgat ttatatgggt ataaatgggc tcgcgataat gtcgggcaat
     1441 caggtgcgac aatctatcgc ttgtatggga agcccgatgc gccagagttg tttctgaaac
     1501 atggcaaagg tagcgttgcc aatgatgtta cagatgagat ggtcagacta aactggctga
     1561 cggaatttat gcctcttccg accatcaagc attttatccg tactcctgat gatgcatggt
     1621 tactcaccac tgcgatcccc ggaaaaacag cattccaggt attagaagaa tatcctgatt
     1681 caggtgaaaa tattgttgat gcgctggcag tgttcctgcg ccggttgcat tcgattcctg
     1741 tttgtaattg tccttttaac agcgatcgcg tatttcgtct cgctcaggcg caatcacgaa
     1801 tgaataacgg tttggttgat gcgagtgatt ttgatgacga gcgtaatggc tggcctgttg
     1861 aacaagtctg gaaagaaatg cataaacttt tgccattctc accggattca gtcgtcactc
     1921 atggtgattt ctcacttgat aaccttattt ttgacgaggg gaaattaata ggttgtattg
     1981 atgttggacg agtcggaatc gcagaccgat accaggatct tgccatccta tggaactgcc
     2041 tcggtgagtt ttctccttca ttacagaaac ggctttttca aaaatatggt attgataatc
     2101 ctgatatgaa taaattgcag tttcatttga tgctcgatga gtttttctaa tcagaattgg
     2161 ttaattggtt gtaacactgg cagagcatta cgctgacttg acgggacggc gcaagctcat
     2221 gaccaaaatc ccttaacgtg agttacgcgt cgttccactg agcgtcagac cccgtagaaa
     2281 agatcaaagg atcttcttga gatccttttt ttctgcgcgt aatctgctgc ttgcaaacaa
     2341 aaaaaccacc gctaccagcg gtggtttgtt tgccggatca agagctacca actctttttc
     2401 cgaaggtaac tggcttcagc agagcgcaga taccaaatac tgtTcttcta gtgtagccgt
     2461 agttaggcca ccacttcaag aactctgtag caccgcctac atacctcgct ctgctaatcc
     2521 tgttaccagt ggctgctgcc agtggcgata agtcgtgtct taccgggttg gactcaagac
     2581 gatagttacc ggataaggcg cagcggtcgg gctgaacggg gggttcgtgc acacagccca
     2641 gcttggagcg aacgacctac accgaactga gatacctaca gcgtgagcat tgagaaagcg
     2701 ccacgcttcc cgaagggaga aaggcggaca ggtatccggt aagcggcagg gtcggaacag
     2761 gagagcgcac gagggagctt ccagggggaa acgcctggta tctttatagt cctgtcgggt
     2821 ttcgccacct ctgacttgag cgtcgatttt tgtgatgctc gtcagggggg cggagcctat
     2881 ggaaaaacgc cagcaacgcg gcctttttac ggttcctggc cttttgctgg ccttttgctc
     2941 acatgtt
//
```

**Plasmid pENTR-U6 LEAPER-70 (eGFP^W58X^) sequence (GenBank format, provided as text)**

```
LOCUS       RP325_pENTR_U6_L        2926 bp ds-DNA     circular     14-SEP-2021
DEFINITION  .
ACCESSION
VERSION
SOURCE      .
  ORGANISM  .
COMMENT
COMMENT
COMMENT     ApEinfo:methylated:1
FEATURES             Location/Qualifiers
     misc_feature    1320..2129
                     /vntifkey="21"
                     /locus_tag="KanR"
                     /ApEinfo_fwdcolor="#7eff74"
                     /ApEinfo_revcolor="#7eff74"
                     /ApEinfo_graphicformat="arrow_data {{0 1 2 0 0 -1} {} 0}
                     width 5 offset 0"
     promoter        962..968
                     /vntifkey="29"
                     /locus_tag="hU6"
                     /ApEinfo_fwdcolor="#346ee0"
                     /ApEinfo_revcolor="#346ee0"
                     /ApEinfo_graphicformat="arrow_data {{0 1 2 0 0 -1} {} 0}
                     width 5 offset 0"
     promoter        705..961
                     /vntifkey="29"
                     /locus_tag="hU6(1)"
                     /ApEinfo_label="hU6"
                     /ApEinfo_fwdcolor="#346ee0"
                     /ApEinfo_revcolor="#346ee0"
                     /ApEinfo_graphicformat="arrow_data {{0 1 2 0 0 -1} {} 0}
                     width 5 offset 0"
     rep_origin      2260..2879
                     /vntifkey="33"
                     /locus_tag="pBR322ori"
                     /ApEinfo_fwdcolor="#999999"
                     /ApEinfo_revcolor="#999999"
                     /ApEinfo_graphicformat="arrow_data {{0 1 2 0 0 -1} {} 0}
                     width 5 offset 0"
     terminator      1041..1046
                     /locus_tag="PolIII terminator"
                     /ApEinfo_fwdcolor="#9d1b1c"
                     /ApEinfo_revcolor="#9d1b1c"
                     /ApEinfo_graphicformat="arrow_data {{0 1 2 0 0 -1} {} 0}
                     width 5 offset 0"
ORIGIN
        1 ctttcctgcg ttatcccctg attctgtgga taaccgtatt accgcctttg agtgagctga
       61 taccgctcgc cgcagccgaa cgaccgagcg cagcgagtca gtgagcgagg aagcggaaga
      121 gcgcccaata cgcaaaccgc ctctccccgc gcgttggccg attcattaat gcagctggca
      181 cgacaggttt cccgactgga aagcgggcag tgagcgcaac gcaattaata cgcgtaccgc
      241 tagccaggaa gagtttgtag aaacgcaaaa aggccatccg tcaggatggc cttctgctta
      301 gtttgatgcc tggcagttta tggcgggcgt cctgcccgcc accctccggg ccgttgcttc
      361 acaacgttca aatccgctcc cggcggattt gtcctactca ggagagcgtt caccgacaaa
      421 caacagataa aacgaaaggc ccagtcttcc gactgagcct ttcgttttat ttgatgcctg
      481 gcagttccct actctcgcgt taacgctagc atggatgttt tcccagtcac gacgttgtaa
      541 aacgacggcc agtcttaagc tcgggcccca aataatgatt ttattttgac tgatagtgac
      601 ctgttcgttg caacaaattg atgagcaatg cttttttata atgccaactt tgtacaaaaa
      661 agcaggcttt aaaggaacca attcagtcga ctggatccgg taccaaggtc gggcaggaag
      721 agggcctatt tcccatgatt ccttcatatt tgcatatacg atacaaggct gttagagaga
      781 taattagaat taatttgact gtaaacacaa agatattagt acaaaatacg tgacgtagaa
      841 agtaataatt tcttgggtag tttgcagttt taaaattatg ttttaaaatg gactatcata
      901 tgcttaccgt aacttgaaag tatttcgatt tcttggcttt atatatcttg tggaaaggac
      961 gaaacaccGG attgcacgcc gtatgtcagg gtagtgacaa gtgttggcCa tggaacaggt
     1021 agttttccag tagtgcaaat ttttttctag acccagcttt cttgtacaaa gttggcatta
     1081 taagaaagca ttgcttatca atttgttgca acgaacaggt cactatcagt caaaataaaa
     1141 tcattatttg ccatccagct gatatcccct atagtgagtc gtattacatg gtcatagctg
     1201 tttcctggca gctctggccc gtgtctcaaa atctctgatg ttacattgca caagataaaa
     1261 atatatcatc atgaacaata aaactgtctg cttacataaa cagtaataca aggggtgtta
     1321 tgagccatat tcaacgggaa acgtcgaggc cgcgattaaa ttccaacatg gatgctgatt
     1381 tatatgggta taaatgggct cgcgataatg tcgggcaatc aggtgcgaca atctatcgct
     1441 tgtatgggaa gcccgatgcg ccagagttgt ttctgaaaca tggcaaaggt agcgttgcca
     1501 atgatgttac agatgagatg gtcagactaa actggctgac ggaatttatg cctcttccga
     1561 ccatcaagca ttttatccgt actcctgatg atgcatggtt actcaccact gcgatccccg
     1621 gaaaaacagc attccaggta ttagaagaat atcctgattc aggtgaaaat attgttgatg
     1681 cgctggcagt gttcctgcgc cggttgcatt cgattcctgt ttgtaattgt ccttttaaca
     1741 gcgatcgcgt atttcgtctc gctcaggcgc aatcacgaat gaataacggt ttggttgatg
     1801 cgagtgattt tgatgacgag cgtaatggct ggcctgttga acaagtctgg aaagaaatgc
     1861 ataaactttt gccattctca ccggattcag tcgtcactca tggtgatttc tcacttgata
     1921 accttatttt tgacgagggg aaattaatag gttgtattga tgttggacga gtcggaatcg
     1981 cagaccgata ccaggatctt gccatcctat ggaactgcct cggtgagttt tctccttcat
     2041 tacagaaacg gctttttcaa aaatatggta ttgataatcc tgatatgaat aaattgcagt
     2101 ttcatttgat gctcgatgag tttttctaat cagaattggt taattggttg taacactggc
     2161 agagcattac gctgacttga cgggacggcg caagctcatg accaaaatcc cttaacgtga
     2221 gttacgcgtc gttccactga gcgtcagacc ccgtagaaaa gatcaaagga tcttcttgag
     2281 atcctttttt tctgcgcgta atctgctgct tgcaaacaaa aaaaccaccg ctaccagcgg
     2341 tggtttgttt gccggatcaa gagctaccaa ctctttttcc gaaggtaact ggcttcagca
     2401 gagcgcagat accaaatact gtTcttctag tgtagccgta gttaggccac cacttcaaga
     2461 actctgtagc accgcctaca tacctcgctc tgctaatcct gttaccagtg gctgctgcca
     2521 gtggcgataa gtcgtgtctt accgggttgg actcaagacg atagttaccg gataaggcgc
     2581 agcggtcggg ctgaacgggg ggttcgtgca cacagcccag cttggagcga acgacctaca
     2641 ccgaactgag atacctacag cgtgagcatt gagaaagcgc cacgcttccc gaagggagaa
     2701 aggcggacag gtatccggta agcggcaggg tcggaacagg agagcgcacg agggagcttc
     2761 cagggggaaa cgcctggtat ctttatagtc ctgtcgggtt tcgccacctc tgacttgagc
     2821 gtcgattttt gtgatgctcg tcaggggggc ggagcctatg gaaaaacgcc agcaacgcgg
     2881 cctttttacg gttcctggcc ttttgctggc cttttgctca catgtt
//
```

**Plasmid pENTR-U6 LEAPER-151 (Qu et al 2019) (eGFP^W58X^) sequence (GenBank format, provided as text)**

```
LOCUS       RP326_pENTR_U6_L        3007 bp ds-DNA     circular     14-SEP-2021
DEFINITION  .
ACCESSION
VERSION
SOURCE      .
  ORGANISM  .
COMMENT
COMMENT     ApEinfo:methylated:1
FEATURES             Location/Qualifiers
     misc_feature    1401..2210
                     /vntifkey="21"
                     /locus_tag="KanR"
                     /ApEinfo_fwdcolor="#7eff74"
                     /ApEinfo_revcolor="#7eff74"
                     /ApEinfo_graphicformat="arrow_data {{0 1 2 0 0 -1} {} 0}
                     width 5 offset 0"
     promoter        962..968
                     /vntifkey="29"
                     /locus_tag="hU6"
                     /ApEinfo_fwdcolor="#346ee0"
                     /ApEinfo_revcolor="#346ee0"
                     /ApEinfo_graphicformat="arrow_data {{0 1 2 0 0 -1} {} 0}
                     width 5 offset 0"
     promoter        705..961
                     /vntifkey="29"
                     /locus_tag="hU6(1)"
                     /ApEinfo_label="hU6"
                     /ApEinfo_fwdcolor="#346ee0"
                     /ApEinfo_revcolor="#346ee0"
                     /ApEinfo_graphicformat="arrow_data {{0 1 2 0 0 -1} {} 0}
                     width 5 offset 0"
     rep_origin      2341..2960
                     /vntifkey="33"
                     /locus_tag="pBR322ori"
                     /ApEinfo_fwdcolor="#999999"
                     /ApEinfo_revcolor="#999999"
                     /ApEinfo_graphicformat="arrow_data {{0 1 2 0 0 -1} {} 0}
                     width 5 offset 0"
     terminator      1122..1127
                     /locus_tag="PolIII terminator"
                     /ApEinfo_fwdcolor="#9d1b1c"
                     /ApEinfo_revcolor="#9d1b1c"
                     /ApEinfo_graphicformat="arrow_data {{0 1 2 0 0 -1} {} 0}
                     width 5 offset 0"
ORIGIN
        1 ctttcctgcg ttatcccctg attctgtgga taaccgtatt accgcctttg agtgagctga
       61 taccgctcgc cgcagccgaa cgaccgagcg cagcgagtca gtgagcgagg aagcggaaga
      121 gcgcccaata cgcaaaccgc ctctccccgc gcgttggccg attcattaat gcagctggca
      181 cgacaggttt cccgactgga aagcgggcag tgagcgcaac gcaattaata cgcgtaccgc
      241 tagccaggaa gagtttgtag aaacgcaaaa aggccatccg tcaggatggc cttctgctta
      301 gtttgatgcc tggcagttta tggcgggcgt cctgcccgcc accctccggg ccgttgcttc
      361 acaacgttca aatccgctcc cggcggattt gtcctactca ggagagcgtt caccgacaaa
      421 caacagataa aacgaaaggc ccagtcttcc gactgagcct ttcgttttat ttgatgcctg
      481 gcagttccct actctcgcgt taacgctagc atggatgttt tcccagtcac gacgttgtaa
      541 aacgacggcc agtcttaagc tcgggcccca aataatgatt ttattttgac tgatagtgac
      601 ctgttcgttg caacaaattg atgagcaatg cttttttata atgccaactt tgtacaaaaa
      661 agcaggcttt aaaggaacca attcagtcga ctggatccgg taccaaggtc gggcaggaag
      721 agggcctatt tcccatgatt ccttcatatt tgcatatacg atacaaggct gttagagaga
      781 taattagaat taatttgact gtaaacacaa agatattagt acaaaatacg tgacgtagaa
      841 agtaataatt tcttgggtag tttgcagttt taaaattatg ttttaaaatg gactatcata
      901 tgcttaccgt aacttgaaag tatttcgatt tcttggcttt atatatcttg tggaaaggac
      961 gaaacaccGG tcgtgctgtt tcatgtggtc ggggtacctg ctgaagcatt gcacgccgta
     1021 tgtcagggta gtgacaagtg ttggcCatgg aacaggtagt tttccagtag tgcaaataaa
     1081 cttcagggtc agcttgccat aggtggcatc gccctctcct tttttttcta gacccagctt
     1141 tcttgtacaa agttggcatt ataagaaagc attgcttatc aatttgttgc aacgaacagg
     1201 tcactatcag tcaaaataaa atcattattt gccatccagc tgatatcccc tatagtgagt
     1261 cgtattacat ggtcatagct gtttcctggc agctctggcc cgtgtctcaa aatctctgat
     1321 gttacattgc acaagataaa aatatatcat catgaacaat aaaactgtct gcttacataa
     1381 acagtaatac aaggggtgtt atgagccata ttcaacggga aacgtcgagg ccgcgattaa
     1441 attccaacat ggatgctgat ttatatgggt ataaatgggc tcgcgataat gtcgggcaat
     1501 caggtgcgac aatctatcgc ttgtatggga agcccgatgc gccagagttg tttctgaaac
     1561 atggcaaagg tagcgttgcc aatgatgtta cagatgagat ggtcagacta aactggctga
     1621 cggaatttat gcctcttccg accatcaagc attttatccg tactcctgat gatgcatggt
     1681 tactcaccac tgcgatcccc ggaaaaacag cattccaggt attagaagaa tatcctgatt
     1741 caggtgaaaa tattgttgat gcgctggcag tgttcctgcg ccggttgcat tcgattcctg
     1801 tttgtaattg tccttttaac agcgatcgcg tatttcgtct cgctcaggcg caatcacgaa
     1861 tgaataacgg tttggttgat gcgagtgatt ttgatgacga gcgtaatggc tggcctgttg
     1921 aacaagtctg gaaagaaatg cataaacttt tgccattctc accggattca gtcgtcactc
     1981 atggtgattt ctcacttgat aaccttattt ttgacgaggg gaaattaata ggttgtattg
     2041 atgttggacg agtcggaatc gcagaccgat accaggatct tgccatccta tggaactgcc
     2101 tcggtgagtt ttctccttca ttacagaaac ggctttttca aaaatatggt attgataatc
     2161 ctgatatgaa taaattgcag tttcatttga tgctcgatga gtttttctaa tcagaattgg
     2221 ttaattggtt gtaacactgg cagagcatta cgctgacttg acgggacggc gcaagctcat
     2281 gaccaaaatc ccttaacgtg agttacgcgt cgttccactg agcgtcagac cccgtagaaa
     2341 agatcaaagg atcttcttga gatccttttt ttctgcgcgt aatctgctgc ttgcaaacaa
     2401 aaaaaccacc gctaccagcg gtggtttgtt tgccggatca agagctacca actctttttc
     2461 cgaaggtaac tggcttcagc agagcgcaga taccaaatac tgtTcttcta gtgtagccgt
     2521 agttaggcca ccacttcaag aactctgtag caccgcctac atacctcgct ctgctaatcc
     2581 tgttaccagt ggctgctgcc agtggcgata agtcgtgtct taccgggttg gactcaagac
     2641 gatagttacc ggataaggcg cagcggtcgg gctgaacggg gggttcgtgc acacagccca
     2701 gcttggagcg aacgacctac accgaactga gatacctaca gcgtgagcat tgagaaagcg
     2761 ccacgcttcc cgaagggaga aaggcggaca ggtatccggt aagcggcagg gtcggaacag
     2821 gagagcgcac gagggagctt ccagggggaa acgcctggta tctttatagt cctgtcgggt
     2881 ttcgccacct ctgacttgag cgtcgatttt tgtgatgctc gtcagggggg cggagcctat
     2941 ggaaaaacgc cagcaacgcg gcctttttac ggttcctggc cttttgctgg ccttttgctc
     3001 acatgtt
//
```

**Plasmid pENTR-U6 LEAPER-151 (38-4ntMM-33-C-75; Uzonyi et al 2021) (eGFP^W58X^) sequence (GenBank format, provided as text)**

```
LOCUS       RP327_pENTR_U6_L        3007 bp ds-DNA     circular     23-OCT-2021
DEFINITION  .
ACCESSION
VERSION
SOURCE      .
  ORGANISM  .
COMMENT
COMMENT     ApEinfo:methylated:1
FEATURES             Location/Qualifiers
     misc_feature    1401..2210
                     /vntifkey="21"
                     /locus_tag="KanR"
                     /ApEinfo_fwdcolor="#7eff74"
                     /ApEinfo_revcolor="#7eff74"
                     /ApEinfo_graphicformat="arrow_data {{0 1 2 0 0 -1} {} 0}
                     width 5 offset 0"
     promoter        962..968
                     /vntifkey="29"
                     /locus_tag="hU6"
                     /ApEinfo_fwdcolor="#346ee0"
                     /ApEinfo_revcolor="#346ee0"
                     /ApEinfo_graphicformat="arrow_data {{0 1 2 0 0 -1} {} 0}
                     width 5 offset 0"
     promoter        705..961
                     /vntifkey="29"
                     /locus_tag="hU6(1)"
                     /ApEinfo_label="hU6"
                     /ApEinfo_fwdcolor="#346ee0"
                     /ApEinfo_revcolor="#346ee0"
                     /ApEinfo_graphicformat="arrow_data {{0 1 2 0 0 -1} {} 0}
                     width 5 offset 0"
     rep_origin      2341..2960
                     /vntifkey="33"
                     /locus_tag="pBR322ori"
                     /ApEinfo_fwdcolor="#999999"
                     /ApEinfo_revcolor="#999999"
                     /ApEinfo_graphicformat="arrow_data {{0 1 2 0 0 -1} {} 0}
                     width 5 offset 0"
     terminator      1122..1127
                     /locus_tag="PolIII terminator"
                     /ApEinfo_fwdcolor="#9d1b1c"
                     /ApEinfo_revcolor="#9d1b1c"
                     /ApEinfo_graphicformat="arrow_data {{0 1 2 0 0 -1} {} 0}
                     width 5 offset 0"
     misc_binding    1009..1012
                     /locus_tag="4nt mismatch (as in Uzonyi et al 2021)"
                     /ApEinfo_fwdcolor="#ff4809"
                     /ApEinfo_revcolor="#ff4809"
                     /ApEinfo_graphicformat="arrow_data {{0 1 2 0 0 -1} {} 0}
                     width 5 offset 0"
ORIGIN
        1 ctttcctgcg ttatcccctg attctgtgga taaccgtatt accgcctttg agtgagctga
       61 taccgctcgc cgcagccgaa cgaccgagcg cagcgagtca gtgagcgagg aagcggaaga
      121 gcgcccaata cgcaaaccgc ctctccccgc gcgttggccg attcattaat gcagctggca
      181 cgacaggttt cccgactgga aagcgggcag tgagcgcaac gcaattaata cgcgtaccgc
      241 tagccaggaa gagtttgtag aaacgcaaaa aggccatccg tcaggatggc cttctgctta
      301 gtttgatgcc tggcagttta tggcgggcgt cctgcccgcc accctccggg ccgttgcttc
      361 acaacgttca aatccgctcc cggcggattt gtcctactca ggagagcgtt caccgacaaa
      421 caacagataa aacgaaaggc ccagtcttcc gactgagcct ttcgttttat ttgatgcctg
      481 gcagttccct actctcgcgt taacgctagc atggatgttt tcccagtcac gacgttgtaa
      541 aacgacggcc agtcttaagc tcgggcccca aataatgatt ttattttgac tgatagtgac
      601 ctgttcgttg caacaaattg atgagcaatg cttttttata atgccaactt tgtacaaaaa
      661 agcaggcttt aaaggaacca attcagtcga ctggatccgg taccaaggtc gggcaggaag
      721 agggcctatt tcccatgatt ccttcatatt tgcatatacg atacaaggct gttagagaga
      781 taattagaat taatttgact gtaaacacaa agatattagt acaaaatacg tgacgtagaa
      841 agtaataatt tcttgggtag tttgcagttt taaaattatg ttttaaaatg gactatcata
      901 tgcttaccgt aacttgaaag tatttcgatt tcttggcttt atatatcttg tggaaaggac
      961 gaaacaccGG tcgtgctgtt tcatgtggtc ggggtacctg ctgaagcaaa cgacgccgta
     1021 tgtcagggta gtgacaagtg ttggcCatgg aacaggtagt tttccagtag tgcaaataaa
     1081 cttcagggtc agcttgccat aggtggcatc gccctctcct tttttttcta gacccagctt
     1141 tcttgtacaa agttggcatt ataagaaagc attgcttatc aatttgttgc aacgaacagg
     1201 tcactatcag tcaaaataaa atcattattt gccatccagc tgatatcccc tatagtgagt
     1261 cgtattacat ggtcatagct gtttcctggc agctctggcc cgtgtctcaa aatctctgat
     1321 gttacattgc acaagataaa aatatatcat catgaacaat aaaactgtct gcttacataa
     1381 acagtaatac aaggggtgtt atgagccata ttcaacggga aacgtcgagg ccgcgattaa
     1441 attccaacat ggatgctgat ttatatgggt ataaatgggc tcgcgataat gtcgggcaat
     1501 caggtgcgac aatctatcgc ttgtatggga agcccgatgc gccagagttg tttctgaaac
     1561 atggcaaagg tagcgttgcc aatgatgtta cagatgagat ggtcagacta aactggctga
     1621 cggaatttat gcctcttccg accatcaagc attttatccg tactcctgat gatgcatggt
     1681 tactcaccac tgcgatcccc ggaaaaacag cattccaggt attagaagaa tatcctgatt
     1741 caggtgaaaa tattgttgat gcgctggcag tgttcctgcg ccggttgcat tcgattcctg
     1801 tttgtaattg tccttttaac agcgatcgcg tatttcgtct cgctcaggcg caatcacgaa
     1861 tgaataacgg tttggttgat gcgagtgatt ttgatgacga gcgtaatggc tggcctgttg
     1921 aacaagtctg gaaagaaatg cataaacttt tgccattctc accggattca gtcgtcactc
     1981 atggtgattt ctcacttgat aaccttattt ttgacgaggg gaaattaata ggttgtattg
     2041 atgttggacg agtcggaatc gcagaccgat accaggatct tgccatccta tggaactgcc
     2101 tcggtgagtt ttctccttca ttacagaaac ggctttttca aaaatatggt attgataatc
     2161 ctgatatgaa taaattgcag tttcatttga tgctcgatga gtttttctaa tcagaattgg
     2221 ttaattggtt gtaacactgg cagagcatta cgctgacttg acgggacggc gcaagctcat
     2281 gaccaaaatc ccttaacgtg agttacgcgt cgttccactg agcgtcagac cccgtagaaa
     2341 agatcaaagg atcttcttga gatccttttt ttctgcgcgt aatctgctgc ttgcaaacaa
     2401 aaaaaccacc gctaccagcg gtggtttgtt tgccggatca agagctacca actctttttc
     2461 cgaaggtaac tggcttcagc agagcgcaga taccaaatac tgtTcttcta gtgtagccgt
     2521 agttaggcca ccacttcaag aactctgtag caccgcctac atacctcgct ctgctaatcc
     2581 tgttaccagt ggctgctgcc agtggcgata agtcgtgtct taccgggttg gactcaagac
     2641 gatagttacc ggataaggcg cagcggtcgg gctgaacggg gggttcgtgc acacagccca
     2701 gcttggagcg aacgacctac accgaactga gatacctaca gcgtgagcat tgagaaagcg
     2761 ccacgcttcc cgaagggaga aaggcggaca ggtatccggt aagcggcagg gtcggaacag
     2821 gagagcgcac gagggagctt ccagggggaa acgcctggta tctttatagt cctgtcgggt
     2881 ttcgccacct ctgacttgag cgtcgatttt tgtgatgctc gtcagggggg cggagcctat
     2941 ggaaaaacgc cagcaacgcg gcctttttac ggttcctggc cttttgctgg ccttttgctc
     3001 acatgtt
//
```

**Plasmid pENTR-U6 38-4ntMM-33-C-31 (sdl 107) (eGFP^W58X^) sequence (GenBank format, provided as text)**

```
LOCUS       RP381_pENTR_U6_L        2963 bp ds-DNA     circular     14-MAR-2022
DEFINITION  .
ACCESSION
VERSION
SOURCE      .
  ORGANISM  .
COMMENT
COMMENT
COMMENT     ApEinfo:methylated:1
FEATURES             Location/Qualifiers
     misc_feature    1357..2166
                     /vntifkey="21"
                     /locus_tag="KanR"
                     /ApEinfo_fwdcolor="#7eff74"
                     /ApEinfo_revcolor="#7eff74"
                     /ApEinfo_graphicformat="arrow_data {{0 1 2 0 0 -1} {} 0}
                     width 5 offset 0"
     promoter        962..968
                     /vntifkey="29"
                     /locus_tag="hU6"
                     /ApEinfo_fwdcolor="#346ee0"
                     /ApEinfo_revcolor="#346ee0"
                     /ApEinfo_graphicformat="arrow_data {{0 1 2 0 0 -1} {} 0}
                     width 5 offset 0"
     promoter        705..961
                     /vntifkey="29"
                     /locus_tag="hU6(1)"
                     /ApEinfo_label="hU6"
                     /ApEinfo_fwdcolor="#346ee0"
                     /ApEinfo_revcolor="#346ee0"
                     /ApEinfo_graphicformat="arrow_data {{0 1 2 0 0 -1} {} 0}
                     width 5 offset 0"
     rep_origin      2297..2916
                     /vntifkey="33"
                     /locus_tag="pBR322ori"
                     /ApEinfo_fwdcolor="#999999"
                     /ApEinfo_revcolor="#999999"
                     /ApEinfo_graphicformat="arrow_data {{0 1 2 0 0 -1} {} 0}
                     width 5 offset 0"
     terminator      1078..1083
                     /locus_tag="PolIII terminator"
                     /ApEinfo_fwdcolor="#9d1b1c"
                     /ApEinfo_revcolor="#9d1b1c"
                     /ApEinfo_graphicformat="arrow_data {{0 1 2 0 0 -1} {} 0}
                     width 5 offset 0"
     misc_binding    1009..1012
                     /locus_tag="4nt mismatch (as in Uzonyi et al 2021)"
                     /ApEinfo_fwdcolor="#ff4809"
                     /ApEinfo_revcolor="#ff4809"
                     /ApEinfo_graphicformat="arrow_data {{0 1 2 0 0 -1} {} 0}
                     width 5 offset 0"
ORIGIN
        1 ctttcctgcg ttatcccctg attctgtgga taaccgtatt accgcctttg agtgagctga
       61 taccgctcgc cgcagccgaa cgaccgagcg cagcgagtca gtgagcgagg aagcggaaga
      121 gcgcccaata cgcaaaccgc ctctccccgc gcgttggccg attcattaat gcagctggca
      181 cgacaggttt cccgactgga aagcgggcag tgagcgcaac gcaattaata cgcgtaccgc
      241 tagccaggaa gagtttgtag aaacgcaaaa aggccatccg tcaggatggc cttctgctta
      301 gtttgatgcc tggcagttta tggcgggcgt cctgcccgcc accctccggg ccgttgcttc
      361 acaacgttca aatccgctcc cggcggattt gtcctactca ggagagcgtt caccgacaaa
      421 caacagataa aacgaaaggc ccagtcttcc gactgagcct ttcgttttat ttgatgcctg
      481 gcagttccct actctcgcgt taacgctagc atggatgttt tcccagtcac gacgttgtaa
      541 aacgacggcc agtcttaagc tcgggcccca aataatgatt ttattttgac tgatagtgac
      601 ctgttcgttg caacaaattg atgagcaatg cttttttata atgccaactt tgtacaaaaa
      661 agcaggcttt aaaggaacca attcagtcga ctggatccgg taccaaggtc gggcaggaag
      721 agggcctatt tcccatgatt ccttcatatt tgcatatacg atacaaggct gttagagaga
      781 taattagaat taatttgact gtaaacacaa agatattagt acaaaatacg tgacgtagaa
      841 agtaataatt tcttgggtag tttgcagttt taaaattatg ttttaaaatg gactatcata
      901 tgcttaccgt aacttgaaag tatttcgatt tcttggcttt atatatcttg tggaaaggac
      961 gaaacaccGG tcgtgctgtt tcatgtggtc ggggtacctg ctgaagcaaa cgacgccgta
     1021 tgtcagggta gtgacaagtg ttggcCatgg aacaggtagt tttccagtag tgcaaatttt
     1081 tttctagacc cagctttctt gtacaaagtt ggcattataa gaaagcattg cttatcaatt
     1141 tgttgcaacg aacaggtcac tatcagtcaa aataaaatca ttatttgcca tccagctgat
     1201 atcccctata gtgagtcgta ttacatggtc atagctgttt cctggcagct ctggcccgtg
     1261 tctcaaaatc tctgatgtta cattgcacaa gataaaaata tatcatcatg aacaataaaa
     1321 ctgtctgctt acataaacag taatacaagg ggtgttatga gccatattca acgggaaacg
     1381 tcgaggccgc gattaaattc caacatggat gctgatttat atgggtataa atgggctcgc
     1441 gataatgtcg ggcaatcagg tgcgacaatc tatcgcttgt atgggaagcc cgatgcgcca
     1501 gagttgtttc tgaaacatgg caaaggtagc gttgccaatg atgttacaga tgagatggtc
     1561 agactaaact ggctgacgga atttatgcct cttccgacca tcaagcattt tatccgtact
     1621 cctgatgatg catggttact caccactgcg atccccggaa aaacagcatt ccaggtatta
     1681 gaagaatatc ctgattcagg tgaaaatatt gttgatgcgc tggcagtgtt cctgcgccgg
     1741 ttgcattcga ttcctgtttg taattgtcct tttaacagcg atcgcgtatt tcgtctcgct
     1801 caggcgcaat cacgaatgaa taacggtttg gttgatgcga gtgattttga tgacgagcgt
     1861 aatggctggc ctgttgaaca agtctggaaa gaaatgcata aacttttgcc attctcaccg
     1921 gattcagtcg tcactcatgg tgatttctca cttgataacc ttatttttga cgaggggaaa
     1981 ttaataggtt gtattgatgt tggacgagtc ggaatcgcag accgatacca ggatcttgcc
     2041 atcctatgga actgcctcgg tgagttttct ccttcattac agaaacggct ttttcaaaaa
     2101 tatggtattg ataatcctga tatgaataaa ttgcagtttc atttgatgct cgatgagttt
     2161 ttctaatcag aattggttaa ttggttgtaa cactggcaga gcattacgct gacttgacgg
     2221 gacggcgcaa gctcatgacc aaaatccctt aacgtgagtt acgcgtcgtt ccactgagcg
     2281 tcagaccccg tagaaaagat caaaggatct tcttgagatc ctttttttct gcgcgtaatc
     2341 tgctgcttgc aaacaaaaaa accaccgcta ccagcggtgg tttgtttgcc ggatcaagag
     2401 ctaccaactc tttttccgaa ggtaactggc ttcagcagag cgcagatacc aaatactgtT
     2461 cttctagtgt agccgtagtt aggccaccac ttcaagaact ctgtagcacc gcctacatac
     2521 ctcgctctgc taatcctgtt accagtggct gctgccagtg gcgataagtc gtgtcttacc
     2581 gggttggact caagacgata gttaccggat aaggcgcagc ggtcgggctg aacggggggt
     2641 tcgtgcacac agcccagctt ggagcgaacg acctacaccg aactgagata cctacagcgt
     2701 gagcattgag aaagcgccac gcttcccgaa gggagaaagg cggacaggta tccggtaagc
     2761 ggcagggtcg gaacaggaga gcgcacgagg gagcttccag ggggaaacgc ctggtatctt
     2821 tatagtcctg tcgggtttcg ccacctctga cttgagcgtc gatttttgtg atgctcgtca
     2881 ggggggcgga gcctatggaa aaacgccagc aacgcggcct ttttacggtt cctggccttt
     2941 tgctggcctt ttgctcacat gtt
//
```

**Plasmid pENTR-U6 38-4ntMM-33-C-16 (sdl 92) (eGFP^W58X^) sequence (GenBank format, provided as text)**

```
LOCUS       RP382_pENTR_U6_L        2948 bp ds-DNA     circular     14-MAR-2022
DEFINITION  .
ACCESSION
VERSION
SOURCE      .
  ORGANISM  .
COMMENT
COMMENT     ApEinfo:methylated:1
FEATURES             Location/Qualifiers
     misc_feature    1342..2151
                     /vntifkey="21"
                     /locus_tag="KanR"
                     /ApEinfo_fwdcolor="#7eff74"
                     /ApEinfo_revcolor="#7eff74"
                     /ApEinfo_graphicformat="arrow_data {{0 1 2 0 0 -1} {} 0}
                     width 5 offset 0"
     promoter        962..968
                     /vntifkey="29"
                     /locus_tag="hU6"
                     /ApEinfo_fwdcolor="#346ee0"
                     /ApEinfo_revcolor="#346ee0"
                     /ApEinfo_graphicformat="arrow_data {{0 1 2 0 0 -1} {} 0}
                     width 5 offset 0"
     promoter        705..961
                     /vntifkey="29"
                     /locus_tag="hU6(1)"
                     /ApEinfo_label="hU6"
                     /ApEinfo_fwdcolor="#346ee0"
                     /ApEinfo_revcolor="#346ee0"
                     /ApEinfo_graphicformat="arrow_data {{0 1 2 0 0 -1} {} 0}
                     width 5 offset 0"
     rep_origin      2282..2901
                     /vntifkey="33"
                     /locus_tag="pBR322ori"
                     /ApEinfo_fwdcolor="#999999"
                     /ApEinfo_revcolor="#999999"
                     /ApEinfo_graphicformat="arrow_data {{0 1 2 0 0 -1} {} 0}
                     width 5 offset 0"
     terminator      1063..1068
                     /locus_tag="PolIII terminator"
                     /ApEinfo_fwdcolor="#9d1b1c"
                     /ApEinfo_revcolor="#9d1b1c"
                     /ApEinfo_graphicformat="arrow_data {{0 1 2 0 0 -1} {} 0}
                     width 5 offset 0"
     misc_binding    1009..1012
                     /locus_tag="4nt mismatch (as in Uzonyi et al 2021)"
                     /ApEinfo_fwdcolor="#ff4809"
                     /ApEinfo_revcolor="#ff4809"
                     /ApEinfo_graphicformat="arrow_data {{0 1 2 0 0 -1} {} 0}
                     width 5 offset 0"
ORIGIN
        1 ctttcctgcg ttatcccctg attctgtgga taaccgtatt accgcctttg agtgagctga
       61 taccgctcgc cgcagccgaa cgaccgagcg cagcgagtca gtgagcgagg aagcggaaga
      121 gcgcccaata cgcaaaccgc ctctccccgc gcgttggccg attcattaat gcagctggca
      181 cgacaggttt cccgactgga aagcgggcag tgagcgcaac gcaattaata cgcgtaccgc
      241 tagccaggaa gagtttgtag aaacgcaaaa aggccatccg tcaggatggc cttctgctta
      301 gtttgatgcc tggcagttta tggcgggcgt cctgcccgcc accctccggg ccgttgcttc
      361 acaacgttca aatccgctcc cggcggattt gtcctactca ggagagcgtt caccgacaaa
      421 caacagataa aacgaaaggc ccagtcttcc gactgagcct ttcgttttat ttgatgcctg
      481 gcagttccct actctcgcgt taacgctagc atggatgttt tcccagtcac gacgttgtaa
      541 aacgacggcc agtcttaagc tcgggcccca aataatgatt ttattttgac tgatagtgac
      601 ctgttcgttg caacaaattg atgagcaatg cttttttata atgccaactt tgtacaaaaa
      661 agcaggcttt aaaggaacca attcagtcga ctggatccgg taccaaggtc gggcaggaag
      721 agggcctatt tcccatgatt ccttcatatt tgcatatacg atacaaggct gttagagaga
      781 taattagaat taatttgact gtaaacacaa agatattagt acaaaatacg tgacgtagaa
      841 agtaataatt tcttgggtag tttgcagttt taaaattatg ttttaaaatg gactatcata
      901 tgcttaccgt aacttgaaag tatttcgatt tcttggcttt atatatcttg tggaaaggac
      961 gaaacaccGG tcgtgctgtt tcatgtggtc ggggtacctg ctgaagcaaa cgacgccgta
     1021 tgtcagggta gtgacaagtg ttggcCatgg aacaggtagt ttttttttct agacccagct
     1081 ttcttgtaca aagttggcat tataagaaag cattgcttat caatttgttg caacgaacag
     1141 gtcactatca gtcaaaataa aatcattatt tgccatccag ctgatatccc ctatagtgag
     1201 tcgtattaca tggtcatagc tgtttcctgg cagctctggc ccgtgtctca aaatctctga
     1261 tgttacattg cacaagataa aaatatatca tcatgaacaa taaaactgtc tgcttacata
     1321 aacagtaata caaggggtgt tatgagccat attcaacggg aaacgtcgag gccgcgatta
     1381 aattccaaca tggatgctga tttatatggg tataaatggg ctcgcgataa tgtcgggcaa
     1441 tcaggtgcga caatctatcg cttgtatggg aagcccgatg cgccagagtt gtttctgaaa
     1501 catggcaaag gtagcgttgc caatgatgtt acagatgaga tggtcagact aaactggctg
     1561 acggaattta tgcctcttcc gaccatcaag cattttatcc gtactcctga tgatgcatgg
     1621 ttactcacca ctgcgatccc cggaaaaaca gcattccagg tattagaaga atatcctgat
     1681 tcaggtgaaa atattgttga tgcgctggca gtgttcctgc gccggttgca ttcgattcct
     1741 gtttgtaatt gtccttttaa cagcgatcgc gtatttcgtc tcgctcaggc gcaatcacga
     1801 atgaataacg gtttggttga tgcgagtgat tttgatgacg agcgtaatgg ctggcctgtt
     1861 gaacaagtct ggaaagaaat gcataaactt ttgccattct caccggattc agtcgtcact
     1921 catggtgatt tctcacttga taaccttatt tttgacgagg ggaaattaat aggttgtatt
     1981 gatgttggac gagtcggaat cgcagaccga taccaggatc ttgccatcct atggaactgc
     2041 ctcggtgagt tttctccttc attacagaaa cggctttttc aaaaatatgg tattgataat
     2101 cctgatatga ataaattgca gtttcatttg atgctcgatg agtttttcta atcagaattg
     2161 gttaattggt tgtaacactg gcagagcatt acgctgactt gacgggacgg cgcaagctca
     2221 tgaccaaaat cccttaacgt gagttacgcg tcgttccact gagcgtcaga ccccgtagaa
     2281 aagatcaaag gatcttcttg agatcctttt tttctgcgcg taatctgctg cttgcaaaca
     2341 aaaaaaccac cgctaccagc ggtggtttgt ttgccggatc aagagctacc aactcttttt
     2401 ccgaaggtaa ctggcttcag cagagcgcag ataccaaata ctgtTcttct agtgtagccg
     2461 tagttaggcc accacttcaa gaactctgta gcaccgccta catacctcgc tctgctaatc
     2521 ctgttaccag tggctgctgc cagtggcgat aagtcgtgtc ttaccgggtt ggactcaaga
     2581 cgatagttac cggataaggc gcagcggtcg ggctgaacgg ggggttcgtg cacacagccc
     2641 agcttggagc gaacgaccta caccgaactg agatacctac agcgtgagca ttgagaaagc
     2701 gccacgcttc ccgaagggag aaaggcggac aggtatccgg taagcggcag ggtcggaaca
     2761 ggagagcgca cgagggagct tccaggggga aacgcctggt atctttatag tcctgtcggg
     2821 tttcgccacc tctgacttga gcgtcgattt ttgtgatgct cgtcaggggg gcggagccta
     2881 tggaaaaacg ccagcaacgc ggccttttta cggttcctgg ccttttgctg gccttttgct
     2941 cacatgtt
//
```

**Plasmid pENTR-U6 20-4ntMM-33-C-75 (sdl 133) (eGFP^W58X^) sequence (GenBank format, provided as text)**

```
LOCUS       RP383_pENTR_U6_L        2989 bp ds-DNA     circular     14-MAR-2022
DEFINITION  .
ACCESSION
VERSION
SOURCE      .
  ORGANISM  .
COMMENT
COMMENT
COMMENT     ApEinfo:methylated:1
FEATURES             Location/Qualifiers
     misc_feature    1383..2192
                     /vntifkey="21"
                     /locus_tag="KanR"
                     /ApEinfo_fwdcolor="#7eff74"
                     /ApEinfo_revcolor="#7eff74"
                     /ApEinfo_graphicformat="arrow_data {{0 1 2 0 0 -1} {} 0}
                     width 5 offset 0"
     promoter        962..968
                     /vntifkey="29"
                     /locus_tag="hU6"
                     /ApEinfo_fwdcolor="#346ee0"
                     /ApEinfo_revcolor="#346ee0"
                     /ApEinfo_graphicformat="arrow_data {{0 1 2 0 0 -1} {} 0}
                     width 5 offset 0"
     promoter        705..961
                     /vntifkey="29"
                     /locus_tag="hU6(1)"
                     /ApEinfo_label="hU6"
                     /ApEinfo_fwdcolor="#346ee0"
                     /ApEinfo_revcolor="#346ee0"
                     /ApEinfo_graphicformat="arrow_data {{0 1 2 0 0 -1} {} 0}
                     width 5 offset 0"
     rep_origin      2323..2942
                     /vntifkey="33"
                     /locus_tag="pBR322ori"
                     /ApEinfo_fwdcolor="#999999"
                     /ApEinfo_revcolor="#999999"
                     /ApEinfo_graphicformat="arrow_data {{0 1 2 0 0 -1} {} 0}
                     width 5 offset 0"
     terminator      1104..1109
                     /locus_tag="PolIII terminator"
                     /ApEinfo_fwdcolor="#9d1b1c"
                     /ApEinfo_revcolor="#9d1b1c"
                     /ApEinfo_graphicformat="arrow_data {{0 1 2 0 0 -1} {} 0}
                     width 5 offset 0"
     misc_binding    991..994
                     /locus_tag="4nt mismatch (as in Uzonyi et al 2021)"
                     /ApEinfo_fwdcolor="#ff4809"
                     /ApEinfo_revcolor="#ff4809"
                     /ApEinfo_graphicformat="arrow_data {{0 1 2 0 0 -1} {} 0}
                     width 5 offset 0"
ORIGIN
        1 ctttcctgcg ttatcccctg attctgtgga taaccgtatt accgcctttg agtgagctga
       61 taccgctcgc cgcagccgaa cgaccgagcg cagcgagtca gtgagcgagg aagcggaaga
      121 gcgcccaata cgcaaaccgc ctctccccgc gcgttggccg attcattaat gcagctggca
      181 cgacaggttt cccgactgga aagcgggcag tgagcgcaac gcaattaata cgcgtaccgc
      241 tagccaggaa gagtttgtag aaacgcaaaa aggccatccg tcaggatggc cttctgctta
      301 gtttgatgcc tggcagttta tggcgggcgt cctgcccgcc accctccggg ccgttgcttc
      361 acaacgttca aatccgctcc cggcggattt gtcctactca ggagagcgtt caccgacaaa
      421 caacagataa aacgaaaggc ccagtcttcc gactgagcct ttcgttttat ttgatgcctg
      481 gcagttccct actctcgcgt taacgctagc atggatgttt tcccagtcac gacgttgtaa
      541 aacgacggcc agtcttaagc tcgggcccca aataatgatt ttattttgac tgatagtgac
      601 ctgttcgttg caacaaattg atgagcaatg cttttttata atgccaactt tgtacaaaaa
      661 agcaggcttt aaaggaacca attcagtcga ctggatccgg taccaaggtc gggcaggaag
      721 agggcctatt tcccatgatt ccttcatatt tgcatatacg atacaaggct gttagagaga
      781 taattagaat taatttgact gtaaacacaa agatattagt acaaaatacg tgacgtagaa
      841 agtaataatt tcttgggtag tttgcagttt taaaattatg ttttaaaatg gactatcata
      901 tgcttaccgt aacttgaaag tatttcgatt tcttggcttt atatatcttg tggaaaggac
      961 gaaacaccGG tcggggtacc tgctgaagca aacgacgccg tatgtcaggg tagtgacaag
     1021 tgttggcCat ggaacaggta gttttccagt agtgcaaata aacttcaggg tcagcttgcc
     1081 ataggtggca tcgccctctc cttttttttc tagacccagc tttcttgtac aaagttggca
     1141 ttataagaaa gcattgctta tcaatttgtt gcaacgaaca ggtcactatc agtcaaaata
     1201 aaatcattat ttgccatcca gctgatatcc cctatagtga gtcgtattac atggtcatag
     1261 ctgtttcctg gcagctctgg cccgtgtctc aaaatctctg atgttacatt gcacaagata
     1321 aaaatatatc atcatgaaca ataaaactgt ctgcttacat aaacagtaat acaaggggtg
     1381 ttatgagcca tattcaacgg gaaacgtcga ggccgcgatt aaattccaac atggatgctg
     1441 atttatatgg gtataaatgg gctcgcgata atgtcgggca atcaggtgcg acaatctatc
     1501 gcttgtatgg gaagcccgat gcgccagagt tgtttctgaa acatggcaaa ggtagcgttg
     1561 ccaatgatgt tacagatgag atggtcagac taaactggct gacggaattt atgcctcttc
     1621 cgaccatcaa gcattttatc cgtactcctg atgatgcatg gttactcacc actgcgatcc
     1681 ccggaaaaac agcattccag gtattagaag aatatcctga ttcaggtgaa aatattgttg
     1741 atgcgctggc agtgttcctg cgccggttgc attcgattcc tgtttgtaat tgtcctttta
     1801 acagcgatcg cgtatttcgt ctcgctcagg cgcaatcacg aatgaataac ggtttggttg
     1861 atgcgagtga ttttgatgac gagcgtaatg gctggcctgt tgaacaagtc tggaaagaaa
     1921 tgcataaact tttgccattc tcaccggatt cagtcgtcac tcatggtgat ttctcacttg
     1981 ataaccttat ttttgacgag gggaaattaa taggttgtat tgatgttgga cgagtcggaa
     2041 tcgcagaccg ataccaggat cttgccatcc tatggaactg cctcggtgag ttttctcctt
     2101 cattacagaa acggcttttt caaaaatatg gtattgataa tcctgatatg aataaattgc
     2161 agtttcattt gatgctcgat gagtttttct aatcagaatt ggttaattgg ttgtaacact
     2221 ggcagagcat tacgctgact tgacgggacg gcgcaagctc atgaccaaaa tcccttaacg
     2281 tgagttacgc gtcgttccac tgagcgtcag accccgtaga aaagatcaaa ggatcttctt
     2341 gagatccttt ttttctgcgc gtaatctgct gcttgcaaac aaaaaaacca ccgctaccag
     2401 cggtggtttg tttgccggat caagagctac caactctttt tccgaaggta actggcttca
     2461 gcagagcgca gataccaaat actgtTcttc tagtgtagcc gtagttaggc caccacttca
     2521 agaactctgt agcaccgcct acatacctcg ctctgctaat cctgttacca gtggctgctg
     2581 ccagtggcga taagtcgtgt cttaccgggt tggactcaag acgatagtta ccggataagg
     2641 cgcagcggtc gggctgaacg gggggttcgt gcacacagcc cagcttggag cgaacgacct
     2701 acaccgaact gagataccta cagcgtgagc attgagaaag cgccacgctt cccgaaggga
     2761 gaaaggcgga caggtatccg gtaagcggca gggtcggaac aggagagcgc acgagggagc
     2821 ttccaggggg aaacgcctgg tatctttata gtcctgtcgg gtttcgccac ctctgacttg
     2881 agcgtcgatt tttgtgatgc tcgtcagggg ggcggagcct atggaaaaac gccagcaacg
     2941 cggccttttt acggttcctg gccttttgct ggccttttgc tcacatgtt
//
```

**Plasmid pENTR-U6 10-4ntMM-33-C-75 (sdl 133) (eGFP^W58X^) sequence (GenBank format, provided as text)**

```
LOCUS       RP384_pENTR_U6_L        2979 bp ds-DNA     circular     14-MAR-2022
DEFINITION  .
ACCESSION
VERSION
SOURCE      .
  ORGANISM  .
COMMENT
COMMENT     ApEinfo:methylated:1
FEATURES             Location/Qualifiers
     misc_feature    1373..2182
                     /vntifkey="21"
                     /locus_tag="KanR"
                     /ApEinfo_fwdcolor="#7eff74"
                     /ApEinfo_revcolor="#7eff74"
                     /ApEinfo_graphicformat="arrow_data {{0 1 2 0 0 -1} {} 0}
                     width 5 offset 0"
     promoter        962..968
                     /vntifkey="29"
                     /locus_tag="hU6"
                     /ApEinfo_fwdcolor="#346ee0"
                     /ApEinfo_revcolor="#346ee0"
                     /ApEinfo_graphicformat="arrow_data {{0 1 2 0 0 -1} {} 0}
                     width 5 offset 0"
     promoter        705..961
                     /vntifkey="29"
                     /locus_tag="hU6(1)"
                     /ApEinfo_label="hU6"
                     /ApEinfo_fwdcolor="#346ee0"
                     /ApEinfo_revcolor="#346ee0"
                     /ApEinfo_graphicformat="arrow_data {{0 1 2 0 0 -1} {} 0}
                     width 5 offset 0"
     rep_origin      2313..2932
                     /vntifkey="33"
                     /locus_tag="pBR322ori"
                     /ApEinfo_fwdcolor="#999999"
                     /ApEinfo_revcolor="#999999"
                     /ApEinfo_graphicformat="arrow_data {{0 1 2 0 0 -1} {} 0}
                     width 5 offset 0"
     terminator      1094..1099
                     /locus_tag="PolIII terminator"
                     /ApEinfo_fwdcolor="#9d1b1c"
                     /ApEinfo_revcolor="#9d1b1c"
                     /ApEinfo_graphicformat="arrow_data {{0 1 2 0 0 -1} {} 0}
                     width 5 offset 0"
     misc_binding    981..984
                     /locus_tag="4nt mismatch (as in Uzonyi et al 2021)"
                     /ApEinfo_fwdcolor="#ff4809"
                     /ApEinfo_revcolor="#ff4809"
                     /ApEinfo_graphicformat="arrow_data {{0 1 2 0 0 -1} {} 0}
                     width 5 offset 0"
ORIGIN
        1 ctttcctgcg ttatcccctg attctgtgga taaccgtatt accgcctttg agtgagctga
       61 taccgctcgc cgcagccgaa cgaccgagcg cagcgagtca gtgagcgagg aagcggaaga
      121 gcgcccaata cgcaaaccgc ctctccccgc gcgttggccg attcattaat gcagctggca
      181 cgacaggttt cccgactgga aagcgggcag tgagcgcaac gcaattaata cgcgtaccgc
      241 tagccaggaa gagtttgtag aaacgcaaaa aggccatccg tcaggatggc cttctgctta
      301 gtttgatgcc tggcagttta tggcgggcgt cctgcccgcc accctccggg ccgttgcttc
      361 acaacgttca aatccgctcc cggcggattt gtcctactca ggagagcgtt caccgacaaa
      421 caacagataa aacgaaaggc ccagtcttcc gactgagcct ttcgttttat ttgatgcctg
      481 gcagttccct actctcgcgt taacgctagc atggatgttt tcccagtcac gacgttgtaa
      541 aacgacggcc agtcttaagc tcgggcccca aataatgatt ttattttgac tgatagtgac
      601 ctgttcgttg caacaaattg atgagcaatg cttttttata atgccaactt tgtacaaaaa
      661 agcaggcttt aaaggaacca attcagtcga ctggatccgg taccaaggtc gggcaggaag
      721 agggcctatt tcccatgatt ccttcatatt tgcatatacg atacaaggct gttagagaga
      781 taattagaat taatttgact gtaaacacaa agatattagt acaaaatacg tgacgtagaa
      841 agtaataatt tcttgggtag tttgcagttt taaaattatg ttttaaaatg gactatcata
      901 tgcttaccgt aacttgaaag tatttcgatt tcttggcttt atatatcttg tggaaaggac
      961 gaaacaccGG tgctgaagca aacgacgccg tatgtcaggg tagtgacaag tgttggcCat
     1021 ggaacaggta gttttccagt agtgcaaata aacttcaggg tcagcttgcc ataggtggca
     1081 tcgccctctc cttttttttc tagacccagc tttcttgtac aaagttggca ttataagaaa
     1141 gcattgctta tcaatttgtt gcaacgaaca ggtcactatc agtcaaaata aaatcattat
     1201 ttgccatcca gctgatatcc cctatagtga gtcgtattac atggtcatag ctgtttcctg
     1261 gcagctctgg cccgtgtctc aaaatctctg atgttacatt gcacaagata aaaatatatc
     1321 atcatgaaca ataaaactgt ctgcttacat aaacagtaat acaaggggtg ttatgagcca
     1381 tattcaacgg gaaacgtcga ggccgcgatt aaattccaac atggatgctg atttatatgg
     1441 gtataaatgg gctcgcgata atgtcgggca atcaggtgcg acaatctatc gcttgtatgg
     1501 gaagcccgat gcgccagagt tgtttctgaa acatggcaaa ggtagcgttg ccaatgatgt
     1561 tacagatgag atggtcagac taaactggct gacggaattt atgcctcttc cgaccatcaa
     1621 gcattttatc cgtactcctg atgatgcatg gttactcacc actgcgatcc ccggaaaaac
     1681 agcattccag gtattagaag aatatcctga ttcaggtgaa aatattgttg atgcgctggc
     1741 agtgttcctg cgccggttgc attcgattcc tgtttgtaat tgtcctttta acagcgatcg
     1801 cgtatttcgt ctcgctcagg cgcaatcacg aatgaataac ggtttggttg atgcgagtga
     1861 ttttgatgac gagcgtaatg gctggcctgt tgaacaagtc tggaaagaaa tgcataaact
     1921 tttgccattc tcaccggatt cagtcgtcac tcatggtgat ttctcacttg ataaccttat
     1981 ttttgacgag gggaaattaa taggttgtat tgatgttgga cgagtcggaa tcgcagaccg
     2041 ataccaggat cttgccatcc tatggaactg cctcggtgag ttttctcctt cattacagaa
     2101 acggcttttt caaaaatatg gtattgataa tcctgatatg aataaattgc agtttcattt
     2161 gatgctcgat gagtttttct aatcagaatt ggttaattgg ttgtaacact ggcagagcat
     2221 tacgctgact tgacgggacg gcgcaagctc atgaccaaaa tcccttaacg tgagttacgc
     2281 gtcgttccac tgagcgtcag accccgtaga aaagatcaaa ggatcttctt gagatccttt
     2341 ttttctgcgc gtaatctgct gcttgcaaac aaaaaaacca ccgctaccag cggtggtttg
     2401 tttgccggat caagagctac caactctttt tccgaaggta actggcttca gcagagcgca
     2461 gataccaaat actgtTcttc tagtgtagcc gtagttaggc caccacttca agaactctgt
     2521 agcaccgcct acatacctcg ctctgctaat cctgttacca gtggctgctg ccagtggcga
     2581 taagtcgtgt cttaccgggt tggactcaag acgatagtta ccggataagg cgcagcggtc
     2641 gggctgaacg gggggttcgt gcacacagcc cagcttggag cgaacgacct acaccgaact
     2701 gagataccta cagcgtgagc attgagaaag cgccacgctt cccgaaggga gaaaggcgga
     2761 caggtatccg gtaagcggca gggtcggaac aggagagcgc acgagggagc ttccaggggg
     2821 aaacgcctgg tatctttata gtcctgtcgg gtttcgccac ctctgacttg agcgtcgatt
     2881 tttgtgatgc tcgtcagggg ggcggagcct atggaaaaac gccagcaacg cggccttttt
     2941 acggttcctg gccttttgct ggccttttgc tcacatgtt
//
```

**Plasmid pENTR-U6 20-4ntMM-33-C-16 (sdl 74, SPEAR) (eGFP^W58X^) sequence (GenBank format, provided as text)**

```
LOCUS       408_RP_pENTR_U6_        2930 bp ds-DNA     circular     28-APR-2022
DEFINITION  .
ACCESSION
VERSION
SOURCE      .
  ORGANISM  .
COMMENT
COMMENT     ApEinfo:methylated:1
FEATURES             Location/Qualifiers
     misc_feature    1324..2133
                     /vntifkey="21"
                     /locus_tag="KanR"
                     /ApEinfo_fwdcolor="#7eff74"
                     /ApEinfo_revcolor="#7eff74"
                     /ApEinfo_graphicformat="arrow_data {{0 1 2 0 0 -1} {} 0}
                     width 5 offset 0"
     promoter        705..968
                     /vntifkey="29"
                     /locus_tag="hU6"
                     /ApEinfo_fwdcolor="#346ee0"
                     /ApEinfo_revcolor="#346ee0"
                     /ApEinfo_graphicformat="arrow_data {{0 1 2 0 0 -1} {} 0}
                     width 5 offset 0"
     rep_origin      2264..2883
                     /vntifkey="33"
                     /locus_tag="pBR322ori"
                     /ApEinfo_fwdcolor="#999999"
                     /ApEinfo_revcolor="#999999"
                     /ApEinfo_graphicformat="arrow_data {{0 1 2 0 0 -1} {} 0}
                     width 5 offset 0"
     terminator      1045..1050
                     /locus_tag="PolIII terminator"
                     /ApEinfo_fwdcolor="#9d1b1c"
                     /ApEinfo_revcolor="#9d1b1c"
                     /ApEinfo_graphicformat="arrow_data {{0 1 2 0 0 -1} {} 0}
                     width 5 offset 0"
     misc_binding    991..994
                     /locus_tag="4nt mismatch (as in Uzonyi et al 2021)"
                     /ApEinfo_fwdcolor="#ff4809"
                     /ApEinfo_revcolor="#ff4809"
                     /ApEinfo_graphicformat="arrow_data {{0 1 2 0 0 -1} {} 0}
                     width 5 offset 0"
ORIGIN
        1 ctttcctgcg ttatcccctg attctgtgga taaccgtatt accgcctttg agtgagctga
       61 taccgctcgc cgcagccgaa cgaccgagcg cagcgagtca gtgagcgagg aagcggaaga
      121 gcgcccaata cgcaaaccgc ctctccccgc gcgttggccg attcattaat gcagctggca
      181 cgacaggttt cccgactgga aagcgggcag tgagcgcaac gcaattaata cgcgtaccgc
      241 tagccaggaa gagtttgtag aaacgcaaaa aggccatccg tcaggatggc cttctgctta
      301 gtttgatgcc tggcagttta tggcgggcgt cctgcccgcc accctccggg ccgttgcttc
      361 acaacgttca aatccgctcc cggcggattt gtcctactca ggagagcgtt caccgacaaa
      421 caacagataa aacgaaaggc ccagtcttcc gactgagcct ttcgttttat ttgatgcctg
      481 gcagttccct actctcgcgt taacgctagc atggatgttt tcccagtcac gacgttgtaa
      541 aacgacggcc agtcttaagc tcgggcccca aataatgatt ttattttgac tgatagtgac
      601 ctgttcgttg caacaaattg atgagcaatg cttttttata atgccaactt tgtacaaaaa
      661 agcaggcttt aaaggaacca attcagtcga ctggatccgg taccaaggtc gggcaggaag
      721 agggcctatt tcccatgatt ccttcatatt tgcatatacg atacaaggct gttagagaga
      781 taattagaat taatttgact gtaaacacaa agatattagt acaaaatacg tgacgtagaa
      841 agtaataatt tcttgggtag tttgcagttt taaaattatg ttttaaaatg gactatcata
      901 tgcttaccgt aacttgaaag tatttcgatt tcttggcttt atatatcttg tggaaaggac
      961 gaaacaccGG tcggggtacc tgctgaagca aacgacgccg tatgtcaggg tagtgacaag
     1021 tgttggcCat ggaacaggta gttttttttt ctagacccag ctttcttgta caaagttggc
     1081 attataagaa agcattgctt atcaatttgt tgcaacgaac aggtcactat cagtcaaaat
     1141 aaaatcatta tttgccatcc agctgatatc ccctatagtg agtcgtatta catggtcata
     1201 gctgtttcct ggcagctctg gcccgtgtct caaaatctct gatgttacat tgcacaagat
     1261 aaaaatatat catcatgaac aataaaactg tctgcttaca taaacagtaa tacaaggggt
     1321 gttatgagcc atattcaacg ggaaacgtcg aggccgcgat taaattccaa catggatgct
     1381 gatttatatg ggtataaatg ggctcgcgat aatgtcgggc aatcaggtgc gacaatctat
     1441 cgcttgtatg ggaagcccga tgcgccagag ttgtttctga aacatggcaa aggtagcgtt
     1501 gccaatgatg ttacagatga gatggtcaga ctaaactggc tgacggaatt tatgcctctt
     1561 ccgaccatca agcattttat ccgtactcct gatgatgcat ggttactcac cactgcgatc
     1621 cccggaaaaa cagcattcca ggtattagaa gaatatcctg attcaggtga aaatattgtt
     1681 gatgcgctgg cagtgttcct gcgccggttg cattcgattc ctgtttgtaa ttgtcctttt
     1741 aacagcgatc gcgtatttcg tctcgctcag gcgcaatcac gaatgaataa cggtttggtt
     1801 gatgcgagtg attttgatga cgagcgtaat ggctggcctg ttgaacaagt ctggaaagaa
     1861 atgcataaac ttttgccatt ctcaccggat tcagtcgtca ctcatggtga tttctcactt
     1921 gataacctta tttttgacga ggggaaatta ataggttgta ttgatgttgg acgagtcgga
     1981 atcgcagacc gataccagga tcttgccatc ctatggaact gcctcggtga gttttctcct
     2041 tcattacaga aacggctttt tcaaaaatat ggtattgata atcctgatat gaataaattg
     2101 cagtttcatt tgatgctcga tgagtttttc taatcagaat tggttaattg gttgtaacac
     2161 tggcagagca ttacgctgac ttgacgggac ggcgcaagct catgaccaaa atcccttaac
     2221 gtgagttacg cgtcgttcca ctgagcgtca gaccccgtag aaaagatcaa aggatcttct
     2281 tgagatcctt tttttctgcg cgtaatctgc tgcttgcaaa caaaaaaacc accgctacca
     2341 gcggtggttt gtttgccgga tcaagagcta ccaactcttt ttccgaaggt aactggcttc
     2401 agcagagcgc agataccaaa tactgttctt ctagtgtagc cgtagttagg ccaccacttc
     2461 aagaactctg tagcaccgcc tacatacctc gctctgctaa tcctgttacc agtggctgct
     2521 gccagtggcg ataagtcgtg tcttaccggg ttggactcaa gacgatagtt accggataag
     2581 gcgcagcggt cgggctgaac ggggggttcg tgcacacagc ccagcttgga gcgaacgacc
     2641 tacaccgaac tgagatacct acagcgtgag cattgagaaa gcgccacgct tcccgaaggg
     2701 agaaaggcgg acaggtatcc ggtaagcggc agggtcggaa caggagagcg cacgagggag
     2761 cttccagggg gaaacgcctg gtatctttat agtcctgtcg ggtttcgcca cctctgactt
     2821 gagcgtcgat ttttgtgatg ctcgtcaggg gggcggagcc tatggaaaaa cgccagcaac
     2881 gcggcctttt tacggttcct ggccttttgc tggccttttg ctcacatgtt
//
```

**Plasmid hU6 NT-SPEAR (control)_hPGK coMART1^WT^-T2A-BFP-P2A-Bsd sequence (GenBank format, provided as text)**

```
LOCUS       RP468_pLV_hU6_AS        8620 bp ds-DNA     circular     28-JAN-2023
DEFINITION
ACCESSION   VB210525-1302qyy
VERSION     VB210525-1302qyy
KEYWORDS    .
SOURCE
  ORGANISM  .
COMMENT
COMMENT
COMMENT     ApEinfo:methylated:1
FEATURES             Location/Qualifiers
     Promoter        1..229
                     /__SeqFeature__="True"
                     /__level__="0"
                     /application_notes="Strong promoter; drives transcription
                     of viral RNA in packaging cells."
                     /description="['Rous sarcoma virus enhancer/promoter']"
                     /full_name="Rous sarcoma virus (RSV) enhancer/promoter"
                     /name="RSV  promoter"
                     /note="color: #946a00; direction: RIGHT"
                     /official_designation="RSV"
                     /uuid="3f2ddca7f0cb4015b6abcec8a25c834e"
                     /vntifkey="21"
                     /locus_tag="RSV  promoter"
                     /ApEinfo_fwdcolor="pink"
                     /ApEinfo_revcolor="pink"
                     /ApEinfo_graphicformat="arrow_data {{0 1 2 0 0 -1} {} 0}
                     width 5 offset 0"
     promoter        1927..2190
                     /vntifkey="29"
                     /locus_tag="hU6"
                     /ApEinfo_fwdcolor="#346ee0"
                     /ApEinfo_revcolor="#346ee0"
                     /ApEinfo_graphicformat="arrow_data {{0 1 2 0 0 -1} {} 0}
                     width 5 offset 0"
     LTR             230..410
                     /application_notes="Allows transcription of viral RNA and
                     its packaging into virus."
                     /description="[Truncated HIV-1 5' long terminal repeat]"
                     /full_name="HIV-1 truncated 5' LTR"
                     /note="color: #512bbd; direction: RIGHT"
                     /official_designation="Delta5' LTR"
                     /uuid="44156a3083604321be44066133e8ecb9"
                     /locus_tag="5' LTR"
                     /ApEinfo_fwdcolor="#86f71d"
                     /ApEinfo_revcolor="#d7336f"
                     /ApEinfo_graphicformat="arrow_data {{0 1 2 0 0 -1} {} 0}
                     width 5 offset 0"
     Miscellaneous   521..565
                     /application_notes="Allows packaging of viral RNA into
                     virus."
                     /description="['HIV-1 packaging signal ']"
                     /full_name="HIV-1 psi packaging signal"
                     /note="color: #d84e4e; direction: RIGHT"
                     /official_designation="Psi"
                     /uuid="284899413ff943a99f8210a11ad8c54e"
                     /locus_tag="ÃÂÃÂÃÂÃÂ¨"
                     /ApEinfo_fwdcolor="pink"
                     /ApEinfo_revcolor="pink"
                     /ApEinfo_graphicformat="arrow_data {{0 1 2 0 0 -1} {} 0}
                     width 5 offset 0"
     misc_binding    2213..2216
                     /locus_tag="4nt mismatch (as in Uzonyi et al 2021)"
                     /ApEinfo_fwdcolor="#ff4809"
                     /ApEinfo_revcolor="#ff4809"
                     /ApEinfo_graphicformat="arrow_data {{0 1 2 0 0 -1} {} 0}
                     width 5 offset 0"
     terminator      2267..2272
                     /locus_tag="PolIII terminator"
                     /ApEinfo_fwdcolor="#9d1b1c"
                     /ApEinfo_revcolor="#9d1b1c"
                     /ApEinfo_graphicformat="arrow_data {{0 1 2 0 0 -1} {} 0}
                     width 5 offset 0"
     Miscellaneous   1075..1308
                     /__SeqFeature__="True"
                     /__level__="0"
                     /application_notes="Rev protein binding site that allows
                     Rev-dependent nuclear export of viral RNA during viral
                     packaging."
                     /description="['HIV-1 Rev response element']"
                     /full_name="HIV-1 Rev response element"
                     /name="RRE"
                     /note="color: #1f36a9; direction: RIGHT"
                     /official_designation="RRE"
                     /uuid="df07099e49674cf3aba3661b54e28bd7"
                     /vntifkey="21"
                     /locus_tag="RRE"
                     /ApEinfo_fwdcolor="pink"
                     /ApEinfo_revcolor="pink"
                     /ApEinfo_graphicformat="arrow_data {{0 1 2 0 0 -1} {} 0}
                     width 5 offset 0"
     ORF             3267..3980
                     /application_notes="Low fluorescence and low
                     photostability."
                     /description="Blue variant of EGFP generated by
                     mutagenesis"
                     /full_name="Enhanced blue fluorescent protein(ns)"
                     /note="color: #6ddaae; direction: RIGHT"
                     /official_designation="EBFP(ns)"
                     /uuid="c22d3886879b4745b895a6f25c150792"
                     /locus_tag="EBFP(ns)"
                     /ApEinfo_fwdcolor="#0f7ffe"
                     /ApEinfo_revcolor="pink"
                     /ApEinfo_graphicformat="arrow_data {{0 1 2 0 0 -1} {} 0}
                     width 5 offset 0"
     Miscellaneous   1803..1920
                     /__SeqFeature__="True"
                     /__level__="0"
                     /application_notes="Facilitates the nuclear import of
                     HIV-1 cDNA through a central DNA flap."
                     /description="['Central polypurine tract']"
                     /full_name="Central polypurine tract"
                     /name="cPPT"
                     /note="color: #db6901; direction: RIGHT"
                     /official_designation="cPPT"
                     /uuid="f7cb29985f9f41a7a5924d3462266f42"
                     /vntifkey="21"
                     /locus_tag="cPPT"
                     /ApEinfo_fwdcolor="pink"
                     /ApEinfo_revcolor="pink"
                     /ApEinfo_graphicformat="arrow_data {{0 1 2 0 0 -1} {} 0}
                     width 5 offset 0"
     Linker          3981..4046
                     /application_notes="Causes co-translational cleavage of
                     the encoded polypeptide. Multiple proteins can be made
                     from a polycistronic transcript containing multiple ORFs
                     separated by 2A. P2A and T2A have higher cleavage
                     efficiency compared to other 2As."
                     /description="Self-cleaving 2A peptide from Porcine
                     teschovirus-1"
                     /full_name="Porcine teschovirus-1 2A peptide"
                     /note="color: #1f36a9; direction: RIGHT"
                     /official_designation="P2A"
                     /uuid="26656daa74294a0f8b331614d72d5f5a"
                     /locus_tag="P2A"
                     /ApEinfo_fwdcolor="#20fefe"
                     /ApEinfo_revcolor="pink"
                     /ApEinfo_graphicformat="arrow_data {{0 1 2 0 0 -1} {} 0}
                     width 5 offset 0"
     promoter        2300..2804
                     /__SeqFeature__="True"
                     /__level__="0"
                     /application_notes="Medium-strength promoter."
                     /description="['Human phosphoglycerate kinase 1
                     promoter']"
                     /name="hPGK promoter"
                     /note="color: #5566f5; direction: RIGHT"
                     /official_designation="hPGK promoter"
                     /uuid="a105ee8f48814e769c9a41c38cfff41e"
                     /vntifkey="21"
                     /locus_tag="hPGK promoter"
                     /ApEinfo_fwdcolor="#346ee0"
                     /ApEinfo_revcolor="#346ee0"
                     /ApEinfo_graphicformat="arrow_data {{0 1 2 0 0 -1} {} 0}
                     width 5 offset 0"
     CDS             4047..4445
                     /__SeqFeature__="True"
                     /__level__="0"
                     /application_notes="Allows cells to be resistant to
                     blasticidin."
                     /description="['Blasticidin resistance gene']"
                     /full_name="Blasticidin resistance gene"
                     /name="Bsd"
                     /note="color: #68e66d; direction: RIGHT"
                     /official_designation="Bsd"
                     /uuid="70e7f83c244f451dae86de92afa03717"
                     /vntifkey="21"
                     /locus_tag="Bsd"
                     /ApEinfo_fwdcolor="#e9d024"
                     /ApEinfo_revcolor="#e9d024"
                     /ApEinfo_graphicformat="arrow_data {{0 1 2 0 0 -1} {} 0}
                     width 5 offset 0"
     misc_signal     4477..5074
                     /__SeqFeature__="True"
                     /__level__="0"
                     /application_notes="Enhances virus stability in packaging
                     cells, leading to higher titer of packaged virus; enhances
                     higher expression of transgenes."
                     /description="['Woodchuck hepatitis virus
                     posttranscriptional regulatory element']"
                     /full_name="Woodchuck hepatitis virus posttranscriptional
                     regulatory element"
                     /name="WPRE"
                     /note="color: #ef6cdf; direction: RIGHT"
                     /official_designation="WPRE"
                     /uuid="0aedff8308474b369c622433538d7836"
                     /vntifkey="21"
                     /locus_tag="WPRE"
                     /ApEinfo_fwdcolor="#008040"
                     /ApEinfo_revcolor="#008040"
                     /ApEinfo_graphicformat="arrow_data {{0 1 2 0 0 -1} {} 0}
                     width 5 offset 0"
     LTR             5140..5374
                     /application_notes="Allows packaging of viral RNA into
                     virus; self-inactivates the 5' LTR by a copying mechanism
                     during viral genome integration; contains polyadenylation
                     signal for transcription termination."
                     /description="[Truncated HIV-1 3' long terminal repeat ]"
                     /full_name="HIV-1 truncated 3' LTR"
                     /note="color: #c54b7c; direction: RIGHT"
                     /official_designation="DeltaU3/3' LTR"
                     /uuid="c958755414c34372b909e20e81ebcef6"
                     /locus_tag="U3/3' LTR"
                     /ApEinfo_fwdcolor="#86f71d"
                     /ApEinfo_revcolor="#d7336f"
                     /ApEinfo_graphicformat="arrow_data {{0 1 2 0 0 -1} {} 0}
                     width 5 offset 0"
     polyA_signal    5447..5581
                     /__SeqFeature__="True"
                     /__level__="0"
                     /application_notes="Allows transcription termination and
                     polyadenylation of mRNA transcribed by Pol II RNA
                     polymerase."
                     /description="['Simian virus 40 early polyadenylation
                     signal']"
                     /full_name="SV40 early polyadenation signal"
                     /name="SV40 early pA"
                     /note="color: #d05c0a; direction: RIGHT"
                     /official_designation="SV40 early pA"
                     /uuid="2fcb13a62e704d8095b188d9625a6a94"
                     /vntifkey="21"
                     /locus_tag="SV40 early pA"
                     /ApEinfo_fwdcolor="#ff3eee"
                     /ApEinfo_revcolor="#ff3eee"
                     /ApEinfo_graphicformat="arrow_data {{0 1 2 0 0 -1} {} 0}
                     width 5 offset 0"
     ORF             6535..7395
                     /__SeqFeature__="True"
                     /__level__="0"
                     /application_notes="Allows E. coli to be resistant to
                     ampicillin."
                     /description="['AmpiciIIin resistance gene']"
                     /full_name="Ampicillin resistance gene"
                     /name="AmpiciIIin"
                     /note="color: #6ddaae; direction: RIGHT"
                     /official_designation="Ampicillin"
                     /uuid="6a7994dc58024d80b36e4a85ad2e1d99"
                     /vntifkey="21"
                     /locus_tag="AmpiciIIin"
                     /ApEinfo_fwdcolor="pink"
                     /ApEinfo_revcolor="pink"
                     /ApEinfo_graphicformat="arrow_data {{0 1 2 0 0 -1} {} 0}
                     width 5 offset 0"
     Rep_origin      7566..8154
                     /__SeqFeature__="True"
                     /__level__="0"
                     /application_notes="Facilitates plasmid replication in E.
                     coli; regulates high-copy plasmid number (500-700)."
                     /description="['pUC origin of replication']"
                     /full_name="pUC origin of replication"
                     /name="pUC ori"
                     /note="color: #fd3434; direction: RIGHT"
                     /official_designation="pUC ori"
                     /uuid="64a8cb3def9f49deb6752795c2d893a8"
                     /vntifkey="21"
                     /locus_tag="pUC ori"
                     /ApEinfo_fwdcolor="pink"
                     /ApEinfo_revcolor="pink"
                     /ApEinfo_graphicformat="arrow_data {{0 1 2 0 0 -1} {} 0}
                     width 5 offset 0"
     primer_bind     8578..8597
                     /locus_tag="T3"
                     /ApEinfo_fwdcolor="cyan"
                     /ApEinfo_revcolor="green"
                     /ApEinfo_graphicformat="arrow_data {{0 1 2 0 0 -1} {} 0}
                     width 5 offset 0"
     primer_bind     complement(5791..5808)
                     /locus_tag="M13-fwd"
                     /ApEinfo_fwdcolor="cyan"
                     /ApEinfo_revcolor="green"
                     /ApEinfo_graphicformat="arrow_data {{0 1 2 0 0 -1} {} 0}
                     width 5 offset 0"
     primer_bind     8540..8560
                     /locus_tag="M13-rev"
                     /ApEinfo_fwdcolor="cyan"
                     /ApEinfo_revcolor="green"
                     /ApEinfo_graphicformat="arrow_data {{0 1 2 0 0 -1} {} 0}
                     width 5 offset 0"
     primer_bind     complement(5762..5782)
                     /locus_tag="T7"
                     /ApEinfo_fwdcolor="cyan"
                     /ApEinfo_revcolor="green"
                     /ApEinfo_graphicformat="arrow_data {{0 1 2 0 0 -1} {} 0}
                     width 5 offset 0"
     rep_origin      5966..6272
                     /locus_tag="F1 ori"
                     /ApEinfo_fwdcolor="gray50"
                     /ApEinfo_revcolor="gray50"
                     /ApEinfo_graphicformat="arrow_data {{0 1 2 0 0 -1} {} 0}
                     width 5 offset 0"
     CDS             5879..5947
                     /locus_tag="LacZ alpha"
                     /ApEinfo_fwdcolor="#6495ed"
                     /ApEinfo_revcolor="#6495ed"
                     /ApEinfo_graphicformat="arrow_data {{0 1 2 0 0 -1} {} 0}
                     width 5 offset 0"
     misc_binding    8512..8534
                     /locus_tag="LacO"
                     /ApEinfo_fwdcolor="#6495ed"
                     /ApEinfo_revcolor="#6495ed"
                     /ApEinfo_graphicformat="arrow_data {{0 1 2 0 0 -1} {} 0}
                     width 5 offset 0"
     CDS             6733..7392
                     /locus_tag="AmpR"
                     /ApEinfo_fwdcolor="yellow"
                     /ApEinfo_revcolor="yellow"
                     /ApEinfo_graphicformat="arrow_data {{0 1 2 0 0 -1} {} 0}
                     width 5 offset 0"
     CDS             2835..3191
                     /standard_name="MLANA_cds"
                     /locus_tag="MLANA_cds"
                     /ApEinfo_fwdcolor="#e9d024"
                     /ApEinfo_revcolor="#e9d024"
                     /ApEinfo_graphicformat="arrow_data {{0 1 2 0 0 -1} {} 0}
                     width 5 offset 0"
     sig_peptide     3213..3266
                     /standard_name="T2A"
                     /note="Geneious type: LINKER"
                     /locus_tag="T2A"
                     /ApEinfo_fwdcolor="cyan"
                     /ApEinfo_revcolor="green"
                     /ApEinfo_graphicformat="arrow_data {{0 1 2 0 0 -1} {} 0}
                     width 5 offset 0"
     misc_feature    2910..2939
                     /standard_name="MART-1_26-35 G6S"
                     /locus_tag="MART-1_26-35"
                     /ApEinfo_fwdcolor="#ff7d78"
                     /ApEinfo_revcolor="#7eff74"
                     /ApEinfo_graphicformat="arrow_data {{0 1 2 0 0 -1} {} 0}
                     width 5 offset 0"
     misc_feature    2925..2927
                     /standard_name="G6S"
                     /locus_tag="G31"
                     /ApEinfo_fwdcolor="#941100"
                     /ApEinfo_revcolor="#7eff74"
                     /ApEinfo_graphicformat="arrow_data {{0 1 2 0 0 -1} {} 0}
                     width 5 offset 0"
     misc_feature    3192..3212
                     /locus_tag="linker"
                     /ApEinfo_fwdcolor="#a8eafe"
                     /ApEinfo_revcolor="green"
                     /ApEinfo_graphicformat="arrow_data {{0 1 2 0 0 -1} {} 0}
                     width 5 offset 0"
ORIGIN
        1 aatgtagtct tatgcaatac tcttgtagtc ttgcaacatg gtaacgatga gttagcaaca
       61 tgccttacaa ggagagaaaa agcaccgtgc atgccgattg gtggaagtaa ggtggtacga
      121 tcgtgcctta ttaggaaggc aacagacggg tctgacatgg attggacgaa ccactgaatt
      181 gccgcattgc agagatattg tatttaagtg cctagctcga tacataaacg ggtctctctg
      241 gttagaccag atctgagcct gggagctctc tggctaacta gggaacccac tgcttaagcc
      301 tcaataaagc ttgccttgag tgcttcaagt agtgtgtgcc cgtctgttgt gtgactctgg
      361 taactagaga tccctcagac ccttttagtc agtgtggaaa atctctagca gtggcgcccg
      421 aacagggact tgaaagcgaa agggaaacca gaggagctct ctcgacgcag gactcggctt
      481 gctgaagcgc gcacggcaag aggcgagggg cggcgactgg tgagtacgcc aaaaattttg
      541 actagcggag gctagaagga gagagatggg tgcgagagcg tcagtattaa gcgggggaga
      601 attagatcgc gatgggaaaa aattcggtta aggccagggg gaaagaaaaa atataaatta
      661 aaacatatag tatgggcaag cagggagcta gaacgattcg cagttaatcc tggcctgtta
      721 gaaacatcag aaggctgtag acaaatactg ggacagctac aaccatccct tcagacagga
      781 tcagaagaac ttagatcatt atataataca gtagcaaccc tctattgtgt gcatcaaagg
      841 atagagataa aagacaccaa ggaagcttta gacaagatag aggaagagca aaacaaaagt
      901 aagaccaccg cacagcaagc ggccgctgat cttcagacct ggaggaggag atatgaggga
      961 caattggaga agtgaattat ataaatataa agtagtaaaa attgaaccat taggagtagc
     1021 acccaccaag gcaaagagaa gagtggtgca gagagaaaaa agagcagtgg gaataggagc
     1081 tttgttcctt gggttcttgg gagcagcagg aagcactatg ggcgcagcgt caatgacgct
     1141 gacggtacag gccagacaat tattgtctgg tatagtgcag cagcagaaca atttgctgag
     1201 ggctattgag gcgcaacagc atctgttgca actcacagtc tggggcatca agcagctcca
     1261 ggcaagaatc ctggctgtgg aaagatacct aaaggatcaa cagctcctgg ggatttgggg
     1321 ttgctctgga aaactcattt gcaccactgc tgtgccttgg aatgctagtt ggagtaataa
     1381 atctctggaa cagatttgga atcacacgac ctggatggag tgggacagag aaattaacaa
     1441 ttacacaagc ttaatacact ccttaattga agaatcgcaa aaccagcaag aaaagaatga
     1501 acaagaatta ttggaattag ataaatgggc aagtttgtgg aattggttta acataacaaa
     1561 ttggctgtgg tatataaaat tattcataat gatagtagga ggcttggtag gtttaagaat
     1621 agtttttgct gtactttcta tagtgaatag agttaggcag ggatattcac cattatcgtt
     1681 tcagacccac ctcccaaccc cgaggggacc cgacaggccc gaaggaatag aagaagaagg
     1741 tggagagaga gacagagaca gatccattcg attagtgaac ggatctcgac ggtatcgcta
     1801 gcttttaaaa gaaaaggggg gattgggggg tacagtgcag gggaaagaat agtagacata
     1861 atagcaacag acatacaaac taaagaatta caaaaacaaa ttacaaaaat tcaaaatttt
     1921 actagtaagg tcgggcagga agagggccta tttcccatga ttccttcata tttgcatata
     1981 cgatacaagg ctgttagaga gataattaga attaatttga ctgtaaacac aaagatatta
     2041 gtacaaaata cgtgacgtag aaagtaataa tttcttgggt agtttgcagt tttaaaatta
     2101 tgttttaaaa tggactatca tatgcttacc gtaacttgaa agtatttcga tttcttggct
     2161 ttatatatct tgtggaaagg acgaaacacc GGtcggggta cctgctgaag caaacgacgc
     2221 cgtatgtcag ggtagtgaca agtgttggcC atggaacagg tagttttttt ttgaattcca
     2281 actttgtata gaaaagttgg ggttgcgcct tttccaaggc agccctgggt ttgcgcaggg
     2341 acgcggctgc tctgggcgtg gttccgggaa acgcagcggc gccgaccctg ggtctcgcac
     2401 attcttcacg tccgttcgca gcgtcacccg gatcttcgcc gctacccttg tgggcccccc
     2461 ggcgacgctt cctgctccgc ccctaagtcg ggaaggttcc ttgcggttcg cggcgtgccg
     2521 gacgtgacaa acggaagccg cacgtctcac tagtaccctc gcagacggac agcgccaggg
     2581 agcaatggca gcgcgccgac cgcgatgggc tgtggccaat agcggctgct cagcagggcg
     2641 cgccgagagc agcggccggg aaggggcggt gcgggaggcg gggtgtgggg cggtagtgtg
     2701 ggccctgttc ctgcccgcgc ggtgttccgc attctgcaag cctccggagc gcacgtcggc
     2761 agtcggctcc ctcgttgacc gaatcaccga cctctctccc caggcaagtt tgtacaaaaa
     2821 agcaggctgc caccatgccc agggaggacg cccacttcat ctacggctac cccaagaagg
     2881 gccacggcca cagctacacc actgcagagg agcttgctgg gatcggcatc ctgacagtga
     2941 tcctaggcgt gctgctgctg atcggctgct ggtactgccg gaggaggaac ggctacaggg
     3001 ccctgatgga caagagcctg cacgtgggca cccagtgcgc cctgaccagg aggtgccccc
     3061 aggagggctt cgaccacagg gacagcaagg tgagcctcca ggagaagaac tgcgagcccg
     3121 tggtgcccaa cgcccccccc gcctacgaga agctgagcgc cgagcagagc cctccaccat
     3181 acagccccgg tggatccgga ggtgcgagcg gcgaggggag gggcagcctt cttacttgcg
     3241 gagatgtgga agagaatcca ggacccgtga gcaagggcga ggagctgttc accggggtgg
     3301 tgcccatcct ggtcgagctg gacggcgacg taaacggcca caagttcagc gtgtccggcg
     3361 agggcgaggg cgatgccacc tacggcaagc tgaccctgaa gttcatctgc accaccggca
     3421 agctgcccgt gccctggccc accctcgtga ccaccctgac ccacggcgtg cagtgcttca
     3481 gccgctaccc cgaccacatg aagcagcacg acttcttcaa gtccgccatg cccgaaggct
     3541 acgtccagga gcgcaccatc ttcttcaagg acgacggcaa ctacaagacc cgcgccgagg
     3601 tgaagttcga gggcgacacc ctggtgaacc gcatcgagct gaagggcatc gacttcaagg
     3661 aggacggcaa catcctgggg cacaagctgg agtacaactt caacagccac aacgtctata
     3721 tcatggccga caagcagaag aacggcatca aggtgaactt caagatccgc cacaacatcg
     3781 aggacggcag cgtgcagctc gccgaccact accagcagaa cacccccatc ggcgacggcc
     3841 ccgtgctgct gcccgacaac cactacctga gcacccagtc cgccctgagc aaagacccca
     3901 acgagaagcg cgatcacatg gtcctgctgg agttcgtgac cgccgccggg atcactctcg
     3961 gcatggacga gctgtacaag ggaagcggag ccacgaactt ctctctgtta aagcaagcag
     4021 gagatgttga agaaaacccc gggcctatgg ccaagccttt gtctcaagaa gaatccaccc
     4081 tcattgaaag agcaacggct acaatcaaca gcatccccat ctctgaagac tacagcgtcg
     4141 ccagcgcagc tctctctagc gacggccgca tcttcactgg tgtcaatgta tatcatttta
     4201 ctgggggacc ttgtgcagaa ctcgtggtgc tgggcactgc tgctgctgcg gcagctggca
     4261 acctgacttg tatcgtcgcg atcggaaatg agaacagggg catcttgagc ccctgcggac
     4321 ggtgccgaca ggtgcttctc gatctgcatc ctgggatcaa agccatagtg aaggacagtg
     4381 atggacagcc gacggcagtt gggattcgtg aattgctgcc ctctggttat gtgtgggagg
     4441 gctaaaccca gctttcttgt acaaagtggt ggtacccgat aatcaacctc tggattacaa
     4501 aatttgtgaa agattgactg gtattcttaa ctatgttgct ccttttacgc tatgtggata
     4561 cgctgcttta atgcctttgt atcatgctat tgcttcccgt atggctttca ttttctcctc
     4621 cttgtataaa tcctggttgc tgtctcttta tgaggagttg tggcccgttg tcaggcaacg
     4681 tggcgtggtg tgcactgtgt ttgctgacgc aacccccact ggttggggca ttgccaccac
     4741 ctgtcagctc ctttccggga ctttcgcttt ccccctccct attgccacgg cggaactcat
     4801 cgccgcctgc cttgcccgct gctggacagg ggctcggctg ttgggcactg acaattccgt
     4861 ggtgttgtcg gggaagctga cgtcctttcc atggctgctc gcctgtgttg ccacctggat
     4921 tctgcgcggg acgtccttct gctacgtccc ttcggccctc aatccagcgg accttccttc
     4981 ccgcggcctg ctgccggctc tgcggcctct tccgcgtctt cgccttcgcc ctcagacgag
     5041 tcggatctcc ctttgggccg cctccccgca tcggctttaa gaccaatgac ttacaaggca
     5101 gctgtagatc ttagccactt tttaaaagaa aaggggggac tggaagggct aattcactcc
     5161 caacgaagac aagatctgct ttttgcttgt actgggtctc tctggttaga ccagatctga
     5221 gcctgggagc tctctggcta actagggaac ccactgctta agcctcaata aagcttgcct
     5281 tgagtgcttc aagtagtgtg tgcccgtctg ttgtgtgact ctggtaacta gagatccctc
     5341 agaccctttt agtcagtgtg gaaaatctct agcagtagta gttcatgtca tcttattatt
     5401 cagtatttat aacttgcaaa gaaatgaata tcagagagtg agaggaactt gtttattgca
     5461 gcttataatg gttacaaata aagcaatagc atcacaaatt tcacaaataa agcatttttt
     5521 tcactgcatt ctagttgtgg tttgtccaaa ctcatcaatg tatcttatca tgtctggctc
     5581 tagctatccc gcccctaact ccgcccatcc cgcccctaac tccgcccagt tccgcccatt
     5641 ctccgcccca tggctgacta atttttttta tttatgcaga ggccgaggcc gcctcggcct
     5701 ctgagctatt ccagaagtag tgaggaggct tttttggagg cctagggacg tacccaattc
     5761 gccctatagt gagtcgtatt acgcgcgctc actggccgtc gttttacaac gtcgtgactg
     5821 ggaaaaccct ggcgttaccc aacttaatcg ccttgcagca catccccctt tcgccagctg
     5881 gcgtaatagc gaagaggccc gcaccgatcg cccttcccaa cagttgcgca gcctgaatgg
     5941 cgaatgggac gcgccctgta gcggcgcatt aagcgcggcg ggtgtggtgg ttacgcgcag
     6001 cgtgaccgct acacttgcca gcgccctagc gcccgctcct ttcgctttct tcccttcctt
     6061 tctcgccacg ttcgccggct ttccccgtca agctctaaat cgggggctcc ctttagggtt
     6121 ccgatttagt gctttacggc acctcgaccc caaaaaactt gattagggtg atggttcacg
     6181 tagtgggcca tcgccctgat agacggtttt tcgccctttg acgttggagt ccacgttctt
     6241 taatagtgga ctcttgttcc aaactggaac aacactcaac cctatctcgg tctattcttt
     6301 tgatttataa gggattttgc cgatttcggc ctattggtta aaaaatgagc tgatttaaca
     6361 aaaatttaac gcgaatttta acaaaatatt aacgcttaca atttaggtgg cacttttcgg
     6421 ggaaatgtgc gcggaacccc tatttgttta tttttctaaa tacattcaaa tatgtatccg
     6481 ctcatgagac aataaccctg ataaatgctt caataatatt gaaaaaggaa gagtatgagt
     6541 attcaacatt tccgtgtcgc ccttattccc ttttttgcgg cattttgcct tcctgttttt
     6601 gctcacccag aaacgctggt gaaagtaaaa gatgctgaag atcagttggg tgcacgagtg
     6661 ggttacatcg aactggatct caacagcggt aagatccttg agagttttcg ccccgaagaa
     6721 cgttttccaa tgatgagcac ttttaaagtt ctgctatgtg gcgcggtatt atcccgtatt
     6781 gacgccgggc aagagcaact cggtcgccgc atacactatt ctcagaatga cttggttgag
     6841 tactcaccag tcacagaaaa gcatcttacg gatggcatga cagtaagaga attatgcagt
     6901 gctgccataa ccatgagtga taacactgcg gccaacttac ttctgacaac gatcggagga
     6961 ccgaaggagc taaccgcttt tttgcacaac atgggggatc atgtaactcg ccttgatcgt
     7021 tgggaaccgg agctgaatga agccatacca aacgacgagc gtgacaccac gatgcctgta
     7081 gcaatggcaa caacgttgcg caaactatta actggcgaac tacttactct agcttcccgg
     7141 caacaattaa tagactggat ggaggcggat aaagttgcag gaccacttct gcgctcggcc
     7201 cttccggctg gctggtttat tgctgataaa tctggagccg gtgagcgtgg gtctcgcggt
     7261 atcattgcag cactggggcc agatggtaag ccctcccgta tcgtagttat ctacacgacg
     7321 gggagtcagg caactatgga tgaacgaaat agacagatcg ctgagatagg tgcctcactg
     7381 attaagcatt ggtaactgtc agaccaagtt tactcatata tactttagat tgatttaaaa
     7441 cttcattttt aatttaaaag gatctaggtg aagatccttt ttgataatct catgaccaaa
     7501 atcccttaac gtgagttttc gttccactga gcgtcagacc ccgtagaaaa gatcaaagga
     7561 tcttcttgag atcctttttt tctgcgcgta atctgctgct tgcaaacaaa aaaaccaccg
     7621 ctaccagcgg tggtttgttt gccggatcaa gagctaccaa ctctttttcc gaaggtaact
     7681 ggcttcagca gagcgcagat accaaatact gttcttctag tgtagccgta gttaggccac
     7741 cacttcaaga actctgtagc accgcctaca tacctcgctc tgctaatcct gttaccagtg
     7801 gctgctgcca gtggcgataa gtcgtgtctt accgggttgg actcaagacg atagttaccg
     7861 gataaggcgc agcggtcggg ctgaacgggg ggttcgtgca cacagcccag cttggagcga
     7921 acgacctaca ccgaactgag atacctacag cgtgagctat gagaaagcgc cacgcttccc
     7981 gaagagagaa aggcggacag gtatccggta agcggcaggg tcggaacagg agagcgcacg
     8041 agggagcttc cagggggaaa cgcctggtat ctttatagtc ctgtcgggtt tcgccacctc
     8101 tgacttgagc gtcgattttt gtgatgctcg tcaggggggc ggagcctatg gaaaaacgcc
     8161 agcaacgcgg cctttttacg gttcctggcc ttttgctggc cttttgctca catgttcttt
     8221 cctgcgttat cccctgattc tgtggataac cgtattaccg cctttgagtg agctgatacc
     8281 gctcgccgca gccgaacgac cgagcgcagc gagtcagtga gcgaggaagc ggaagagcgc
     8341 ccaatacgca aaccgcctct ccccgcgcgt tggccgattc attaatgcag ctggcacgac
     8401 aggtttcccg actggaaagc gggcagtgag cgcaacgcaa ttaatgtgag ttagctcact
     8461 cattaggcac cccaggcttt acactttatg cttccggctc gtatgttgtg tggaattgtg
     8521 agcggataac aatttcacac aggaaacagc tatgaccatg attacgccaa gcgcgcaatt
     8581 aaccctcact aaagggaaca aaagctggag ctgcaagctt
//
```

**Plasmid hU6 NT-SPEAR (control)_hPGK coMART1^G31S^-T2A-BFP-P2A-Bsd sequence (GenBank format, provided as text)**

```
LOCUS       RP469_pLV_hU6_AS        8620 bp ds-DNA     circular     04-JAN-2023
DEFINITION
ACCESSION   VB210525-1302qyy
VERSION     VB210525-1302qyy
KEYWORDS    .
SOURCE
  ORGANISM  .
COMMENT
COMMENT
COMMENT     ApEinfo:methylated:1
FEATURES             Location/Qualifiers
     Promoter        1..229
                     /__SeqFeature__="True"
                     /__level__="0"
                     /application_notes="Strong promoter; drives transcription
                     of viral RNA in packaging cells."
                     /description="['Rous sarcoma virus enhancer/promoter']"
                     /full_name="Rous sarcoma virus (RSV) enhancer/promoter"
                     /name="RSV  promoter"
                     /note="color: #946a00; direction: RIGHT"
                     /official_designation="RSV"
                     /uuid="3f2ddca7f0cb4015b6abcec8a25c834e"
                     /vntifkey="21"
                     /locus_tag="RSV  promoter"
                     /ApEinfo_fwdcolor="pink"
                     /ApEinfo_revcolor="pink"
                     /ApEinfo_graphicformat="arrow_data {{0 1 2 0 0 -1} {} 0}
                     width 5 offset 0"
     promoter        1927..2190
                     /vntifkey="29"
                     /locus_tag="hU6"
                     /ApEinfo_fwdcolor="#346ee0"
                     /ApEinfo_revcolor="#346ee0"
                     /ApEinfo_graphicformat="arrow_data {{0 1 2 0 0 -1} {} 0}
                     width 5 offset 0"
     LTR             230..410
                     /application_notes="Allows transcription of viral RNA and
                     its packaging into virus."
                     /description="[Truncated HIV-1 5' long terminal repeat]"
                     /full_name="HIV-1 truncated 5' LTR"
                     /note="color: #512bbd; direction: RIGHT"
                     /official_designation="Delta5' LTR"
                     /uuid="44156a3083604321be44066133e8ecb9"
                     /locus_tag="5' LTR"
                     /ApEinfo_fwdcolor="#86f71d"
                     /ApEinfo_revcolor="#d7336f"
                     /ApEinfo_graphicformat="arrow_data {{0 1 2 0 0 -1} {} 0}
                     width 5 offset 0"
     Miscellaneous   521..565
                     /application_notes="Allows packaging of viral RNA into
                     virus."
                     /description="['HIV-1 packaging signal ']"
                     /full_name="HIV-1 psi packaging signal"
                     /note="color: #d84e4e; direction: RIGHT"
                     /official_designation="Psi"
                     /uuid="284899413ff943a99f8210a11ad8c54e"
                     /locus_tag="ÃÂÃÂÃÂÃÂ¨"
                     /ApEinfo_fwdcolor="pink"
                     /ApEinfo_revcolor="pink"
                     /ApEinfo_graphicformat="arrow_data {{0 1 2 0 0 -1} {} 0}
                     width 5 offset 0"
     misc_binding    2213..2216
                     /locus_tag="4nt mismatch (as in Uzonyi et al 2021)"
                     /ApEinfo_fwdcolor="#ff4809"
                     /ApEinfo_revcolor="#ff4809"
                     /ApEinfo_graphicformat="arrow_data {{0 1 2 0 0 -1} {} 0}
                     width 5 offset 0"
     terminator      2267..2272
                     /locus_tag="PolIII terminator"
                     /ApEinfo_fwdcolor="#9d1b1c"
                     /ApEinfo_revcolor="#9d1b1c"
                     /ApEinfo_graphicformat="arrow_data {{0 1 2 0 0 -1} {} 0}
                     width 5 offset 0"
     Miscellaneous   1075..1308
                     /__SeqFeature__="True"
                     /__level__="0"
                     /application_notes="Rev protein binding site that allows
                     Rev-dependent nuclear export of viral RNA during viral
                     packaging."
                     /description="['HIV-1 Rev response element']"
                     /full_name="HIV-1 Rev response element"
                     /name="RRE"
                     /note="color: #1f36a9; direction: RIGHT"
                     /official_designation="RRE"
                     /uuid="df07099e49674cf3aba3661b54e28bd7"
                     /vntifkey="21"
                     /locus_tag="RRE"
                     /ApEinfo_fwdcolor="pink"
                     /ApEinfo_revcolor="pink"
                     /ApEinfo_graphicformat="arrow_data {{0 1 2 0 0 -1} {} 0}
                     width 5 offset 0"
     ORF             3267..3980
                     /application_notes="Low fluorescence and low
                     photostability."
                     /description="Blue variant of EGFP generated by
                     mutagenesis"
                     /full_name="Enhanced blue fluorescent protein(ns)"
                     /note="color: #6ddaae; direction: RIGHT"
                     /official_designation="EBFP(ns)"
                     /uuid="c22d3886879b4745b895a6f25c150792"
                     /locus_tag="EBFP(ns)"
                     /ApEinfo_fwdcolor="#0f7ffe"
                     /ApEinfo_revcolor="pink"
                     /ApEinfo_graphicformat="arrow_data {{0 1 2 0 0 -1} {} 0}
                     width 5 offset 0"
     Miscellaneous   1803..1920
                     /__SeqFeature__="True"
                     /__level__="0"
                     /application_notes="Facilitates the nuclear import of
                     HIV-1 cDNA through a central DNA flap."
                     /description="['Central polypurine tract']"
                     /full_name="Central polypurine tract"
                     /name="cPPT"
                     /note="color: #db6901; direction: RIGHT"
                     /official_designation="cPPT"
                     /uuid="f7cb29985f9f41a7a5924d3462266f42"
                     /vntifkey="21"
                     /locus_tag="cPPT"
                     /ApEinfo_fwdcolor="pink"
                     /ApEinfo_revcolor="pink"
                     /ApEinfo_graphicformat="arrow_data {{0 1 2 0 0 -1} {} 0}
                     width 5 offset 0"
     Linker          3981..4046
                     /application_notes="Causes co-translational cleavage of
                     the encoded polypeptide. Multiple proteins can be made
                     from a polycistronic transcript containing multiple ORFs
                     separated by 2A. P2A and T2A have higher cleavage
                     efficiency compared to other 2As."
                     /description="Self-cleaving 2A peptide from Porcine
                     teschovirus-1"
                     /full_name="Porcine teschovirus-1 2A peptide"
                     /note="color: #1f36a9; direction: RIGHT"
                     /official_designation="P2A"
                     /uuid="26656daa74294a0f8b331614d72d5f5a"
                     /locus_tag="P2A"
                     /ApEinfo_fwdcolor="#20fefe"
                     /ApEinfo_revcolor="pink"
                     /ApEinfo_graphicformat="arrow_data {{0 1 2 0 0 -1} {} 0}
                     width 5 offset 0"
     promoter        2300..2804
                     /__SeqFeature__="True"
                     /__level__="0"
                     /application_notes="Medium-strength promoter."
                     /description="['Human phosphoglycerate kinase 1
                     promoter']"
                     /name="hPGK promoter"
                     /note="color: #5566f5; direction: RIGHT"
                     /official_designation="hPGK promoter"
                     /uuid="a105ee8f48814e769c9a41c38cfff41e"
                     /vntifkey="21"
                     /locus_tag="hPGK promoter"
                     /ApEinfo_fwdcolor="#346ee0"
                     /ApEinfo_revcolor="#346ee0"
                     /ApEinfo_graphicformat="arrow_data {{0 1 2 0 0 -1} {} 0}
                     width 5 offset 0"
     CDS             4047..4445
                     /__SeqFeature__="True"
                     /__level__="0"
                     /application_notes="Allows cells to be resistant to
                     blasticidin."
                     /description="['Blasticidin resistance gene']"
                     /full_name="Blasticidin resistance gene"
                     /name="Bsd"
                     /note="color: #68e66d; direction: RIGHT"
                     /official_designation="Bsd"
                     /uuid="70e7f83c244f451dae86de92afa03717"
                     /vntifkey="21"
                     /locus_tag="Bsd"
                     /ApEinfo_fwdcolor="#e9d024"
                     /ApEinfo_revcolor="#e9d024"
                     /ApEinfo_graphicformat="arrow_data {{0 1 2 0 0 -1} {} 0}
                     width 5 offset 0"
     misc_signal     4477..5074
                     /__SeqFeature__="True"
                     /__level__="0"
                     /application_notes="Enhances virus stability in packaging
                     cells, leading to higher titer of packaged virus; enhances
                     higher expression of transgenes."
                     /description="['Woodchuck hepatitis virus
                     posttranscriptional regulatory element']"
                     /full_name="Woodchuck hepatitis virus posttranscriptional
                     regulatory element"
                     /name="WPRE"
                     /note="color: #ef6cdf; direction: RIGHT"
                     /official_designation="WPRE"
                     /uuid="0aedff8308474b369c622433538d7836"
                     /vntifkey="21"
                     /locus_tag="WPRE"
                     /ApEinfo_fwdcolor="#008040"
                     /ApEinfo_revcolor="#008040"
                     /ApEinfo_graphicformat="arrow_data {{0 1 2 0 0 -1} {} 0}
                     width 5 offset 0"
     LTR             5140..5374
                     /application_notes="Allows packaging of viral RNA into
                     virus; self-inactivates the 5' LTR by a copying mechanism
                     during viral genome integration; contains polyadenylation
                     signal for transcription termination."
                     /description="[Truncated HIV-1 3' long terminal repeat ]"
                     /full_name="HIV-1 truncated 3' LTR"
                     /note="color: #c54b7c; direction: RIGHT"
                     /official_designation="DeltaU3/3' LTR"
                     /uuid="c958755414c34372b909e20e81ebcef6"
                     /locus_tag="U3/3' LTR"
                     /ApEinfo_fwdcolor="#86f71d"
                     /ApEinfo_revcolor="#d7336f"
                     /ApEinfo_graphicformat="arrow_data {{0 1 2 0 0 -1} {} 0}
                     width 5 offset 0"
     polyA_signal    5447..5581
                     /__SeqFeature__="True"
                     /__level__="0"
                     /application_notes="Allows transcription termination and
                     polyadenylation of mRNA transcribed by Pol II RNA
                     polymerase."
                     /description="['Simian virus 40 early polyadenylation
                     signal']"
                     /full_name="SV40 early polyadenation signal"
                     /name="SV40 early pA"
                     /note="color: #d05c0a; direction: RIGHT"
                     /official_designation="SV40 early pA"
                     /uuid="2fcb13a62e704d8095b188d9625a6a94"
                     /vntifkey="21"
                     /locus_tag="SV40 early pA"
                     /ApEinfo_fwdcolor="#ff3eee"
                     /ApEinfo_revcolor="#ff3eee"
                     /ApEinfo_graphicformat="arrow_data {{0 1 2 0 0 -1} {} 0}
                     width 5 offset 0"
     ORF             6535..7395
                     /__SeqFeature__="True"
                     /__level__="0"
                     /application_notes="Allows E. coli to be resistant to
                     ampicillin."
                     /description="['AmpiciIIin resistance gene']"
                     /full_name="Ampicillin resistance gene"
                     /name="AmpiciIIin"
                     /note="color: #6ddaae; direction: RIGHT"
                     /official_designation="Ampicillin"
                     /uuid="6a7994dc58024d80b36e4a85ad2e1d99"
                     /vntifkey="21"
                     /locus_tag="AmpiciIIin"
                     /ApEinfo_fwdcolor="pink"
                     /ApEinfo_revcolor="pink"
                     /ApEinfo_graphicformat="arrow_data {{0 1 2 0 0 -1} {} 0}
                     width 5 offset 0"
     Rep_origin      7566..8154
                     /__SeqFeature__="True"
                     /__level__="0"
                     /application_notes="Facilitates plasmid replication in E.
                     coli; regulates high-copy plasmid number (500-700)."
                     /description="['pUC origin of replication']"
                     /full_name="pUC origin of replication"
                     /name="pUC ori"
                     /note="color: #fd3434; direction: RIGHT"
                     /official_designation="pUC ori"
                     /uuid="64a8cb3def9f49deb6752795c2d893a8"
                     /vntifkey="21"
                     /locus_tag="pUC ori"
                     /ApEinfo_fwdcolor="pink"
                     /ApEinfo_revcolor="pink"
                     /ApEinfo_graphicformat="arrow_data {{0 1 2 0 0 -1} {} 0}
                     width 5 offset 0"
     primer_bind     8578..8597
                     /locus_tag="T3"
                     /ApEinfo_fwdcolor="cyan"
                     /ApEinfo_revcolor="green"
                     /ApEinfo_graphicformat="arrow_data {{0 1 2 0 0 -1} {} 0}
                     width 5 offset 0"
     primer_bind     complement(5791..5808)
                     /locus_tag="M13-fwd"
                     /ApEinfo_fwdcolor="cyan"
                     /ApEinfo_revcolor="green"
                     /ApEinfo_graphicformat="arrow_data {{0 1 2 0 0 -1} {} 0}
                     width 5 offset 0"
     primer_bind     8540..8560
                     /locus_tag="M13-rev"
                     /ApEinfo_fwdcolor="cyan"
                     /ApEinfo_revcolor="green"
                     /ApEinfo_graphicformat="arrow_data {{0 1 2 0 0 -1} {} 0}
                     width 5 offset 0"
     primer_bind     complement(5762..5782)
                     /locus_tag="T7"
                     /ApEinfo_fwdcolor="cyan"
                     /ApEinfo_revcolor="green"
                     /ApEinfo_graphicformat="arrow_data {{0 1 2 0 0 -1} {} 0}
                     width 5 offset 0"
     rep_origin      5966..6272
                     /locus_tag="F1 ori"
                     /ApEinfo_fwdcolor="gray50"
                     /ApEinfo_revcolor="gray50"
                     /ApEinfo_graphicformat="arrow_data {{0 1 2 0 0 -1} {} 0}
                     width 5 offset 0"
     CDS             5879..5947
                     /locus_tag="LacZ alpha"
                     /ApEinfo_fwdcolor="#6495ed"
                     /ApEinfo_revcolor="#6495ed"
                     /ApEinfo_graphicformat="arrow_data {{0 1 2 0 0 -1} {} 0}
                     width 5 offset 0"
     misc_binding    8512..8534
                     /locus_tag="LacO"
                     /ApEinfo_fwdcolor="#6495ed"
                     /ApEinfo_revcolor="#6495ed"
                     /ApEinfo_graphicformat="arrow_data {{0 1 2 0 0 -1} {} 0}
                     width 5 offset 0"
     CDS             6733..7392
                     /locus_tag="AmpR"
                     /ApEinfo_fwdcolor="yellow"
                     /ApEinfo_revcolor="yellow"
                     /ApEinfo_graphicformat="arrow_data {{0 1 2 0 0 -1} {} 0}
                     width 5 offset 0"
     CDS             2835..3191
                     /standard_name="MLANA_cds"
                     /locus_tag="MLANA_cds"
                     /ApEinfo_fwdcolor="#e9d024"
                     /ApEinfo_revcolor="#e9d024"
                     /ApEinfo_graphicformat="arrow_data {{0 1 2 0 0 -1} {} 0}
                     width 5 offset 0"
     sig_peptide     3213..3266
                     /standard_name="T2A"
                     /note="Geneious type: LINKER"
                     /locus_tag="T2A"
                     /ApEinfo_fwdcolor="cyan"
                     /ApEinfo_revcolor="green"
                     /ApEinfo_graphicformat="arrow_data {{0 1 2 0 0 -1} {} 0}
                     width 5 offset 0"
     misc_feature    2910..2939
                     /standard_name="MART-1_26-35 G6S"
                     /locus_tag="MART-1_26-35"
                     /ApEinfo_fwdcolor="#ff7d78"
                     /ApEinfo_revcolor="#7eff74"
                     /ApEinfo_graphicformat="arrow_data {{0 1 2 0 0 -1} {} 0}
                     width 5 offset 0"
     misc_feature    3192..3212
                     /locus_tag="linker"
                     /ApEinfo_fwdcolor="#a8eafe"
                     /ApEinfo_revcolor="green"
                     /ApEinfo_graphicformat="arrow_data {{0 1 2 0 0 -1} {} 0}
                     width 5 offset 0"
     misc_feature    2925..2927
                     /standard_name="G6S"
                     /locus_tag="G31S"
                     /ApEinfo_fwdcolor="#941100"
                     /ApEinfo_revcolor="#7eff74"
                     /ApEinfo_graphicformat="arrow_data {{0 1 2 0 0 -1} {} 0}
                     width 5 offset 0"
     variation       2925..2925
                     /locus_tag="target A"
                     /ApEinfo_fwdcolor="#ff9e39"
                     /ApEinfo_revcolor="#ff9e39"
                     /ApEinfo_graphicformat="arrow_data {{0 1 2 0 0 -1} {} 0}
                     width 5 offset 0"
ORIGIN
        1 aatgtagtct tatgcaatac tcttgtagtc ttgcaacatg gtaacgatga gttagcaaca
       61 tgccttacaa ggagagaaaa agcaccgtgc atgccgattg gtggaagtaa ggtggtacga
      121 tcgtgcctta ttaggaaggc aacagacggg tctgacatgg attggacgaa ccactgaatt
      181 gccgcattgc agagatattg tatttaagtg cctagctcga tacataaacg ggtctctctg
      241 gttagaccag atctgagcct gggagctctc tggctaacta gggaacccac tgcttaagcc
      301 tcaataaagc ttgccttgag tgcttcaagt agtgtgtgcc cgtctgttgt gtgactctgg
      361 taactagaga tccctcagac ccttttagtc agtgtggaaa atctctagca gtggcgcccg
      421 aacagggact tgaaagcgaa agggaaacca gaggagctct ctcgacgcag gactcggctt
      481 gctgaagcgc gcacggcaag aggcgagggg cggcgactgg tgagtacgcc aaaaattttg
      541 actagcggag gctagaagga gagagatggg tgcgagagcg tcagtattaa gcgggggaga
      601 attagatcgc gatgggaaaa aattcggtta aggccagggg gaaagaaaaa atataaatta
      661 aaacatatag tatgggcaag cagggagcta gaacgattcg cagttaatcc tggcctgtta
      721 gaaacatcag aaggctgtag acaaatactg ggacagctac aaccatccct tcagacagga
      781 tcagaagaac ttagatcatt atataataca gtagcaaccc tctattgtgt gcatcaaagg
      841 atagagataa aagacaccaa ggaagcttta gacaagatag aggaagagca aaacaaaagt
      901 aagaccaccg cacagcaagc ggccgctgat cttcagacct ggaggaggag atatgaggga
      961 caattggaga agtgaattat ataaatataa agtagtaaaa attgaaccat taggagtagc
     1021 acccaccaag gcaaagagaa gagtggtgca gagagaaaaa agagcagtgg gaataggagc
     1081 tttgttcctt gggttcttgg gagcagcagg aagcactatg ggcgcagcgt caatgacgct
     1141 gacggtacag gccagacaat tattgtctgg tatagtgcag cagcagaaca atttgctgag
     1201 ggctattgag gcgcaacagc atctgttgca actcacagtc tggggcatca agcagctcca
     1261 ggcaagaatc ctggctgtgg aaagatacct aaaggatcaa cagctcctgg ggatttgggg
     1321 ttgctctgga aaactcattt gcaccactgc tgtgccttgg aatgctagtt ggagtaataa
     1381 atctctggaa cagatttgga atcacacgac ctggatggag tgggacagag aaattaacaa
     1441 ttacacaagc ttaatacact ccttaattga agaatcgcaa aaccagcaag aaaagaatga
     1501 acaagaatta ttggaattag ataaatgggc aagtttgtgg aattggttta acataacaaa
     1561 ttggctgtgg tatataaaat tattcataat gatagtagga ggcttggtag gtttaagaat
     1621 agtttttgct gtactttcta tagtgaatag agttaggcag ggatattcac cattatcgtt
     1681 tcagacccac ctcccaaccc cgaggggacc cgacaggccc gaaggaatag aagaagaagg
     1741 tggagagaga gacagagaca gatccattcg attagtgaac ggatctcgac ggtatcgcta
     1801 gcttttaaaa gaaaaggggg gattgggggg tacagtgcag gggaaagaat agtagacata
     1861 atagcaacag acatacaaac taaagaatta caaaaacaaa ttacaaaaat tcaaaatttt
     1921 actagtaagg tcgggcagga agagggccta tttcccatga ttccttcata tttgcatata
     1981 cgatacaagg ctgttagaga gataattaga attaatttga ctgtaaacac aaagatatta
     2041 gtacaaaata cgtgacgtag aaagtaataa tttcttgggt agtttgcagt tttaaaatta
     2101 tgttttaaaa tggactatca tatgcttacc gtaacttgaa agtatttcga tttcttggct
     2161 ttatatatct tgtggaaagg acgaaacacc GGtcggggta cctgctgaag caaacgacgc
     2221 cgtatgtcag ggtagtgaca agtgttggcC atggaacagg tagttttttt ttgaattcca
     2281 actttgtata gaaaagttgg ggttgcgcct tttccaaggc agccctgggt ttgcgcaggg
     2341 acgcggctgc tctgggcgtg gttccgggaa acgcagcggc gccgaccctg ggtctcgcac
     2401 attcttcacg tccgttcgca gcgtcacccg gatcttcgcc gctacccttg tgggcccccc
     2461 ggcgacgctt cctgctccgc ccctaagtcg ggaaggttcc ttgcggttcg cggcgtgccg
     2521 gacgtgacaa acggaagccg cacgtctcac tagtaccctc gcagacggac agcgccaggg
     2581 agcaatggca gcgcgccgac cgcgatgggc tgtggccaat agcggctgct cagcagggcg
     2641 cgccgagagc agcggccggg aaggggcggt gcgggaggcg gggtgtgggg cggtagtgtg
     2701 ggccctgttc ctgcccgcgc ggtgttccgc attctgcaag cctccggagc gcacgtcggc
     2761 agtcggctcc ctcgttgacc gaatcaccga cctctctccc caggcaagtt tgtacaaaaa
     2821 agcaggctgc caccatgccc agggaggacg cccacttcat ctacggctac cccaagaagg
     2881 gccacggcca cagctacacc actgcagagg agcttgctgg gatcagcatc ctgacagtga
     2941 tcctaggcgt gctgctgctg atcggctgct ggtactgccg gaggaggaac ggctacaggg
     3001 ccctgatgga caagagcctg cacgtgggca cccagtgcgc cctgaccagg aggtgccccc
     3061 aggagggctt cgaccacagg gacagcaagg tgagcctcca ggagaagaac tgcgagcccg
     3121 tggtgcccaa cgcccccccc gcctacgaga agctgagcgc cgagcagagc cctccaccat
     3181 acagccccgg tggatccgga ggtgcgagcg gcgaggggag gggcagcctt cttacttgcg
     3241 gagatgtgga agagaatcca ggacccgtga gcaagggcga ggagctgttc accggggtgg
     3301 tgcccatcct ggtcgagctg gacggcgacg taaacggcca caagttcagc gtgtccggcg
     3361 agggcgaggg cgatgccacc tacggcaagc tgaccctgaa gttcatctgc accaccggca
     3421 agctgcccgt gccctggccc accctcgtga ccaccctgac ccacggcgtg cagtgcttca
     3481 gccgctaccc cgaccacatg aagcagcacg acttcttcaa gtccgccatg cccgaaggct
     3541 acgtccagga gcgcaccatc ttcttcaagg acgacggcaa ctacaagacc cgcgccgagg
     3601 tgaagttcga gggcgacacc ctggtgaacc gcatcgagct gaagggcatc gacttcaagg
     3661 aggacggcaa catcctgggg cacaagctgg agtacaactt caacagccac aacgtctata
     3721 tcatggccga caagcagaag aacggcatca aggtgaactt caagatccgc cacaacatcg
     3781 aggacggcag cgtgcagctc gccgaccact accagcagaa cacccccatc ggcgacggcc
     3841 ccgtgctgct gcccgacaac cactacctga gcacccagtc cgccctgagc aaagacccca
     3901 acgagaagcg cgatcacatg gtcctgctgg agttcgtgac cgccgccggg atcactctcg
     3961 gcatggacga gctgtacaag ggaagcggag ccacgaactt ctctctgtta aagcaagcag
     4021 gagatgttga agaaaacccc gggcctatgg ccaagccttt gtctcaagaa gaatccaccc
     4081 tcattgaaag agcaacggct acaatcaaca gcatccccat ctctgaagac tacagcgtcg
     4141 ccagcgcagc tctctctagc gacggccgca tcttcactgg tgtcaatgta tatcatttta
     4201 ctgggggacc ttgtgcagaa ctcgtggtgc tgggcactgc tgctgctgcg gcagctggca
     4261 acctgacttg tatcgtcgcg atcggaaatg agaacagggg catcttgagc ccctgcggac
     4321 ggtgccgaca ggtgcttctc gatctgcatc ctgggatcaa agccatagtg aaggacagtg
     4381 atggacagcc gacggcagtt gggattcgtg aattgctgcc ctctggttat gtgtgggagg
     4441 gctaaaccca gctttcttgt acaaagtggt ggtacccgat aatcaacctc tggattacaa
     4501 aatttgtgaa agattgactg gtattcttaa ctatgttgct ccttttacgc tatgtggata
     4561 cgctgcttta atgcctttgt atcatgctat tgcttcccgt atggctttca ttttctcctc
     4621 cttgtataaa tcctggttgc tgtctcttta tgaggagttg tggcccgttg tcaggcaacg
     4681 tggcgtggtg tgcactgtgt ttgctgacgc aacccccact ggttggggca ttgccaccac
     4741 ctgtcagctc ctttccggga ctttcgcttt ccccctccct attgccacgg cggaactcat
     4801 cgccgcctgc cttgcccgct gctggacagg ggctcggctg ttgggcactg acaattccgt
     4861 ggtgttgtcg gggaagctga cgtcctttcc atggctgctc gcctgtgttg ccacctggat
     4921 tctgcgcggg acgtccttct gctacgtccc ttcggccctc aatccagcgg accttccttc
     4981 ccgcggcctg ctgccggctc tgcggcctct tccgcgtctt cgccttcgcc ctcagacgag
     5041 tcggatctcc ctttgggccg cctccccgca tcggctttaa gaccaatgac ttacaaggca
     5101 gctgtagatc ttagccactt tttaaaagaa aaggggggac tggaagggct aattcactcc
     5161 caacgaagac aagatctgct ttttgcttgt actgggtctc tctggttaga ccagatctga
     5221 gcctgggagc tctctggcta actagggaac ccactgctta agcctcaata aagcttgcct
     5281 tgagtgcttc aagtagtgtg tgcccgtctg ttgtgtgact ctggtaacta gagatccctc
     5341 agaccctttt agtcagtgtg gaaaatctct agcagtagta gttcatgtca tcttattatt
     5401 cagtatttat aacttgcaaa gaaatgaata tcagagagtg agaggaactt gtttattgca
     5461 gcttataatg gttacaaata aagcaatagc atcacaaatt tcacaaataa agcatttttt
     5521 tcactgcatt ctagttgtgg tttgtccaaa ctcatcaatg tatcttatca tgtctggctc
     5581 tagctatccc gcccctaact ccgcccatcc cgcccctaac tccgcccagt tccgcccatt
     5641 ctccgcccca tggctgacta atttttttta tttatgcaga ggccgaggcc gcctcggcct
     5701 ctgagctatt ccagaagtag tgaggaggct tttttggagg cctagggacg tacccaattc
     5761 gccctatagt gagtcgtatt acgcgcgctc actggccgtc gttttacaac gtcgtgactg
     5821 ggaaaaccct ggcgttaccc aacttaatcg ccttgcagca catccccctt tcgccagctg
     5881 gcgtaatagc gaagaggccc gcaccgatcg cccttcccaa cagttgcgca gcctgaatgg
     5941 cgaatgggac gcgccctgta gcggcgcatt aagcgcggcg ggtgtggtgg ttacgcgcag
     6001 cgtgaccgct acacttgcca gcgccctagc gcccgctcct ttcgctttct tcccttcctt
     6061 tctcgccacg ttcgccggct ttccccgtca agctctaaat cgggggctcc ctttagggtt
     6121 ccgatttagt gctttacggc acctcgaccc caaaaaactt gattagggtg atggttcacg
     6181 tagtgggcca tcgccctgat agacggtttt tcgccctttg acgttggagt ccacgttctt
     6241 taatagtgga ctcttgttcc aaactggaac aacactcaac cctatctcgg tctattcttt
     6301 tgatttataa gggattttgc cgatttcggc ctattggtta aaaaatgagc tgatttaaca
     6361 aaaatttaac gcgaatttta acaaaatatt aacgcttaca atttaggtgg cacttttcgg
     6421 ggaaatgtgc gcggaacccc tatttgttta tttttctaaa tacattcaaa tatgtatccg
     6481 ctcatgagac aataaccctg ataaatgctt caataatatt gaaaaaggaa gagtatgagt
     6541 attcaacatt tccgtgtcgc ccttattccc ttttttgcgg cattttgcct tcctgttttt
     6601 gctcacccag aaacgctggt gaaagtaaaa gatgctgaag atcagttggg tgcacgagtg
     6661 ggttacatcg aactggatct caacagcggt aagatccttg agagttttcg ccccgaagaa
     6721 cgttttccaa tgatgagcac ttttaaagtt ctgctatgtg gcgcggtatt atcccgtatt
     6781 gacgccgggc aagagcaact cggtcgccgc atacactatt ctcagaatga cttggttgag
     6841 tactcaccag tcacagaaaa gcatcttacg gatggcatga cagtaagaga attatgcagt
     6901 gctgccataa ccatgagtga taacactgcg gccaacttac ttctgacaac gatcggagga
     6961 ccgaaggagc taaccgcttt tttgcacaac atgggggatc atgtaactcg ccttgatcgt
     7021 tgggaaccgg agctgaatga agccatacca aacgacgagc gtgacaccac gatgcctgta
     7081 gcaatggcaa caacgttgcg caaactatta actggcgaac tacttactct agcttcccgg
     7141 caacaattaa tagactggat ggaggcggat aaagttgcag gaccacttct gcgctcggcc
     7201 cttccggctg gctggtttat tgctgataaa tctggagccg gtgagcgtgg gtctcgcggt
     7261 atcattgcag cactggggcc agatggtaag ccctcccgta tcgtagttat ctacacgacg
     7321 gggagtcagg caactatgga tgaacgaaat agacagatcg ctgagatagg tgcctcactg
     7381 attaagcatt ggtaactgtc agaccaagtt tactcatata tactttagat tgatttaaaa
     7441 cttcattttt aatttaaaag gatctaggtg aagatccttt ttgataatct catgaccaaa
     7501 atcccttaac gtgagttttc gttccactga gcgtcagacc ccgtagaaaa gatcaaagga
     7561 tcttcttgag atcctttttt tctgcgcgta atctgctgct tgcaaacaaa aaaaccaccg
     7621 ctaccagcgg tggtttgttt gccggatcaa gagctaccaa ctctttttcc gaaggtaact
     7681 ggcttcagca gagcgcagat accaaatact gttcttctag tgtagccgta gttaggccac
     7741 cacttcaaga actctgtagc accgcctaca tacctcgctc tgctaatcct gttaccagtg
     7801 gctgctgcca gtggcgataa gtcgtgtctt accgggttgg actcaagacg atagttaccg
     7861 gataaggcgc agcggtcggg ctgaacgggg ggttcgtgca cacagcccag cttggagcga
     7921 acgacctaca ccgaactgag atacctacag cgtgagctat gagaaagcgc cacgcttccc
     7981 gaagagagaa aggcggacag gtatccggta agcggcaggg tcggaacagg agagcgcacg
     8041 agggagcttc cagggggaaa cgcctggtat ctttatagtc ctgtcgggtt tcgccacctc
     8101 tgacttgagc gtcgattttt gtgatgctcg tcaggggggc ggagcctatg gaaaaacgcc
     8161 agcaacgcgg cctttttacg gttcctggcc ttttgctggc cttttgctca catgttcttt
     8221 cctgcgttat cccctgattc tgtggataac cgtattaccg cctttgagtg agctgatacc
     8281 gctcgccgca gccgaacgac cgagcgcagc gagtcagtga gcgaggaagc ggaagagcgc
     8341 ccaatacgca aaccgcctct ccccgcgcgt tggccgattc attaatgcag ctggcacgac
     8401 aggtttcccg actggaaagc gggcagtgag cgcaacgcaa ttaatgtgag ttagctcact
     8461 cattaggcac cccaggcttt acactttatg cttccggctc gtatgttgtg tggaattgtg
     8521 agcggataac aatttcacac aggaaacagc tatgaccatg attacgccaa gcgcgcaatt
     8581 aaccctcact aaagggaaca aaagctggag ctgcaagctt
//
```

**Plasmid hU6 T-SPEAR (coMART-1^G31S^)_hPGK coMART1^G31S^-T2A-BFP-P2A-Bsd sequence (GenBank format, provided as text)**

```
LOCUS       RP500_pLV_hU6_20        8620 bp    DNA        circular     21-APR-2023
DEFINITION
ACCESSION   VB210525-1302qyy
VERSION     VB210525-1302qyy
KEYWORDS    .
SOURCE
  ORGANISM  .
COMMENT
COMMENT     ApEinfo:methylated:1
FEATURES             Location/Qualifiers
     Promoter        1..229
                     /__SeqFeature__="True"
                     /__level__="0"
                     /application_notes="Strong promoter; drives transcription
                     of viral RNA in packaging cells."
                     /description="['Rous sarcoma virus enhancer/promoter']"
                     /full_name="Rous sarcoma virus (RSV) enhancer/promoter"
                     /name="RSV  promoter"
                     /note="color: #946a00; direction: RIGHT"
                     /official_designation="RSV"
                     /uuid="3f2ddca7f0cb4015b6abcec8a25c834e"
                     /vntifkey="21"
                     /locus_tag="RSV  promoter"
                     /label="RSV  promoter"
                     /ApEinfo_label="RSV  promoter"
                     /ApEinfo_fwdcolor="pink"
                     /ApEinfo_revcolor="pink"
                     /ApEinfo_graphicformat="arrow_data {{0 1 2 0 0 -1} {} 0}
                     width 5 offset 0"
     promoter        1927..2190
                     /vntifkey="29"
                     /locus_tag="hU6"
                     /label="hU6"
                     /ApEinfo_label="hU6"
                     /ApEinfo_fwdcolor="#346ee0"
                     /ApEinfo_revcolor="#346ee0"
                     /ApEinfo_graphicformat="arrow_data {{0 1 2 0 0 -1} {} 0}
                     width 5 offset 0"
     LTR             230..410
                     /application_notes="Allows transcription of viral RNA and
                     its packaging into virus."
                     /description="[Truncated HIV-1 5' long terminal repeat]"
                     /full_name="HIV-1 truncated 5' LTR"
                     /note="color: #512bbd; direction: RIGHT"
                     /official_designation="Delta5' LTR"
                     /uuid="44156a3083604321be44066133e8ecb9"
                     /locus_tag="5' LTR"
                     /label="5' LTR"
                     /ApEinfo_label="5' LTR"
                     /ApEinfo_fwdcolor="#86f71d"
                     /ApEinfo_revcolor="#d7336f"
                     /ApEinfo_graphicformat="arrow_data {{0 1 2 0 0 -1} {} 0}
                     width 5 offset 0"
     Miscellaneous   521..565
                     /application_notes="Allows packaging of viral RNA into
                     virus."
                     /description="['HIV-1 packaging signal ']"
                     /full_name="HIV-1 psi packaging signal"
                     /note="color: #d84e4e; direction: RIGHT"
                     /official_designation="Psi"
                     /uuid="284899413ff943a99f8210a11ad8c54e"
                     /locus_tag="ÃÂÃÂÃÂÃÂ¨"
                     /label="ÃÂÃÂÃÂÃÂ¨"
                     /ApEinfo_label="ÃÂÃÂÃÂÃÂ¨"
                     /ApEinfo_fwdcolor="pink"
                     /ApEinfo_revcolor="pink"
                     /ApEinfo_graphicformat="arrow_data {{0 1 2 0 0 -1} {} 0}
                     width 5 offset 0"
     terminator      2267..2272
                     /locus_tag="PolIII terminator"
                     /label="PolIII terminator"
                     /ApEinfo_label="PolIII terminator"
                     /ApEinfo_fwdcolor="#9d1b1c"
                     /ApEinfo_revcolor="#9d1b1c"
                     /ApEinfo_graphicformat="arrow_data {{0 1 2 0 0 -1} {} 0}
                     width 5 offset 0"
     misc_binding    2213..2216
                     /bound_moiety=""
                     /locus_tag="4nt mismatch (as in Uzonyi et al 2021)"
                     /label="4nt mismatch (as in Uzonyi et al 2021)"
                     /ApEinfo_label="4nt mismatch (as in Uzonyi et al 2021)"
                     /ApEinfo_fwdcolor="#ff4809"
                     /ApEinfo_revcolor="#ff4809"
                     /ApEinfo_graphicformat="arrow_data {{0 0.5 0 1 2 0 0 -1 0
                     -0.5} {} 0} width 5 offset 0"
     Miscellaneous   1075..1308
                     /__SeqFeature__="True"
                     /__level__="0"
                     /application_notes="Rev protein binding site that allows
                     Rev-dependent nuclear export of viral RNA during viral
                     packaging."
                     /description="['HIV-1 Rev response element']"
                     /full_name="HIV-1 Rev response element"
                     /name="RRE"
                     /note="color: #1f36a9; direction: RIGHT"
                     /official_designation="RRE"
                     /uuid="df07099e49674cf3aba3661b54e28bd7"
                     /vntifkey="21"
                     /locus_tag="RRE"
                     /label="RRE"
                     /ApEinfo_label="RRE"
                     /ApEinfo_fwdcolor="pink"
                     /ApEinfo_revcolor="pink"
                     /ApEinfo_graphicformat="arrow_data {{0 1 2 0 0 -1} {} 0}
                     width 5 offset 0"
     ORF             3267..3980
                     /application_notes="Low fluorescence and low
                     photostability."
                     /description="Blue variant of EGFP generated by
                     mutagenesis"
                     /full_name="Enhanced blue fluorescent protein(ns)"
                     /note="color: #6ddaae; direction: RIGHT"
                     /official_designation="EBFP(ns)"
                     /uuid="c22d3886879b4745b895a6f25c150792"
                     /locus_tag="EBFP(ns)"
                     /label="EBFP(ns)"
                     /ApEinfo_label="EBFP(ns)"
                     /ApEinfo_fwdcolor="#0f7ffe"
                     /ApEinfo_revcolor="pink"
                     /ApEinfo_graphicformat="arrow_data {{0 1 2 0 0 -1} {} 0}
                     width 5 offset 0"
     Miscellaneous   1803..1920
                     /__SeqFeature__="True"
                     /__level__="0"
                     /application_notes="Facilitates the nuclear import of
                     HIV-1 cDNA through a central DNA flap."
                     /description="['Central polypurine tract']"
                     /full_name="Central polypurine tract"
                     /name="cPPT"
                     /note="color: #db6901; direction: RIGHT"
                     /official_designation="cPPT"
                     /uuid="f7cb29985f9f41a7a5924d3462266f42"
                     /vntifkey="21"
                     /locus_tag="cPPT"
                     /label="cPPT"
                     /ApEinfo_label="cPPT"
                     /ApEinfo_fwdcolor="pink"
                     /ApEinfo_revcolor="pink"
                     /ApEinfo_graphicformat="arrow_data {{0 1 2 0 0 -1} {} 0}
                     width 5 offset 0"
     Linker          3981..4046
                     /application_notes="Causes co-translational cleavage of
                     the encoded polypeptide. Multiple proteins can be made
                     from a polycistronic transcript containing multiple ORFs
                     separated by 2A. P2A and T2A have higher cleavage
                     efficiency compared to other 2As."
                     /description="Self-cleaving 2A peptide from Porcine
                     teschovirus-1"
                     /full_name="Porcine teschovirus-1 2A peptide"
                     /note="color: #1f36a9; direction: RIGHT"
                     /official_designation="P2A"
                     /uuid="26656daa74294a0f8b331614d72d5f5a"
                     /locus_tag="P2A"
                     /label="P2A"
                     /ApEinfo_label="P2A"
                     /ApEinfo_fwdcolor="#20fefe"
                     /ApEinfo_revcolor="pink"
                     /ApEinfo_graphicformat="arrow_data {{0 1 2 0 0 -1} {} 0}
                     width 5 offset 0"
     promoter        2300..2804
                     /__SeqFeature__="True"
                     /__level__="0"
                     /application_notes="Medium-strength promoter."
                     /description="['Human phosphoglycerate kinase 1
                     promoter']"
                     /name="hPGK promoter"
                     /note="color: #5566f5; direction: RIGHT"
                     /official_designation="hPGK promoter"
                     /uuid="a105ee8f48814e769c9a41c38cfff41e"
                     /vntifkey="21"
                     /locus_tag="hPGK promoter"
                     /label="hPGK promoter"
                     /ApEinfo_label="hPGK promoter"
                     /ApEinfo_fwdcolor="#346ee0"
                     /ApEinfo_revcolor="#346ee0"
                     /ApEinfo_graphicformat="arrow_data {{0 1 2 0 0 -1} {} 0}
                     width 5 offset 0"
     CDS             4047..4445
                     /__SeqFeature__="True"
                     /__level__="0"
                     /application_notes="Allows cells to be resistant to
                     blasticidin."
                     /description="['Blasticidin resistance gene']"
                     /full_name="Blasticidin resistance gene"
                     /name="Bsd"
                     /note="color: #68e66d; direction: RIGHT"
                     /official_designation="Bsd"
                     /uuid="70e7f83c244f451dae86de92afa03717"
                     /vntifkey="21"
                     /locus_tag="Bsd"
                     /label="Bsd"
                     /ApEinfo_label="Bsd"
                     /ApEinfo_fwdcolor="#e9d024"
                     /ApEinfo_revcolor="#e9d024"
                     /ApEinfo_graphicformat="arrow_data {{0 1 2 0 0 -1} {} 0}
                     width 5 offset 0"
     misc_signal     4477..5074
                     /__SeqFeature__="True"
                     /__level__="0"
                     /application_notes="Enhances virus stability in packaging
                     cells, leading to higher titer of packaged virus; enhances
                     higher expression of transgenes."
                     /description="['Woodchuck hepatitis virus
                     posttranscriptional regulatory element']"
                     /full_name="Woodchuck hepatitis virus posttranscriptional
                     regulatory element"
                     /name="WPRE"
                     /note="color: #ef6cdf; direction: RIGHT"
                     /official_designation="WPRE"
                     /uuid="0aedff8308474b369c622433538d7836"
                     /vntifkey="21"
                     /locus_tag="WPRE"
                     /label="WPRE"
                     /ApEinfo_label="WPRE"
                     /ApEinfo_fwdcolor="#008040"
                     /ApEinfo_revcolor="#008040"
                     /ApEinfo_graphicformat="arrow_data {{0 1 2 0 0 -1} {} 0}
                     width 5 offset 0"
     LTR             5140..5374
                     /application_notes="Allows packaging of viral RNA into
                     virus; self-inactivates the 5' LTR by a copying mechanism
                     during viral genome integration; contains polyadenylation
                     signal for transcription termination."
                     /description="[Truncated HIV-1 3' long terminal repeat ]"
                     /full_name="HIV-1 truncated 3' LTR"
                     /note="color: #c54b7c; direction: RIGHT"
                     /official_designation="DeltaU3/3' LTR"
                     /uuid="c958755414c34372b909e20e81ebcef6"
                     /locus_tag="U3/3' LTR"
                     /label="U3/3' LTR"
                     /ApEinfo_label="U3/3' LTR"
                     /ApEinfo_fwdcolor="#86f71d"
                     /ApEinfo_revcolor="#d7336f"
                     /ApEinfo_graphicformat="arrow_data {{0 1 2 0 0 -1} {} 0}
                     width 5 offset 0"
     polyA_signal    5447..5581
                     /__SeqFeature__="True"
                     /__level__="0"
                     /application_notes="Allows transcription termination and
                     polyadenylation of mRNA transcribed by Pol II RNA
                     polymerase."
                     /description="['Simian virus 40 early polyadenylation
                     signal']"
                     /full_name="SV40 early polyadenation signal"
                     /name="SV40 early pA"
                     /note="color: #d05c0a; direction: RIGHT"
                     /official_designation="SV40 early pA"
                     /uuid="2fcb13a62e704d8095b188d9625a6a94"
                     /vntifkey="21"
                     /locus_tag="SV40 early pA"
                     /label="SV40 early pA"
                     /ApEinfo_label="SV40 early pA"
                     /ApEinfo_fwdcolor="#ff3eee"
                     /ApEinfo_revcolor="#ff3eee"
                     /ApEinfo_graphicformat="arrow_data {{0 1 2 0 0 -1} {} 0}
                     width 5 offset 0"
     ORF             6535..7395
                     /__SeqFeature__="True"
                     /__level__="0"
                     /application_notes="Allows E. coli to be resistant to
                     ampicillin."
                     /description="['AmpiciIIin resistance gene']"
                     /full_name="Ampicillin resistance gene"
                     /name="AmpiciIIin"
                     /note="color: #6ddaae; direction: RIGHT"
                     /official_designation="Ampicillin"
                     /uuid="6a7994dc58024d80b36e4a85ad2e1d99"
                     /vntifkey="21"
                     /locus_tag="AmpiciIIin"
                     /label="AmpiciIIin"
                     /ApEinfo_label="AmpiciIIin"
                     /ApEinfo_fwdcolor="pink"
                     /ApEinfo_revcolor="pink"
                     /ApEinfo_graphicformat="arrow_data {{0 1 2 0 0 -1} {} 0}
                     width 5 offset 0"
     Rep_origin      7566..8154
                     /__SeqFeature__="True"
                     /__level__="0"
                     /application_notes="Facilitates plasmid replication in E.
                     coli; regulates high-copy plasmid number (500-700)."
                     /description="['pUC origin of replication']"
                     /full_name="pUC origin of replication"
                     /name="pUC ori"
                     /note="color: #fd3434; direction: RIGHT"
                     /official_designation="pUC ori"
                     /uuid="64a8cb3def9f49deb6752795c2d893a8"
                     /vntifkey="21"
                     /locus_tag="pUC ori"
                     /label="pUC ori"
                     /ApEinfo_label="pUC ori"
                     /ApEinfo_fwdcolor="pink"
                     /ApEinfo_revcolor="pink"
                     /ApEinfo_graphicformat="arrow_data {{0 1 2 0 0 -1} {} 0}
                     width 5 offset 0"
     primer_bind     8578..8597
                     /locus_tag="T3"
                     /label="T3"
                     /ApEinfo_label="T3"
                     /ApEinfo_fwdcolor="cyan"
                     /ApEinfo_revcolor="green"
                     /ApEinfo_graphicformat="arrow_data {{0 1 2 0 0 -1} {} 0}
                     width 5 offset 0"
     primer_bind     complement(5791..5808)
                     /locus_tag="M13-fwd"
                     /label="M13-fwd"
                     /ApEinfo_label="M13-fwd"
                     /ApEinfo_fwdcolor="cyan"
                     /ApEinfo_revcolor="green"
                     /ApEinfo_graphicformat="arrow_data {{0 1 2 0 0 -1} {} 0}
                     width 5 offset 0"
     primer_bind     8540..8560
                     /locus_tag="M13-rev"
                     /label="M13-rev"
                     /ApEinfo_label="M13-rev"
                     /ApEinfo_fwdcolor="cyan"
                     /ApEinfo_revcolor="green"
                     /ApEinfo_graphicformat="arrow_data {{0 1 2 0 0 -1} {} 0}
                     width 5 offset 0"
     primer_bind     complement(5762..5782)
                     /locus_tag="T7"
                     /label="T7"
                     /ApEinfo_label="T7"
                     /ApEinfo_fwdcolor="cyan"
                     /ApEinfo_revcolor="green"
                     /ApEinfo_graphicformat="arrow_data {{0 1 2 0 0 -1} {} 0}
                     width 5 offset 0"
     rep_origin      5966..6272
                     /locus_tag="F1 ori"
                     /label="F1 ori"
                     /ApEinfo_label="F1 ori"
                     /ApEinfo_fwdcolor="gray50"
                     /ApEinfo_revcolor="gray50"
                     /ApEinfo_graphicformat="arrow_data {{0 1 2 0 0 -1} {} 0}
                     width 5 offset 0"
     CDS             5879..5947
                     /locus_tag="LacZ alpha"
                     /label="LacZ alpha"
                     /ApEinfo_label="LacZ alpha"
                     /ApEinfo_fwdcolor="#6495ed"
                     /ApEinfo_revcolor="#6495ed"
                     /ApEinfo_graphicformat="arrow_data {{0 1 2 0 0 -1} {} 0}
                     width 5 offset 0"
     misc_binding    8512..8534
                     /locus_tag="LacO"
                     /label="LacO"
                     /ApEinfo_label="LacO"
                     /ApEinfo_fwdcolor="#6495ed"
                     /ApEinfo_revcolor="#6495ed"
                     /ApEinfo_graphicformat="arrow_data {{0 1 2 0 0 -1} {} 0}
                     width 5 offset 0"
     CDS             6733..7392
                     /locus_tag="AmpR"
                     /label="AmpR"
                     /ApEinfo_label="AmpR"
                     /ApEinfo_fwdcolor="yellow"
                     /ApEinfo_revcolor="yellow"
                     /ApEinfo_graphicformat="arrow_data {{0 1 2 0 0 -1} {} 0}
                     width 5 offset 0"
     CDS             2835..3191
                     /standard_name="MLANA_cds"
                     /locus_tag="MLANA_cds"
                     /label="MLANA_cds"
                     /ApEinfo_label="MLANA_cds"
                     /ApEinfo_fwdcolor="#e9d024"
                     /ApEinfo_revcolor="#e9d024"
                     /ApEinfo_graphicformat="arrow_data {{0 1 2 0 0 -1} {} 0}
                     width 5 offset 0"
     sig_peptide     3213..3266
                     /standard_name="T2A"
                     /note="Geneious type: LINKER"
                     /locus_tag="T2A"
                     /label="T2A"
                     /ApEinfo_label="T2A"
                     /ApEinfo_fwdcolor="cyan"
                     /ApEinfo_revcolor="green"
                     /ApEinfo_graphicformat="arrow_data {{0 1 2 0 0 -1} {} 0}
                     width 5 offset 0"
     misc_feature    2910..2939
                     /standard_name="MART-1_26-35 G6S"
                     /locus_tag="MART-1_26-35"
                     /label="MART-1_26-35"
                     /ApEinfo_label="MART-1_26-35"
                     /ApEinfo_fwdcolor="#ff7d78"
                     /ApEinfo_revcolor="#7eff74"
                     /ApEinfo_graphicformat="arrow_data {{0 1 2 0 0 -1} {} 0}
                     width 5 offset 0"
     misc_feature    3192..3212
                     /locus_tag="linker"
                     /label="linker"
                     /ApEinfo_label="linker"
                     /ApEinfo_fwdcolor="#a8eafe"
                     /ApEinfo_revcolor="green"
                     /ApEinfo_graphicformat="arrow_data {{0 1 2 0 0 -1} {} 0}
                     width 5 offset 0"
     misc_feature    2925..2927
                     /standard_name="G6S"
                     /locus_tag="G31S"
                     /label="G31S"
                     /ApEinfo_label="G31S"
                     /ApEinfo_fwdcolor="#941100"
                     /ApEinfo_revcolor="#7eff74"
                     /ApEinfo_graphicformat="arrow_data {{0 1 2 0 0 -1} {} 0}
                     width 5 offset 0"
     variation       2925..2925
                     /locus_tag="target A"
                     /label="target A"
                     /ApEinfo_label="target A"
                     /ApEinfo_fwdcolor="#ff9e39"
                     /ApEinfo_revcolor="#ff9e39"
                     /ApEinfo_graphicformat="arrow_data {{0 1 2 0 0 -1} {} 0}
                     width 5 offset 0"
     variation       2928..2928
                     /locus_tag="off-target"
                     /label="off-target"
                     /ApEinfo_label="off-target"
                     /ApEinfo_fwdcolor="#ff0617"
                     /ApEinfo_revcolor="#ff9e39"
                     /ApEinfo_graphicformat="arrow_data {{0 0.5 0 1 2 0 0 -1 0
                     -0.5} {} 0} width 5 offset 0"
ORIGIN
        1 aatgtagtct tatgcaatac tcttgtagtc ttgcaacatg gtaacgatga gttagcaaca
       61 tgccttacaa ggagagaaaa agcaccgtgc atgccgattg gtggaagtaa ggtggtacga
      121 tcgtgcctta ttaggaaggc aacagacggg tctgacatgg attggacgaa ccactgaatt
      181 gccgcattgc agagatattg tatttaagtg cctagctcga tacataaacg ggtctctctg
      241 gttagaccag atctgagcct gggagctctc tggctaacta gggaacccac tgcttaagcc
      301 tcaataaagc ttgccttgag tgcttcaagt agtgtgtgcc cgtctgttgt gtgactctgg
      361 taactagaga tccctcagac ccttttagtc agtgtggaaa atctctagca gtggcgcccg
      421 aacagggact tgaaagcgaa agggaaacca gaggagctct ctcgacgcag gactcggctt
      481 gctgaagcgc gcacggcaag aggcgagggg cggcgactgg tgagtacgcc aaaaattttg
      541 actagcggag gctagaagga gagagatggg tgcgagagcg tcagtattaa gcgggggaga
      601 attagatcgc gatgggaaaa aattcggtta aggccagggg gaaagaaaaa atataaatta
      661 aaacatatag tatgggcaag cagggagcta gaacgattcg cagttaatcc tggcctgtta
      721 gaaacatcag aaggctgtag acaaatactg ggacagctac aaccatccct tcagacagga
      781 tcagaagaac ttagatcatt atataataca gtagcaaccc tctattgtgt gcatcaaagg
      841 atagagataa aagacaccaa ggaagcttta gacaagatag aggaagagca aaacaaaagt
      901 aagaccaccg cacagcaagc ggccgctgat cttcagacct ggaggaggag atatgaggga
      961 caattggaga agtgaattat ataaatataa agtagtaaaa attgaaccat taggagtagc
     1021 acccaccaag gcaaagagaa gagtggtgca gagagaaaaa agagcagtgg gaataggagc
     1081 tttgttcctt gggttcttgg gagcagcagg aagcactatg ggcgcagcgt caatgacgct
     1141 gacggtacag gccagacaat tattgtctgg tatagtgcag cagcagaaca atttgctgag
     1201 ggctattgag gcgcaacagc atctgttgca actcacagtc tggggcatca agcagctcca
     1261 ggcaagaatc ctggctgtgg aaagatacct aaaggatcaa cagctcctgg ggatttgggg
     1321 ttgctctgga aaactcattt gcaccactgc tgtgccttgg aatgctagtt ggagtaataa
     1381 atctctggaa cagatttgga atcacacgac ctggatggag tgggacagag aaattaacaa
     1441 ttacacaagc ttaatacact ccttaattga agaatcgcaa aaccagcaag aaaagaatga
     1501 acaagaatta ttggaattag ataaatgggc aagtttgtgg aattggttta acataacaaa
     1561 ttggctgtgg tatataaaat tattcataat gatagtagga ggcttggtag gtttaagaat
     1621 agtttttgct gtactttcta tagtgaatag agttaggcag ggatattcac cattatcgtt
     1681 tcagacccac ctcccaaccc cgaggggacc cgacaggccc gaaggaatag aagaagaagg
     1741 tggagagaga gacagagaca gatccattcg attagtgaac ggatctcgac ggtatcgcta
     1801 gcttttaaaa gaaaaggggg gattgggggg tacagtgcag gggaaagaat agtagacata
     1861 atagcaacag acatacaaac taaagaatta caaaaacaaa ttacaaaaat tcaaaatttt
     1921 actagtaagg tcgggcagga agagggccta tttcccatga ttccttcata tttgcatata
     1981 cgatacaagg ctgttagaga gataattaga attaatttga ctgtaaacac aaagatatta
     2041 gtacaaaata cgtgacgtag aaagtaataa tttcttgggt agtttgcagt tttaaaatta
     2101 tgttttaaaa tggactatca tatgcttacc gtaacttgaa agtatttcga tttcttggct
     2161 ttatatatct tgtggaaagg acgaaacacc GGtccggcag taccagcagc cgTAGTgcag
     2221 cagcacgcct aggatcactg tcaggaGgcC gatcccagca agctcctttt ttgaattcca
     2281 actttgtata gaaaagttgg ggttgcgcct tttccaaggc agccctgggt ttgcgcaggg
     2341 acgcggctgc tctgggcgtg gttccgggaa acgcagcggc gccgaccctg ggtctcgcac
     2401 attcttcacg tccgttcgca gcgtcacccg gatcttcgcc gctacccttg tgggcccccc
     2461 ggcgacgctt cctgctccgc ccctaagtcg ggaaggttcc ttgcggttcg cggcgtgccg
     2521 gacgtgacaa acggaagccg cacgtctcac tagtaccctc gcagacggac agcgccaggg
     2581 agcaatggca gcgcgccgac cgcgatgggc tgtggccaat agcggctgct cagcagggcg
     2641 cgccgagagc agcggccggg aaggggcggt gcgggaggcg gggtgtgggg cggtagtgtg
     2701 ggccctgttc ctgcccgcgc ggtgttccgc attctgcaag cctccggagc gcacgtcggc
     2761 agtcggctcc ctcgttgacc gaatcaccga cctctctccc caggcaagtt tgtacaaaaa
     2821 agcaggctgc caccatgccc agggaggacg cccacttcat ctacggctac cccaagaagg
     2881 gccacggcca cagctacacc actgcagagg agcttgctgg gatcagcatc ctgacagtga
     2941 tcctaggcgt gctgctgctg atcggctgct ggtactgccg gaggaggaac ggctacaggg
     3001 ccctgatgga caagagcctg cacgtgggca cccagtgcgc cctgaccagg aggtgccccc
     3061 aggagggctt cgaccacagg gacagcaagg tgagcctcca ggagaagaac tgcgagcccg
     3121 tggtgcccaa cgcccccccc gcctacgaga agctgagcgc cgagcagagc cctccaccat
     3181 acagccccgg tggatccgga ggtgcgagcg gcgaggggag gggcagcctt cttacttgcg
     3241 gagatgtgga agagaatcca ggacccgtga gcaagggcga ggagctgttc accggggtgg
     3301 tgcccatcct ggtcgagctg gacggcgacg taaacggcca caagttcagc gtgtccggcg
     3361 agggcgaggg cgatgccacc tacggcaagc tgaccctgaa gttcatctgc accaccggca
     3421 agctgcccgt gccctggccc accctcgtga ccaccctgac ccacggcgtg cagtgcttca
     3481 gccgctaccc cgaccacatg aagcagcacg acttcttcaa gtccgccatg cccgaaggct
     3541 acgtccagga gcgcaccatc ttcttcaagg acgacggcaa ctacaagacc cgcgccgagg
     3601 tgaagttcga gggcgacacc ctggtgaacc gcatcgagct gaagggcatc gacttcaagg
     3661 aggacggcaa catcctgggg cacaagctgg agtacaactt caacagccac aacgtctata
     3721 tcatggccga caagcagaag aacggcatca aggtgaactt caagatccgc cacaacatcg
     3781 aggacggcag cgtgcagctc gccgaccact accagcagaa cacccccatc ggcgacggcc
     3841 ccgtgctgct gcccgacaac cactacctga gcacccagtc cgccctgagc aaagacccca
     3901 acgagaagcg cgatcacatg gtcctgctgg agttcgtgac cgccgccggg atcactctcg
     3961 gcatggacga gctgtacaag ggaagcggag ccacgaactt ctctctgtta aagcaagcag
     4021 gagatgttga agaaaacccc gggcctatgg ccaagccttt gtctcaagaa gaatccaccc
     4081 tcattgaaag agcaacggct acaatcaaca gcatccccat ctctgaagac tacagcgtcg
     4141 ccagcgcagc tctctctagc gacggccgca tcttcactgg tgtcaatgta tatcatttta
     4201 ctgggggacc ttgtgcagaa ctcgtggtgc tgggcactgc tgctgctgcg gcagctggca
     4261 acctgacttg tatcgtcgcg atcggaaatg agaacagggg catcttgagc ccctgcggac
     4321 ggtgccgaca ggtgcttctc gatctgcatc ctgggatcaa agccatagtg aaggacagtg
     4381 atggacagcc gacggcagtt gggattcgtg aattgctgcc ctctggttat gtgtgggagg
     4441 gctaaaccca gctttcttgt acaaagtggt ggtacccgat aatcaacctc tggattacaa
     4501 aatttgtgaa agattgactg gtattcttaa ctatgttgct ccttttacgc tatgtggata
     4561 cgctgcttta atgcctttgt atcatgctat tgcttcccgt atggctttca ttttctcctc
     4621 cttgtataaa tcctggttgc tgtctcttta tgaggagttg tggcccgttg tcaggcaacg
     4681 tggcgtggtg tgcactgtgt ttgctgacgc aacccccact ggttggggca ttgccaccac
     4741 ctgtcagctc ctttccggga ctttcgcttt ccccctccct attgccacgg cggaactcat
     4801 cgccgcctgc cttgcccgct gctggacagg ggctcggctg ttgggcactg acaattccgt
     4861 ggtgttgtcg gggaagctga cgtcctttcc atggctgctc gcctgtgttg ccacctggat
     4921 tctgcgcggg acgtccttct gctacgtccc ttcggccctc aatccagcgg accttccttc
     4981 ccgcggcctg ctgccggctc tgcggcctct tccgcgtctt cgccttcgcc ctcagacgag
     5041 tcggatctcc ctttgggccg cctccccgca tcggctttaa gaccaatgac ttacaaggca
     5101 gctgtagatc ttagccactt tttaaaagaa aaggggggac tggaagggct aattcactcc
     5161 caacgaagac aagatctgct ttttgcttgt actgggtctc tctggttaga ccagatctga
     5221 gcctgggagc tctctggcta actagggaac ccactgctta agcctcaata aagcttgcct
     5281 tgagtgcttc aagtagtgtg tgcccgtctg ttgtgtgact ctggtaacta gagatccctc
     5341 agaccctttt agtcagtgtg gaaaatctct agcagtagta gttcatgtca tcttattatt
     5401 cagtatttat aacttgcaaa gaaatgaata tcagagagtg agaggaactt gtttattgca
     5461 gcttataatg gttacaaata aagcaatagc atcacaaatt tcacaaataa agcatttttt
     5521 tcactgcatt ctagttgtgg tttgtccaaa ctcatcaatg tatcttatca tgtctggctc
     5581 tagctatccc gcccctaact ccgcccatcc cgcccctaac tccgcccagt tccgcccatt
     5641 ctccgcccca tggctgacta atttttttta tttatgcaga ggccgaggcc gcctcggcct
     5701 ctgagctatt ccagaagtag tgaggaggct tttttggagg cctagggacg tacccaattc
     5761 gccctatagt gagtcgtatt acgcgcgctc actggccgtc gttttacaac gtcgtgactg
     5821 ggaaaaccct ggcgttaccc aacttaatcg ccttgcagca catccccctt tcgccagctg
     5881 gcgtaatagc gaagaggccc gcaccgatcg cccttcccaa cagttgcgca gcctgaatgg
     5941 cgaatgggac gcgccctgta gcggcgcatt aagcgcggcg ggtgtggtgg ttacgcgcag
     6001 cgtgaccgct acacttgcca gcgccctagc gcccgctcct ttcgctttct tcccttcctt
     6061 tctcgccacg ttcgccggct ttccccgtca agctctaaat cgggggctcc ctttagggtt
     6121 ccgatttagt gctttacggc acctcgaccc caaaaaactt gattagggtg atggttcacg
     6181 tagtgggcca tcgccctgat agacggtttt tcgccctttg acgttggagt ccacgttctt
     6241 taatagtgga ctcttgttcc aaactggaac aacactcaac cctatctcgg tctattcttt
     6301 tgatttataa gggattttgc cgatttcggc ctattggtta aaaaatgagc tgatttaaca
     6361 aaaatttaac gcgaatttta acaaaatatt aacgcttaca atttaggtgg cacttttcgg
     6421 ggaaatgtgc gcggaacccc tatttgttta tttttctaaa tacattcaaa tatgtatccg
     6481 ctcatgagac aataaccctg ataaatgctt caataatatt gaaaaaggaa gagtatgagt
     6541 attcaacatt tccgtgtcgc ccttattccc ttttttgcgg cattttgcct tcctgttttt
     6601 gctcacccag aaacgctggt gaaagtaaaa gatgctgaag atcagttggg tgcacgagtg
     6661 ggttacatcg aactggatct caacagcggt aagatccttg agagttttcg ccccgaagaa
     6721 cgttttccaa tgatgagcac ttttaaagtt ctgctatgtg gcgcggtatt atcccgtatt
     6781 gacgccgggc aagagcaact cggtcgccgc atacactatt ctcagaatga cttggttgag
     6841 tactcaccag tcacagaaaa gcatcttacg gatggcatga cagtaagaga attatgcagt
     6901 gctgccataa ccatgagtga taacactgcg gccaacttac ttctgacaac gatcggagga
     6961 ccgaaggagc taaccgcttt tttgcacaac atgggggatc atgtaactcg ccttgatcgt
     7021 tgggaaccgg agctgaatga agccatacca aacgacgagc gtgacaccac gatgcctgta
     7081 gcaatggcaa caacgttgcg caaactatta actggcgaac tacttactct agcttcccgg
     7141 caacaattaa tagactggat ggaggcggat aaagttgcag gaccacttct gcgctcggcc
     7201 cttccggctg gctggtttat tgctgataaa tctggagccg gtgagcgtgg gtctcgcggt
     7261 atcattgcag cactggggcc agatggtaag ccctcccgta tcgtagttat ctacacgacg
     7321 gggagtcagg caactatgga tgaacgaaat agacagatcg ctgagatagg tgcctcactg
     7381 attaagcatt ggtaactgtc agaccaagtt tactcatata tactttagat tgatttaaaa
     7441 cttcattttt aatttaaaag gatctaggtg aagatccttt ttgataatct catgaccaaa
     7501 atcccttaac gtgagttttc gttccactga gcgtcagacc ccgtagaaaa gatcaaagga
     7561 tcttcttgag atcctttttt tctgcgcgta atctgctgct tgcaaacaaa aaaaccaccg
     7621 ctaccagcgg tggtttgttt gccggatcaa gagctaccaa ctctttttcc gaaggtaact
     7681 ggcttcagca gagcgcagat accaaatact gttcttctag tgtagccgta gttaggccac
     7741 cacttcaaga actctgtagc accgcctaca tacctcgctc tgctaatcct gttaccagtg
     7801 gctgctgcca gtggcgataa gtcgtgtctt accgggttgg actcaagacg atagttaccg
     7861 gataaggcgc agcggtcggg ctgaacgggg ggttcgtgca cacagcccag cttggagcga
     7921 acgacctaca ccgaactgag atacctacag cgtgagctat gagaaagcgc cacgcttccc
     7981 gaagagagaa aggcggacag gtatccggta agcggcaggg tcggaacagg agagcgcacg
     8041 agggagcttc cagggggaaa cgcctggtat ctttatagtc ctgtcgggtt tcgccacctc
     8101 tgacttgagc gtcgattttt gtgatgctcg tcaggggggc ggagcctatg gaaaaacgcc
     8161 agcaacgcgg cctttttacg gttcctggcc ttttgctggc cttttgctca catgttcttt
     8221 cctgcgttat cccctgattc tgtggataac cgtattaccg cctttgagtg agctgatacc
     8281 gctcgccgca gccgaacgac cgagcgcagc gagtcagtga gcgaggaagc ggaagagcgc
     8341 ccaatacgca aaccgcctct ccccgcgcgt tggccgattc attaatgcag ctggcacgac
     8401 aggtttcccg actggaaagc gggcagtgag cgcaacgcaa ttaatgtgag ttagctcact
     8461 cattaggcac cccaggcttt acactttatg cttccggctc gtatgttgtg tggaattgtg
     8521 agcggataac aatttcacac aggaaacagc tatgaccatg attacgccaa gcgcgcaatt
     8581 aaccctcact aaagggaaca aaagctggag ctgcaagctt
//
Plasmid pT_hU6 NT-SPEAR (control)_hPGK BFP-P2A-Bsd sequence (GenBank format, provided as text)
LOCUS       RP641_pT_U6_20_4        5929 bp    DNA        circular     29-JAN-2025
DEFINITION  synthetic circular DNA
ACCESSION   .
VERSION     .
KEYWORDS    .
SOURCE      synthetic DNA construct
  ORGANISM  synthetic DNA construct
REFERENCE   1  (bases 1 to 5504)
  AUTHORS   .
  TITLE     Direct Submission
  JOURNAL   Exported Jul 23, 2024 from SnapGene Viewer 7.2.1
            https://www.snapgene.com
COMMENT     [DEFINITION]: Sequence
COMMENT     ApEinfo:methylated:1
FEATURES             Location/Qualifiers
     source          join(1..452,823..877,2825..5929)
                     /note="organism="Sequence" - mol_type="other DNA" -
                     1..5504"
                     /organism="synthetic DNA construct"
                     /locus_tag="organism=Sequence - mol_type=other DNA -
                     1..5504"
                     /label="organism=Sequence - mol_type=other DNA - 1..5504"
                     /ApEinfo_label="organism=Sequence - mol_type=other DNA -
                     1..5504"
                     /ApEinfo_fwdcolor="#00DF87"
                     /ApEinfo_revcolor="#00DF87"
                     /ApEinfo_graphicformat="arrow_data {{0 0.5 0 1 2 0 0 -1 0
                     -0.5} {0 .5 .1 .5 .1 -.5 0 -.5} 0} width 5 offset 0"
                     /ApEinfo_hidden
     misc_feature    complement(join(3..452,823..877,2825..5784))
                     /note="label="lacZ_a" -
                     translation="MSIQHFRVALIPFFAAFCLPVFAHPETLVKVKDAEDQLGARVGY
                     -
                     IELDLNSGKILESFRPEERFPMMSTFKVLLCGAVLSRIDAGQEQLGRRIHYSQNDLVE
                     -
                     YSPVTEKHLTDGMTVRELCSAAITMSDNTAANLLLTTIGGPKELTAFLHNMGDHVTRL
                     -
                     DRWEPELNEAIPNDERDTTMPVAMATTLRKLLTGELLTLASRQQLIDWMEADKVAG"
                     /locus_tag="lacZ_a"
                     /label="lacZ_a"
                     /ApEinfo_label="lacZ_a"
                     /ApEinfo_fwdcolor="#7eff74"
                     /ApEinfo_revcolor="#7eff74"
                     /ApEinfo_graphicformat="arrow_data {{0 0.5 0 1 2 0 0 -1 0
                     -0.5} {0 .5 .1 .5 .1 -.5 0 -.5} 0} width 5 offset 0"
                     /ApEinfo_hidden
     misc_feature    22..53
                     /locus_tag="LOR"
                     /label="LOR"
                     /ApEinfo_label="LOR"
                     /ApEinfo_fwdcolor="#7eff74"
                     /ApEinfo_revcolor="#7eff74"
                     /ApEinfo_graphicformat="arrow_data {{0 0.5 0 1 2 0 0 -1 0
                     -0.5} {0 .5 .1 .5 .1 -.5 0 -.5} 0} width 5 offset 0"
     promoter        710..716
                     /vntifkey="29"
                     /locus_tag="hU6"
                     /label="hU6"
                     /ApEinfo_label="hU6"
                     /ApEinfo_fwdcolor="#346ee0"
                     /ApEinfo_revcolor="#346ee0"
                     /ApEinfo_graphicformat="arrow_data {{0 1 2 0 0 -1} {} 0}
                     width 5 offset 0"
     misc_feature    219..250
                     /locus_tag="LIR"
                     /label="LIR"
                     /ApEinfo_label="LIR"
                     /ApEinfo_fwdcolor="#7eff74"
                     /ApEinfo_revcolor="#7eff74"
                     /ApEinfo_graphicformat="arrow_data {{0 0.5 0 1 2 0 0 -1 0
                     -0.5} {0 .5 .1 .5 .1 -.5 0 -.5} 0} width 5 offset 0"
     misc_feature    complement(382..402)
                     /note="label="pBABE_3_primer" - complement(382..402)"
                     /locus_tag="pBABE_3_primer"
                     /label="pBABE_3_primer"
                     /ApEinfo_label="pBABE_3_primer"
                     /ApEinfo_fwdcolor="#7eff74"
                     /ApEinfo_revcolor="#7eff74"
                     /ApEinfo_graphicformat="arrow_data {{0 0.5 0 1 2 0 0 -1 0
                     -0.5} {0 .5 .1 .5 .1 -.5 0 -.5} 0} width 5 offset 0"
     promoter        453..709
                     /vntifkey="29"
                     /locus_tag="hU6(1)"
                     /label="hU6(1)"
                     /ApEinfo_label="hU6"
                     /ApEinfo_fwdcolor="#346ee0"
                     /ApEinfo_revcolor="#346ee0"
                     /ApEinfo_graphicformat="arrow_data {{0 1 2 0 0 -1} {} 0}
                     width 5 offset 0"
     ORF             1418..2134
                     /application_notes="Low fluorescence and low
                     photostability."
                     /description="Blue variant of EGFP generated by
                     mutagenesis"
                     /full_name="Enhanced blue fluorescent protein(ns)"
                     /note="color: #6ddaae; direction: RIGHT"
                     /official_designation="EBFP(ns)"
                     /uuid="c22d3886879b4745b895a6f25c150792"
                     /locus_tag="EBFP(ns)"
                     /label="EBFP(ns)"
                     /ApEinfo_label="EBFP(ns)"
                     /ApEinfo_fwdcolor="#0f7ffe"
                     /ApEinfo_revcolor="pink"
                     /ApEinfo_graphicformat="arrow_data {{0 1 2 0 0 -1} {} 0}
                     width 5 offset 0"
     primer_bind     complement(2835..2856)
                     /locus_tag="pT3_rev"
                     /label="pT3_rev"
                     /ApEinfo_label="pT3_rev"
                     /ApEinfo_fwdcolor="#14c0bd"
                     /ApEinfo_revcolor="#4ec02b"
                     /ApEinfo_graphicformat="arrow_data {{0 0.5 0 1 2 0 0 -1 0
                     -0.5} {0 .5 .1 .5 .1 -.5 0 -.5} 0} width 5 offset 0"
     misc_feature    2845..2864
                     /note="label="EBV_rev_primer" -
                     translation="MTEYKLVVVGAVGVGKSALTIQLIQNHFVDEYDPTIEDSYRKQV
                     -
                     VIDGETCLLDILDTAGQEEYSAMRDQYMRTGEGFLCVFAINNTKSFEDIHHYREQIKR
                     -
                     VKDSEDVPMVLVGNKCDLPSRTVDTKQAQELARSYGIPFIETSAKTRQGVDDAFYTLV
                     - REIRKHKEKMSKDGKKKKKKSRTRCTVM*" - 2420..2439"
                     /locus_tag="EBV_rev_primer"
                     /label="EBV_rev_primer"
                     /ApEinfo_label="EBV_rev_primer"
                     /ApEinfo_fwdcolor="#7eff74"
                     /ApEinfo_revcolor="#7eff74"
                     /ApEinfo_graphicformat="arrow_data {{0 0.5 0 1 2 0 0 -1 0
                     -0.5} {0 .5 .1 .5 .1 -.5 0 -.5} 0} width 5 offset 0"
     Linker          2135..2200
                     /application_notes="Causes co-translational cleavage of
                     the encoded polypeptide. Multiple proteins can be made
                     from a polycistronic transcript containing multiple ORFs
                     separated by 2A. P2A and T2A have higher cleavage
                     efficiency compared to other 2As."
                     /description="Self-cleaving 2A peptide from Porcine
                     teschovirus-1"
                     /full_name="Porcine teschovirus-1 2A peptide"
                     /note="color: #1f36a9; direction: RIGHT"
                     /official_designation="P2A"
                     /uuid="26656daa74294a0f8b331614d72d5f5a"
                     /locus_tag="P2A"
                     /label="P2A"
                     /ApEinfo_label="P2A"
                     /ApEinfo_fwdcolor="#20fefe"
                     /ApEinfo_revcolor="pink"
                     /ApEinfo_graphicformat="arrow_data {{0 1 2 0 0 -1} {} 0}
                     width 5 offset 0"
     primer_bind     3004..3025
                     /locus_tag="RIR Primer"
                     /label="RIR Primer"
                     /ApEinfo_label="RIR Primer"
                     /ApEinfo_fwdcolor="#14c0bd"
                     /ApEinfo_revcolor="#4ec02b"
                     /ApEinfo_graphicformat="arrow_data {{0 0.5 0 1 2 0 0 -1 0
                     -0.5} {0 .5 .1 .5 .1 -.5 0 -.5} 0} width 5 offset 0"
     promoter        883..1387
                     /__SeqFeature__="True"
                     /__level__="0"
                     /application_notes="Medium-strength promoter."
                     /description="['Human phosphoglycerate kinase 1
                     promoter']"
                     /name="hPGK promoter"
                     /note="color: #5566f5; direction: RIGHT"
                     /official_designation="hPGK promoter"
                     /uuid="a105ee8f48814e769c9a41c38cfff41e"
                     /vntifkey="21"
                     /locus_tag="hPGK promoter"
                     /label="hPGK promoter"
                     /ApEinfo_label="hPGK promoter"
                     /ApEinfo_fwdcolor="#346ee0"
                     /ApEinfo_revcolor="#346ee0"
                     /ApEinfo_graphicformat="arrow_data {{0 1 2 0 0 -1} {} 0}
                     width 5 offset 0"
     misc_feature    3032..3062
                     /locus_tag="RIR"
                     /label="RIR"
                     /ApEinfo_label="RIR"
                     /ApEinfo_fwdcolor="#7eff74"
                     /ApEinfo_revcolor="#7eff74"
                     /ApEinfo_graphicformat="arrow_data {{0 0.5 0 1 2 0 0 -1 0
                     -0.5} {0 .5 .1 .5 .1 -.5 0 -.5} 0} width 5 offset 0"
     CDS             2201..2599
                     /__SeqFeature__="True"
                     /__level__="0"
                     /application_notes="Allows cells to be resistant to
                     blasticidin."
                     /description="['Blasticidin resistance gene']"
                     /full_name="Blasticidin resistance gene"
                     /name="Bsd"
                     /note="color: #68e66d; direction: RIGHT"
                     /official_designation="Bsd"
                     /uuid="70e7f83c244f451dae86de92afa03717"
                     /vntifkey="21"
                     /locus_tag="Bsd"
                     /label="Bsd"
                     /ApEinfo_label="Bsd"
                     /ApEinfo_fwdcolor="#e9d024"
                     /ApEinfo_revcolor="#e9d024"
                     /ApEinfo_graphicformat="arrow_data {{0 1 2 0 0 -1} {} 0}
                     width 5 offset 0"
     misc_feature    3229..3260
                     /locus_tag="ROR"
                     /label="ROR"
                     /ApEinfo_label="ROR"
                     /ApEinfo_fwdcolor="#7eff74"
                     /ApEinfo_revcolor="#7eff74"
                     /ApEinfo_graphicformat="arrow_data {{0 0.5 0 1 2 0 0 -1 0
                     -0.5} {0 .5 .1 .5 .1 -.5 0 -.5} 0} width 5 offset 0"
     promoter        complement(3309..3327)
                     /note="label="M13_reverse_primer" -
                     translation="MTEYKLVVVGAVGVGKSALTIQLIQNHFVDEYDPTIEDSYRKQV
                     -
                     VIDGETCLLDILDTAGQEEYSAMRDQYMRTGEGFLCVFAINNTKSFEDIHHYREQIKR
                     -
                     VKDSEDVPMVLVGNKCDLPSRTVDTKQAQELARSYGIPFIETSAKTRQGVDDAFYTLV
                     - REIRKHKEKMSKDGKKKKKKSRTRCTVM*" - complement("
                     /locus_tag="M13_reverse_primer"
                     /label="M13_reverse_primer"
                     /ApEinfo_label="M13_reverse_primer"
                     /ApEinfo_fwdcolor="#346ee0"
                     /ApEinfo_revcolor="#346ee0"
                     /ApEinfo_graphicformat="arrow_data {{0 0.5 0 1 2 0 0 -1 0
                     -0.5} {0 .5 .1 .5 .1 -.5 0 -.5} 0} width 5 offset 0"
     primer_bind     complement(3313..3329)
                     /note="common sequencing primer, one of multiple similar
                     variants"
                     /locus_tag="M13 rev"
                     /label="M13 rev"
                     /ApEinfo_label="M13 rev"
                     /ApEinfo_fwdcolor="#14c0bd"
                     /ApEinfo_revcolor="#4ec02b"
                     /ApEinfo_graphicformat="arrow_data {{0 0.5 0 1 2 0 0 -1 0
                     -0.5} {0 .5 .1 .5 .1 -.5 0 -.5} 0} width 5 offset 0"
     misc_feature    complement(3326..3348)
                     /note="label="M13_pUC_rev_primer" -
                     translation="MTEYKLVVVGAVGVGKSALTIQLIQNHFVDEYDPTIEDSYRKQV
                     -
                     VIDGETCLLDILDTAGQEEYSAMRDQYMRTGEGFLCVFAINNTKSFEDIHHYREQIKR
                     -
                     VKDSEDVPMVLVGNKCDLPSRTVDTKQAQELARSYGIPFIETSAKTRQGVDDAFYTLV
                     - REIRKHKEKMSKDGKKKKKKSRTRCTVM*" - complement("
                     /locus_tag="M13_pUC_rev_primer"
                     /label="M13_pUC_rev_primer"
                     /ApEinfo_label="M13_pUC_rev_primer"
                     /ApEinfo_fwdcolor="#7eff74"
                     /ApEinfo_revcolor="#7eff74"
                     /ApEinfo_graphicformat="arrow_data {{0 0.5 0 1 2 0 0 -1 0
                     -0.5} {0 .5 .1 .5 .1 -.5 0 -.5} 0} width 5 offset 0"
     protein_bind    3337..3353
                     /bound_moiety="lac repressor encoded by lacI"
                     /note="The lac repressor binds to the lac operator to
                     inhibit transcription in E. coli. This inhibition can be
                     relieved by adding lactose or
                     isopropyl-beta-D-thiogalactopyranoside (IPTG)."
                     /locus_tag="lac operator"
                     /label="lac operator"
                     /ApEinfo_label="lac operator"
                     /ApEinfo_fwdcolor="#00D95A"
                     /ApEinfo_revcolor="#00D95A"
                     /ApEinfo_graphicformat="arrow_data {{0 0.5 0 1 2 0 0 -1 0
                     -0.5} {0 .5 .1 .5 .1 -.5 0 -.5} 0} width 5 offset 0"
     promoter        complement(3361..3391)
                     /note="promoter for the E. coli lac operon"
                     /locus_tag="lac promoter"
                     /label="lac promoter"
                     /ApEinfo_label="lac promoter"
                     /ApEinfo_fwdcolor="#346ee0"
                     /ApEinfo_revcolor="#346ee0"
                     /ApEinfo_graphicformat="arrow_data {{0 0.5 0 1 2 0 0 -1 0
                     -0.5} {0 .5 .1 .5 .1 -.5 0 -.5} 0} width 5 offset 0"
     promoter        complement(3362..3391)
                     /note="label="lac_promoter" -
                     translation="MTEYKLVVVGAVGVGKSALTIQLIQNHFVDEYDPTIEDSYRKQV
                     -
                     VIDGETCLLDILDTAGQEEYSAMRDQYMRTGEGFLCVFAINNTKSFEDIHHYREQIKR
                     -
                     VKDSEDVPMVLVGNKCDLPSRTVDTKQAQELARSYGIPFIETSAKTRQGVDDAFYTLV
                     - REIRKHKEKMSKDGKKKKKKSRTRCTVM*" - complement(2937.."
                     /locus_tag="lac_promoter"
                     /label="lac_promoter"
                     /ApEinfo_label="lac_promoter"
                     /ApEinfo_fwdcolor="#346ee0"
                     /ApEinfo_revcolor="#346ee0"
                     /ApEinfo_graphicformat="arrow_data {{0 0.5 0 1 2 0 0 -1 0
                     -0.5} {0 .5 .1 .5 .1 -.5 0 -.5} 0} width 5 offset 0"
     protein_bind    3406..3427
                     /bound_moiety="E. coli catabolite activator protein"
                     /note="CAP binding activates transcription in the presence
                     of cAMP."
                     /locus_tag="CAP binding site"
                     /label="CAP binding site"
                     /ApEinfo_label="CAP binding site"
                     /ApEinfo_fwdcolor="#008065"
                     /ApEinfo_revcolor="#008065"
                     /ApEinfo_graphicformat="arrow_data {{0 0.5 0 1 2 0 0 -1 0
                     -0.5} {0 .5 .1 .5 .1 -.5 0 -.5} 0} width 5 offset 0"
     source          complement(2600..2824)
                     /locus_tag="source"
                     /label="source"
                     /ApEinfo_label="source"
                     /ApEinfo_fwdcolor="#00DF87"
                     /ApEinfo_revcolor="#00DF87"
                     /ApEinfo_graphicformat="arrow_data {{0 0.5 0 1 2 0 0 -1 0
                     -0.5} {0 .5 .1 .5 .1 -.5 0 -.5} 0} width 5 offset 0"
                     /ApEinfo_hidden
     rep_origin      complement(3715..4303)
                     /direction=LEFT
                     /note="high-copy-number ColE1/pMB1/pBR322/pUC origin of
                     replication"
                     /locus_tag="ori"
                     /label="ori"
                     /ApEinfo_label="ori"
                     /ApEinfo_fwdcolor="#999999"
                     /ApEinfo_revcolor="#999999"
                     /ApEinfo_graphicformat="arrow_data {{0 0.5 0 1 2 0 0 -1 0
                     -0.5} {0 .5 .1 .5 .1 -.5 0 -.5} 0} width 5 offset 0"
     gene            complement(4474..5334)
                     /note="label="Ampicillin" - gene="Ampicillin" -
                     translation="MTEYKLVVVGAVGVGKSALTIQLIQNHFVDEYDPTIEDSYRKQV
                     -
                     VIDGETCLLDILDTAGQEEYSAMRDQYMRTGEGFLCVFAINNTKSFEDIHHYREQIKR
                     -
                     VKDSEDVPMVLVGNKCDLPSRTVDTKQAQELARSYGIPFIETSAKTRQGVDDAFYTLV
                     - REIRKHKEKMSKDGKKKKKKSRTRCTVM*" -"
                     /locus_tag="AmpR"
                     /label="AmpR"
                     /ApEinfo_label="AmpR"
                     /ApEinfo_fwdcolor="#ff797d"
                     /ApEinfo_revcolor="#ff102f"
                     /ApEinfo_graphicformat="arrow_data {{0 0.5 0 1 2 0 0 -1 0
                     -0.5} {0 .5 .1 .5 .1 -.5 0 -.5} 0} width 5 offset 0"
     CDS             complement(4474..5334)
                     /note="label="ORF frame 2" -
                     translation="MSIQHFRVALIPFFAAFCLPVFAHPETLVKVKDAEDQLGARVGY
                     -
                     IELDLNSGKILESFRPEERFPMMSTFKVLLCGAVLSRIDAGQEQLGRRIHYSQNDLVE
                     -
                     YSPVTEKHLTDGMTVRELCSAAITMSDNTAANLLLTTIGGPKELTAFLHNMGDHVTRL
                     - DRWEPELNEAIPNDERDTTMPVAMATTLRKLLTGELLTLASRQQLIDWMEA"
                     /locus_tag="ORF frame 2"
                     /label="ORF frame 2"
                     /ApEinfo_label="ORF frame 2"
                     /ApEinfo_fwdcolor="#e9d024"
                     /ApEinfo_revcolor="#e9d024"
                     /ApEinfo_graphicformat="arrow_data {{0 0.5 0 1 2 0 0 -1 0
                     -0.5} {0 .5 .1 .5 .1 -.5 0 -.5} 0} width 5 offset 0"
     misc_binding    739..742
                     /locus_tag="4nt mismatch (as in Uzonyi et al 2021)"
                     /label="4nt mismatch (as in Uzonyi et al 2021)"
                     /ApEinfo_label="4nt mismatch (as in Uzonyi et al 2021)"
                     /ApEinfo_fwdcolor="#ff4809"
                     /ApEinfo_revcolor="#ff4809"
                     /ApEinfo_graphicformat="arrow_data {{0 1 2 0 0 -1} {} 0}
                     width 5 offset 0"
     promoter        complement(5335..5439)
                     /gene="bla"
                     /locus_tag="AmpR promoter"
                     /label="AmpR promoter"
                     /ApEinfo_label="AmpR promoter"
                     /ApEinfo_fwdcolor="#346ee0"
                     /ApEinfo_revcolor="#346ee0"
                     /ApEinfo_graphicformat="arrow_data {{0 0.5 0 1 2 0 0 -1 0
                     -0.5} {0 .5 .1 .5 .1 -.5 0 -.5} 0} width 5 offset 0"
     promoter        complement(5376..5404)
                     /note="label="AmpR_promoter" -
                     translation="MSIQHFRVALIPFFAAFCLPVFAHPETLVKVKDAEDQLGARVGY
                     -
                     IELDLNSGKILESFRPEERFPMMSTFKVLLCGAVLSRIDAGQEQLGRRIHYSQNDLVE
                     -
                     YSPVTEKHLTDGMTVRELCSAAITMSDNTAANLLLTTIGGPKELTAFLHNMGDHVTRL
                     - DRWEPELNEAIPNDERDTTMPVAMATTLRKLLTGELLTLASRQQLIDWM"
                     /locus_tag="AmpR_promoter"
                     /label="AmpR_promoter"
                     /ApEinfo_label="AmpR_promoter"
                     /ApEinfo_fwdcolor="#346ee0"
                     /ApEinfo_revcolor="#346ee0"
                     /ApEinfo_graphicformat="arrow_data {{0 0.5 0 1 2 0 0 -1 0
                     -0.5} {0 .5 .1 .5 .1 -.5 0 -.5} 0} width 5 offset 0"
     misc_feature    complement(5563..5585)
                     /note="label="pGEX_3_primer" -
                     translation="MSIQHFRVALIPFFAAFCLPVFAHPETLVKVKDAEDQLGARVGY
                     -
                     IELDLNSGKILESFRPEERFPMMSTFKVLLCGAVLSRIDAGQEQLGRRIHYSQNDLVE
                     -
                     YSPVTEKHLTDGMTVRELCSAAITMSDNTAANLLLTTIGGPKELTAFLHNMGDHVTRL
                     - DRWEPELNEAIPNDERDTTMPVAMATTLRKLLTGELLTLASRQQLIDWM"
                     /locus_tag="pGEX_3_primer"
                     /label="pGEX_3_primer"
                     /ApEinfo_label="pGEX_3_primer"
                     /ApEinfo_fwdcolor="#7eff74"
                     /ApEinfo_revcolor="#7eff74"
                     /ApEinfo_graphicformat="arrow_data {{0 0.5 0 1 2 0 0 -1 0
                     -0.5} {0 .5 .1 .5 .1 -.5 0 -.5} 0} width 5 offset 0"
     misc_feature    2600..2824
                     /__SeqFeature__="True"
                     /__level__="0"
                     /application_notes="Allows transcription termination and
                     polyadenylation of mRNA transcribed by Pol II RNA
                     polymerase."
                     /full_name="Bovine growth hormone polyadenylation"
                     /name="BGH pA"
                     /note="color: #46c6ef; direction: LEFT"
                     /official_designation="BGH pA"
                     /uuid="439e92d8cb8e406dbfacd87ce28d14dc"
                     /vntifkey="21"
                     /locus_tag="BGH pA"
                     /label="BGH pA"
                     /ApEinfo_label="BGH pA"
                     /ApEinfo_fwdcolor="#7eff74"
                     /ApEinfo_revcolor="#7eff74"
                     /ApEinfo_graphicformat="arrow_data {{0 0.5 0 1 2 0 0 -1 0
                     -0.5} {0 .5 .1 .5 .1 -.5 0 -.5} 0} width 5 offset 0"
     misc_feature    5898..5920
                     /note="label="M13_pUC_fwd_primer" -
                     translation="MSIQHFRVALIPFFAAFCLPVFAHPETLVKVKDAEDQLGARVGY
                     -
                     IELDLNSGKILESFRPEERFPMMSTFKVLLCGAVLSRIDAGQEQLGRRIHYSQNDLVE
                     -
                     YSPVTEKHLTDGMTVRELCSAAITMSDNTAANLLLTTIGGPKELTAFLHNMGDHVTRL
                     - DRWEPELNEAIPNDERDTTMPVAMATTLRKLLTGELLTLASRQQ"
                     /locus_tag="M13_pUC_fwd_primer"
                     /label="M13_pUC_fwd_primer"
                     /ApEinfo_label="M13_pUC_fwd_primer"
                     /ApEinfo_fwdcolor="#7eff74"
                     /ApEinfo_revcolor="#7eff74"
                     /ApEinfo_graphicformat="arrow_data {{0 0.5 0 1 2 0 0 -1 0
                     -0.5} {0 .5 .1 .5 .1 -.5 0 -.5} 0} width 5 offset 0"
     promoter        5913..5929
                     /note="label="M13_forward20_primer" -
                     translation="MSIQHFRVALIPFFAAFCLPVFAHPETLVKVKDAEDQLGARVGY
                     -
                     IELDLNSGKILESFRPEERFPMMSTFKVLLCGAVLSRIDAGQEQLGRRIHYSQNDLVE
                     -
                     YSPVTEKHLTDGMTVRELCSAAITMSDNTAANLLLTTIGGPKELTAFLHNMGDHVTRL
                     - DRWEPELNEAIPNDERDTTMPVAMATTLRKLLTGELLTLASR"
                     /locus_tag="M13_forward20_primer"
                     /label="M13_forward20_primer"
                     /ApEinfo_label="M13_forward20_primer"
                     /ApEinfo_fwdcolor="#346ee0"
                     /ApEinfo_revcolor="#346ee0"
                     /ApEinfo_graphicformat="arrow_data {{0 0.5 0 1 2 0 0 -1 0
                     -0.5} {0 .5 .1 .5 .1 -.5 0 -.5} 0} width 5 offset 0"
     primer_bind     5913..5929
                     /note="common sequencing primer, one of multiple similar
                     variants"
                     /locus_tag="M13 fwd"
                     /label="M13 fwd"
                     /ApEinfo_label="M13 fwd"
                     /ApEinfo_fwdcolor="#14c0bd"
                     /ApEinfo_revcolor="#4ec02b"
                     /ApEinfo_graphicformat="arrow_data {{0 0.5 0 1 2 0 0 -1 0
                     -0.5} {0 .5 .1 .5 .1 -.5 0 -.5} 0} width 5 offset 0"
     terminator      793..798
                     /locus_tag="PolIII terminator"
                     /label="PolIII terminator"
                     /ApEinfo_label="PolIII terminator"
                     /ApEinfo_fwdcolor="#9d1b1c"
                     /ApEinfo_revcolor="#9d1b1c"
                     /ApEinfo_graphicformat="arrow_data {{0 1 2 0 0 -1} {} 0}
                     width 5 offset 0"
ORIGIN
        1 gaattcgagc tcggtaccct acagttgaag tcggaagttt acatacactt aagttggagt
       61 cattaaaact cgtttttcaa ctactccaca aatttcttgt taacaaacaa tagttttggc
      121 aagtcagtta ggacatctac tttgtgcatg acacaagtca tttttccaac aattgtttac
      181 agacagatta tttcacttat aattcactgt atcacaattc cagtgggtca gaagtttaca
      241 tacactaagt tgactgtgcc tttaaacagc ttggaaaatt ccagaaaatg atgtcatggc
      301 tttagaagct tctgatagac taattgacat catttgagtc aattggaggt gtacctgtgg
      361 atgtatttca aggaattctg tggaatgtgt gtcagttagg gtgtggaaag tccccaggct
      421 ccccagcagg cagaagtatg caaagcatgc ataaggtcgg gcaggaagag ggcctatttc
      481 ccatgattcc ttcatatttg catatacgat acaaggctgt tagagagata attagaatta
      541 atttgactgt aaacacaaag atattagtac aaaatacgtg acgtagaaag taataatttc
      601 ttgggtagtt tgcagtttta aaattatgtt ttaaaatgga ctatcatatg cttaccgtaa
      661 cttgaaagta tttcgatttc ttggctttat atatcttgtg gaaaggacga aacaccGGtc
      721 ggggtacctg ctgaagcaaa cgacgccgta tgtcagggta gtgacaagtg ttggcCatgg
      781 aacaggtagt ttttttttct agacccagct ttcttgtaca aactcgagaa taaaggtacc
      841 ttattttcat tagatttgtg tgttggtttt ttgtgtgCTA GCgggttgcg ccttttccaa
      901 ggcagccctg ggtttgcgca gggacgcggc tgctctgggc gtggttccgg gaaacgcagc
      961 ggcgccgacc ctgggtctcg cacattcttc acgtccgttc gcagcgtcac ccggatcttc
     1021 gccgctaccc ttgtgggccc cccggcgacg cttcctgctc cgcccctaag tcgggaaggt
     1081 tccttgcggt tcgcggcgtg ccggacgtga caaacggaag ccgcacgtct cactagtacc
     1141 ctcgcagacg gacagcgcca gggagcaatg gcagcgcgcc gaccgcgatg ggctgtggcc
     1201 aatagcggct gctcagcagg gcgcgccgag agcagcggcc gggaaggggc ggtgcgggag
     1261 gcggggtgtg gggcggtagt gtgggccctg ttcctgcccg cgcggtgttc cgcattctgc
     1321 aagcctccgg agcgcacgtc ggcagtcggc tccctcgttg accgaatcac cgacctctct
     1381 ccccaggcaa gtttgtacaa aaaagcaggc tgccaccatg gtgagcaagg gcgaggagct
     1441 gttcaccggg gtggtgccca tcctggtcga gctggacggc gacgtaaacg gccacaagtt
     1501 cagcgtgtcc ggcgagggcg agggcgatgc cacctacggc aagctgaccc tgaagttcat
     1561 ctgcaccacc ggcaagctgc ccgtgccctg gcccaccctc gtgaccaccc tgacccacgg
     1621 cgtgcagtgc ttcagccgct accccgacca catgaagcag cacgacttct tcaagtccgc
     1681 catgcccgaa ggctacgtcc aggagcgcac catcttcttc aaggacgacg gcaactacaa
     1741 gacccgcgcc gaggtgaagt tcgagggcga caccctggtg aaccgcatcg agctgaaggg
     1801 catcgacttc aaggaggacg gcaacatcct ggggcacaag ctggagtaca acttcaacag
     1861 ccacaacgtc tatatcatgg ccgacaagca gaagaacggc atcaaggtga acttcaagat
     1921 ccgccacaac atcgaggacg gcagcgtgca gctcgccgac cactaccagc agaacacccc
     1981 catcggcgac ggccccgtgc tgctgcccga caaccactac ctgagcaccc agtccgccct
     2041 gagcaaagac cccaacgaga agcgcgatca catggtcctg ctggagttcg tgaccgccgc
     2101 cgggatcact ctcggcatgg acgagctgta caagggaagc ggagccacga acttctctct
     2161 gttaaagcaa gcaggagatg ttgaagaaaa ccccgggcct atggccaagc ctttgtctca
     2221 agaagaatcc accctcattg aaagagcaac ggctacaatc aacagcatcc ccatctctga
     2281 agactacagc gtcgccagcg cagctctctc tagcgacggc cgcatcttca ctggtgtcaa
     2341 tgtatatcat tttactgggg gaccttgtgc agaactcgtg gtgctgggca ctgctgctgc
     2401 tgcggcagct ggcaacctga cttgtatcgt cgcgatcgga aatgagaaca ggggcatctt
     2461 gagcccctgc ggacggtgcc gacaggtgct tctcgatctg catcctggga tcaaagccat
     2521 agtgaaggac agtgatggac agccgacggc agttgggatt cgtgaattgc tgccctctgg
     2581 ttatgtgtgg gagggctaac tgtgccttct agttgccagc catctgttgt ttgcccctcc
     2641 cccgtgcctt ccttgaccct ggaaggtgcc actcccactg tcctttccta ataaaatgag
     2701 gaaattgcat cgcattgtct gagtaggtgt cattctattc tggggggtgg ggtggggcag
     2761 gacagcaagg gggaggattg ggaagacaat agcaggcatg ctggggatgc ggtgggctct
     2821 atgggcggcc gctgcattct agttgtggtt tgtccaaact catcaatgta tcttatcatg
     2881 tctggatccc atcacaaagc tctgacctca atcctataga aaggaggaat gagccaaaat
     2941 tcacccaact tattgtggga agcttgtgga aggctactcg aaatgtttga cccaagttaa
     3001 acaatttaaa ggcaatgcta ccaaatacta attgagtgta tgttaacttc tgacccactg
     3061 ggaatgtgat gaaagaaata aaagctgaaa tgaatcattc tctctactat tattctgata
     3121 tttcacattc ttaaaataaa gtggtgatcc taactgacct taagacaggg aatctttact
     3181 cggattaaat gtcaggaatt gtgaaaaagt gagtttaaat gtatttggct aaggtgtatg
     3241 taaacttccg acttcaactg taggggatcc tctagagtcg acctgcaggc atgcaagctt
     3301 ggcgtaatca tggtcatagc tgtttcctgt gtgaaattgt tatccgctca caattccaca
     3361 caacatacga gccggaagca taaagtgtaa agcctggggt gcctaatgag tgagctaact
     3421 cacattaatt gcgttgcgct cactgcccgc tttccagtcg ggaaacctgt cgtgccagct
     3481 gcattaatga atcggccaac gcgcggggag aggcggtttg cgtattgggc gctcttccgc
     3541 ttcctcgctc actgactcgc tgcgctcggt cgttcggctg cggcgagcgg tatcagctca
     3601 ctcaaaggcg gtaatacggt tatccacaga atcaggggat aacgcaggaa agaacatgtg
     3661 agcaaaaggc cagcaaaagg ccaggaaccg taaaaaggcc gcgttgctgg cgtttttcca
     3721 taggctccgc ccccctgacg agcatcacaa aaatcgacgc tcaagtcaga ggtggcgaaa
     3781 cccgacagga ctataaagat accaggcgtt tccccctgga agctccctcg tgcgctctcc
     3841 tgttccgacc ctgccgctta ccggatacct gtccgccttt ctcccttcgg gaagcgtggc
     3901 gctttctcat agctcacgct gtaggtatct cagttcggtg taggtcgttc gctccaagct
     3961 gggctgtgtg cacgaacccc ccgttcagcc cgaccgctgc gccttatccg gtaactatcg
     4021 tcttgagtcc aacccggtaa gacacgactt atcgccactg gcagcagcca ctggtaacag
     4081 gattagcaga gcgaggtatg taggcggtgc tacagagttc ttgaagtggt ggcctaacta
     4141 cggctacact agaagaacag tatttggtat ctgcgctctg ctgaagccag ttaccttcgg
     4201 aaaaagagtt ggtagctctt gatccggcaa acaaaccacc gctggtagcg gtggtttttt
     4261 tgtttgcaag cagcagatta cgcgcagaaa aaaaggatct caagaagatc ctttgatctt
     4321 ttctacgggg tctgacgctc agtggaacga aaactcacgt taagggattt tggtcatgag
     4381 attatcaaaa aggatcttca cctagatcct tttaaattaa aaatgaagtt ttaaatcaat
     4441 ctaaagtata tatgagtaaa cttggtctga cagttaccaa tgcttaatca gtgaggcacc
     4501 tatctcagcg atctgtctat ttcgttcatc catagttgcc tgactccccg tcgtgtagat
     4561 aactacgata cgggagggct taccatctgg ccccagtgct gcaatgatac cgcgagaccc
     4621 acgctcaccg gctccagatt tatcagcaat aaaccagcca gccggaaggg ccgagcgcag
     4681 aagtggtcct gcaactttat ccgcctccat ccagtctatt aattgttgcc gggaagctag
     4741 agtaagtagt tcgccagtta atagtttgcg caacgttgtt gccattgcta caggcatcgt
     4801 ggtgtcacgc tcgtcgtttg gtatggcttc attcagctcc ggttcccaac gatcaaggcg
     4861 agttacatga tcccccatgt tgtgcaaaaa agcggttagc tccttcggtc ctccgatcgt
     4921 tgtcagaagt aagttggccg cagtgttatc actcatggtt atggcagcac tgcataattc
     4981 tcttactgtc atgccatccg taagatgctt ttctgtgact ggtgagtact caaccaagtc
     5041 attctgagaa tagtgtatgc ggcgaccgag ttgctcttgc ccggcgtcaa tacgggataa
     5101 taccgcgcca catagcagaa ctttaaaagt gctcatcatt ggaaaacgtt cttcggggcg
     5161 aaaactctca aggatcttac cgctgttgag atccagttcg atgtaaccca ctcgtgcacc
     5221 caactgatct tcagcatctt ttactttcac cagcgtttct gggtgagcaa aaacaggaag
     5281 gcaaaatgcc gcaaaaaagg gaataagggc gacacggaaa tgttgaatac tcatactctt
     5341 cctttttcaa tattattgaa gcatttatca gggttattgt ctcatgagcg gatacatatt
     5401 tgaatgtatt tagaaaaata aacaaatagg ggttccgcgc acatttcccc gaaaagtgcc
     5461 acctgacgtc taagaaacca ttattatcat gacattaacc tataaaaata ggcgtatcac
     5521 gaggcccttt cgtctcgcgc gtttcggtga tgacggtgaa aacctctgac acatgcagct
     5581 cccggagacg gtcacagctt gtctgtaagc ggatgccggg agcagacaag cccgtcaggg
     5641 cgcgtcagcg ggtgttggcg ggtgtcgggg ctggcttaac tatgcggcat cagagcagat
     5701 tgtactgaga gtgcaccata tgcggtgtga aataccgcac agatgcgtaa ggagaaaata
     5761 ccgcatcagg cgccattcgc cattcaggct gcgcaactgt tgggaagggc gatcggtgcg
     5821 ggcctcttcg ctattacgcc agctggcgaa agggggatgt gctgcaaggc gattaagttg
     5881 ggtaacgcca gggttttccc agtcacgacg ttgtaaaacg acggccagt
//
```

**Plasmid pT_hU6 T-SPEAR (MART-1^G31S^)_hPGK BFP-P2A-Bsd sequence (GenBank format, provided as text)**

```
LOCUS       RP640_pT_U6_20_4        5929 bp    DNA        circular     29-JAN-2025
DEFINITION  synthetic circular DNA
ACCESSION   .
VERSION     .
KEYWORDS    .
SOURCE      synthetic DNA construct
  ORGANISM  synthetic DNA construct
REFERENCE   1  (bases 1 to 5504)
  AUTHORS   .
  TITLE     Direct Submission
  JOURNAL   Exported Jul 23, 2024 from SnapGene Viewer 7.2.1
            https://www.snapgene.com
COMMENT     [DEFINITION]: Sequence
COMMENT
COMMENT     ApEinfo:methylated:1
FEATURES             Location/Qualifiers
     source          join(1..452,823..877,2825..5929)
                     /note="organism="Sequence" - mol_type="other DNA" -
                     1..5504"
                     /organism="synthetic DNA construct"
                     /locus_tag="organism=Sequence - mol_type=other DNA -
                     1..5504"
                     /label="organism=Sequence - mol_type=other DNA - 1..5504"
                     /ApEinfo_label="organism=Sequence - mol_type=other DNA -
                     1..5504"
                     /ApEinfo_fwdcolor="#00DF87"
                     /ApEinfo_revcolor="#00DF87"
                     /ApEinfo_graphicformat="arrow_data {{0 0.5 0 1 2 0 0 -1 0
                     -0.5} {0 .5 .1 .5 .1 -.5 0 -.5} 0} width 5 offset 0"
                     /ApEinfo_hidden
     misc_feature    complement(join(3..452,823..877,2825..5784))
                     /note="label="lacZ_a" -
                     translation="MSIQHFRVALIPFFAAFCLPVFAHPETLVKVKDAEDQLGARVGY
                     -
                     IELDLNSGKILESFRPEERFPMMSTFKVLLCGAVLSRIDAGQEQLGRRIHYSQNDLVE
                     -
                     YSPVTEKHLTDGMTVRELCSAAITMSDNTAANLLLTTIGGPKELTAFLHNMGDHVTRL
                     -
                     DRWEPELNEAIPNDERDTTMPVAMATTLRKLLTGELLTLASRQQLIDWMEADKVAG"
                     /locus_tag="lacZ_a"
                     /label="lacZ_a"
                     /ApEinfo_label="lacZ_a"
                     /ApEinfo_fwdcolor="#7eff74"
                     /ApEinfo_revcolor="#7eff74"
                     /ApEinfo_graphicformat="arrow_data {{0 0.5 0 1 2 0 0 -1 0
                     -0.5} {0 .5 .1 .5 .1 -.5 0 -.5} 0} width 5 offset 0"
                     /ApEinfo_hidden
     misc_feature    complement(776..791)
                     /locus_tag="EAAGIGILTV"
                     /label="EAAGIGILTV"
                     /ApEinfo_label="EAAGIGILTV"
                     /ApEinfo_fwdcolor="#009192"
                     /ApEinfo_revcolor="#ff7d78"
                     /ApEinfo_graphicformat="arrow_data {{0 1 2 0 0 -1} {} 0}
                     width 5 offset 0"
     misc_feature    22..53
                     /locus_tag="LOR"
                     /label="LOR"
                     /ApEinfo_label="LOR"
                     /ApEinfo_fwdcolor="#7eff74"
                     /ApEinfo_revcolor="#7eff74"
                     /ApEinfo_graphicformat="arrow_data {{0 0.5 0 1 2 0 0 -1 0
                     -0.5} {0 .5 .1 .5 .1 -.5 0 -.5} 0} width 5 offset 0"
     promoter        710..716
                     /vntifkey="29"
                     /locus_tag="hU6"
                     /label="hU6"
                     /ApEinfo_label="hU6"
                     /ApEinfo_fwdcolor="#346ee0"
                     /ApEinfo_revcolor="#346ee0"
                     /ApEinfo_graphicformat="arrow_data {{0 1 2 0 0 -1} {} 0}
                     width 5 offset 0"
     misc_feature    219..250
                     /locus_tag="LIR"
                     /label="LIR"
                     /ApEinfo_label="LIR"
                     /ApEinfo_fwdcolor="#7eff74"
                     /ApEinfo_revcolor="#7eff74"
                     /ApEinfo_graphicformat="arrow_data {{0 0.5 0 1 2 0 0 -1 0
                     -0.5} {0 .5 .1 .5 .1 -.5 0 -.5} 0} width 5 offset 0"
     misc_feature    complement(774..775)
                     /locus_tag="G31"
                     /label="G31"
                     /ApEinfo_label="G31"
                     /ApEinfo_fwdcolor="#ff7d78"
                     /ApEinfo_revcolor="green"
                     /ApEinfo_graphicformat="arrow_data {{0 1 2 0 0 -1} {} 0}
                     width 5 offset 0"
     misc_feature    complement(382..402)
                     /note="label="pBABE_3_primer" - complement(382..402)"
                     /locus_tag="pBABE_3_primer"
                     /label="pBABE_3_primer"
                     /ApEinfo_label="pBABE_3_primer"
                     /ApEinfo_fwdcolor="#7eff74"
                     /ApEinfo_revcolor="#7eff74"
                     /ApEinfo_graphicformat="arrow_data {{0 0.5 0 1 2 0 0 -1 0
                     -0.5} {0 .5 .1 .5 .1 -.5 0 -.5} 0} width 5 offset 0"
     promoter        453..709
                     /vntifkey="29"
                     /locus_tag="hU6(1)"
                     /label="hU6(1)"
                     /ApEinfo_label="hU6"
                     /ApEinfo_fwdcolor="#346ee0"
                     /ApEinfo_revcolor="#346ee0"
                     /ApEinfo_graphicformat="arrow_data {{0 1 2 0 0 -1} {} 0}
                     width 5 offset 0"
     ORF             1418..2134
                     /application_notes="Low fluorescence and low
                     photostability."
                     /description="Blue variant of EGFP generated by
                     mutagenesis"
                     /full_name="Enhanced blue fluorescent protein(ns)"
                     /note="color: #6ddaae; direction: RIGHT"
                     /official_designation="EBFP(ns)"
                     /uuid="c22d3886879b4745b895a6f25c150792"
                     /locus_tag="EBFP(ns)"
                     /label="EBFP(ns)"
                     /ApEinfo_label="EBFP(ns)"
                     /ApEinfo_fwdcolor="#0f7ffe"
                     /ApEinfo_revcolor="pink"
                     /ApEinfo_graphicformat="arrow_data {{0 1 2 0 0 -1} {} 0}
                     width 5 offset 0"
     primer_bind     complement(2835..2856)
                     /locus_tag="pT3_rev"
                     /label="pT3_rev"
                     /ApEinfo_label="pT3_rev"
                     /ApEinfo_fwdcolor="#14c0bd"
                     /ApEinfo_revcolor="#4ec02b"
                     /ApEinfo_graphicformat="arrow_data {{0 0.5 0 1 2 0 0 -1 0
                     -0.5} {0 .5 .1 .5 .1 -.5 0 -.5} 0} width 5 offset 0"
     misc_feature    2845..2864
                     /note="label="EBV_rev_primer" -
                     translation="MTEYKLVVVGAVGVGKSALTIQLIQNHFVDEYDPTIEDSYRKQV
                     -
                     VIDGETCLLDILDTAGQEEYSAMRDQYMRTGEGFLCVFAINNTKSFEDIHHYREQIKR
                     -
                     VKDSEDVPMVLVGNKCDLPSRTVDTKQAQELARSYGIPFIETSAKTRQGVDDAFYTLV
                     - REIRKHKEKMSKDGKKKKKKSRTRCTVM*" - 2420..2439"
                     /locus_tag="EBV_rev_primer"
                     /label="EBV_rev_primer"
                     /ApEinfo_label="EBV_rev_primer"
                     /ApEinfo_fwdcolor="#7eff74"
                     /ApEinfo_revcolor="#7eff74"
                     /ApEinfo_graphicformat="arrow_data {{0 0.5 0 1 2 0 0 -1 0
                     -0.5} {0 .5 .1 .5 .1 -.5 0 -.5} 0} width 5 offset 0"
     Linker          2135..2200
                     /application_notes="Causes co-translational cleavage of
                     the encoded polypeptide. Multiple proteins can be made
                     from a polycistronic transcript containing multiple ORFs
                     separated by 2A. P2A and T2A have higher cleavage
                     efficiency compared to other 2As."
                     /description="Self-cleaving 2A peptide from Porcine
                     teschovirus-1"
                     /full_name="Porcine teschovirus-1 2A peptide"
                     /note="color: #1f36a9; direction: RIGHT"
                     /official_designation="P2A"
                     /uuid="26656daa74294a0f8b331614d72d5f5a"
                     /locus_tag="P2A"
                     /label="P2A"
                     /ApEinfo_label="P2A"
                     /ApEinfo_fwdcolor="#20fefe"
                     /ApEinfo_revcolor="pink"
                     /ApEinfo_graphicformat="arrow_data {{0 1 2 0 0 -1} {} 0}
                     width 5 offset 0"
     primer_bind     3004..3025
                     /locus_tag="RIR Primer"
                     /label="RIR Primer"
                     /ApEinfo_label="RIR Primer"
                     /ApEinfo_fwdcolor="#14c0bd"
                     /ApEinfo_revcolor="#4ec02b"
                     /ApEinfo_graphicformat="arrow_data {{0 0.5 0 1 2 0 0 -1 0
                     -0.5} {0 .5 .1 .5 .1 -.5 0 -.5} 0} width 5 offset 0"
     promoter        883..1387
                     /__SeqFeature__="True"
                     /__level__="0"
                     /application_notes="Medium-strength promoter."
                     /description="['Human phosphoglycerate kinase 1
                     promoter']"
                     /name="hPGK promoter"
                     /note="color: #5566f5; direction: RIGHT"
                     /official_designation="hPGK promoter"
                     /uuid="a105ee8f48814e769c9a41c38cfff41e"
                     /vntifkey="21"
                     /locus_tag="hPGK promoter"
                     /label="hPGK promoter"
                     /ApEinfo_label="hPGK promoter"
                     /ApEinfo_fwdcolor="#346ee0"
                     /ApEinfo_revcolor="#346ee0"
                     /ApEinfo_graphicformat="arrow_data {{0 1 2 0 0 -1} {} 0}
                     width 5 offset 0"
     misc_feature    3032..3062
                     /locus_tag="RIR"
                     /label="RIR"
                     /ApEinfo_label="RIR"
                     /ApEinfo_fwdcolor="#7eff74"
                     /ApEinfo_revcolor="#7eff74"
                     /ApEinfo_graphicformat="arrow_data {{0 0.5 0 1 2 0 0 -1 0
                     -0.5} {0 .5 .1 .5 .1 -.5 0 -.5} 0} width 5 offset 0"
     CDS             2201..2599
                     /__SeqFeature__="True"
                     /__level__="0"
                     /application_notes="Allows cells to be resistant to
                     blasticidin."
                     /description="['Blasticidin resistance gene']"
                     /full_name="Blasticidin resistance gene"
                     /name="Bsd"
                     /note="color: #68e66d; direction: RIGHT"
                     /official_designation="Bsd"
                     /uuid="70e7f83c244f451dae86de92afa03717"
                     /vntifkey="21"
                     /locus_tag="Bsd"
                     /label="Bsd"
                     /ApEinfo_label="Bsd"
                     /ApEinfo_fwdcolor="#e9d024"
                     /ApEinfo_revcolor="#e9d024"
                     /ApEinfo_graphicformat="arrow_data {{0 1 2 0 0 -1} {} 0}
                     width 5 offset 0"
     misc_feature    3229..3260
                     /locus_tag="ROR"
                     /label="ROR"
                     /ApEinfo_label="ROR"
                     /ApEinfo_fwdcolor="#7eff74"
                     /ApEinfo_revcolor="#7eff74"
                     /ApEinfo_graphicformat="arrow_data {{0 0.5 0 1 2 0 0 -1 0
                     -0.5} {0 .5 .1 .5 .1 -.5 0 -.5} 0} width 5 offset 0"
     promoter        complement(3309..3327)
                     /note="label="M13_reverse_primer" -
                     translation="MTEYKLVVVGAVGVGKSALTIQLIQNHFVDEYDPTIEDSYRKQV
                     -
                     VIDGETCLLDILDTAGQEEYSAMRDQYMRTGEGFLCVFAINNTKSFEDIHHYREQIKR
                     -
                     VKDSEDVPMVLVGNKCDLPSRTVDTKQAQELARSYGIPFIETSAKTRQGVDDAFYTLV
                     - REIRKHKEKMSKDGKKKKKKSRTRCTVM*" - complement("
                     /locus_tag="M13_reverse_primer"
                     /label="M13_reverse_primer"
                     /ApEinfo_label="M13_reverse_primer"
                     /ApEinfo_fwdcolor="#346ee0"
                     /ApEinfo_revcolor="#346ee0"
                     /ApEinfo_graphicformat="arrow_data {{0 0.5 0 1 2 0 0 -1 0
                     -0.5} {0 .5 .1 .5 .1 -.5 0 -.5} 0} width 5 offset 0"
     primer_bind     complement(3313..3329)
                     /note="common sequencing primer, one of multiple similar
                     variants"
                     /locus_tag="M13 rev"
                     /label="M13 rev"
                     /ApEinfo_label="M13 rev"
                     /ApEinfo_fwdcolor="#14c0bd"
                     /ApEinfo_revcolor="#4ec02b"
                     /ApEinfo_graphicformat="arrow_data {{0 0.5 0 1 2 0 0 -1 0
                     -0.5} {0 .5 .1 .5 .1 -.5 0 -.5} 0} width 5 offset 0"
     misc_feature    complement(3326..3348)
                     /note="label="M13_pUC_rev_primer" -
                     translation="MTEYKLVVVGAVGVGKSALTIQLIQNHFVDEYDPTIEDSYRKQV
                     -
                     VIDGETCLLDILDTAGQEEYSAMRDQYMRTGEGFLCVFAINNTKSFEDIHHYREQIKR
                     -
                     VKDSEDVPMVLVGNKCDLPSRTVDTKQAQELARSYGIPFIETSAKTRQGVDDAFYTLV
                     - REIRKHKEKMSKDGKKKKKKSRTRCTVM*" - complement("
                     /locus_tag="M13_pUC_rev_primer"
                     /label="M13_pUC_rev_primer"
                     /ApEinfo_label="M13_pUC_rev_primer"
                     /ApEinfo_fwdcolor="#7eff74"
                     /ApEinfo_revcolor="#7eff74"
                     /ApEinfo_graphicformat="arrow_data {{0 0.5 0 1 2 0 0 -1 0
                     -0.5} {0 .5 .1 .5 .1 -.5 0 -.5} 0} width 5 offset 0"
     protein_bind    3337..3353
                     /bound_moiety="lac repressor encoded by lacI"
                     /note="The lac repressor binds to the lac operator to
                     inhibit transcription in E. coli. This inhibition can be
                     relieved by adding lactose or
                     isopropyl-beta-D-thiogalactopyranoside (IPTG)."
                     /locus_tag="lac operator"
                     /label="lac operator"
                     /ApEinfo_label="lac operator"
                     /ApEinfo_fwdcolor="#00D95A"
                     /ApEinfo_revcolor="#00D95A"
                     /ApEinfo_graphicformat="arrow_data {{0 0.5 0 1 2 0 0 -1 0
                     -0.5} {0 .5 .1 .5 .1 -.5 0 -.5} 0} width 5 offset 0"
     promoter        complement(3361..3391)
                     /note="promoter for the E. coli lac operon"
                     /locus_tag="lac promoter"
                     /label="lac promoter"
                     /ApEinfo_label="lac promoter"
                     /ApEinfo_fwdcolor="#346ee0"
                     /ApEinfo_revcolor="#346ee0"
                     /ApEinfo_graphicformat="arrow_data {{0 0.5 0 1 2 0 0 -1 0
                     -0.5} {0 .5 .1 .5 .1 -.5 0 -.5} 0} width 5 offset 0"
     promoter        complement(3362..3391)
                     /note="label="lac_promoter" -
                     translation="MTEYKLVVVGAVGVGKSALTIQLIQNHFVDEYDPTIEDSYRKQV
                     -
                     VIDGETCLLDILDTAGQEEYSAMRDQYMRTGEGFLCVFAINNTKSFEDIHHYREQIKR
                     -
                     VKDSEDVPMVLVGNKCDLPSRTVDTKQAQELARSYGIPFIETSAKTRQGVDDAFYTLV
                     - REIRKHKEKMSKDGKKKKKKSRTRCTVM*" - complement(2937.."
                     /locus_tag="lac_promoter"
                     /label="lac_promoter"
                     /ApEinfo_label="lac_promoter"
                     /ApEinfo_fwdcolor="#346ee0"
                     /ApEinfo_revcolor="#346ee0"
                     /ApEinfo_graphicformat="arrow_data {{0 0.5 0 1 2 0 0 -1 0
                     -0.5} {0 .5 .1 .5 .1 -.5 0 -.5} 0} width 5 offset 0"
     protein_bind    3406..3427
                     /bound_moiety="E. coli catabolite activator protein"
                     /note="CAP binding activates transcription in the presence
                     of cAMP."
                     /locus_tag="CAP binding site"
                     /label="CAP binding site"
                     /ApEinfo_label="CAP binding site"
                     /ApEinfo_fwdcolor="#008065"
                     /ApEinfo_revcolor="#008065"
                     /ApEinfo_graphicformat="arrow_data {{0 0.5 0 1 2 0 0 -1 0
                     -0.5} {0 .5 .1 .5 .1 -.5 0 -.5} 0} width 5 offset 0"
     source          complement(2600..2824)
                     /locus_tag="source"
                     /label="source"
                     /ApEinfo_label="source"
                     /ApEinfo_fwdcolor="#00DF87"
                     /ApEinfo_revcolor="#00DF87"
                     /ApEinfo_graphicformat="arrow_data {{0 0.5 0 1 2 0 0 -1 0
                     -0.5} {0 .5 .1 .5 .1 -.5 0 -.5} 0} width 5 offset 0"
                     /ApEinfo_hidden
     rep_origin      complement(3715..4303)
                     /direction=LEFT
                     /note="high-copy-number ColE1/pMB1/pBR322/pUC origin of
                     replication"
                     /locus_tag="ori"
                     /label="ori"
                     /ApEinfo_label="ori"
                     /ApEinfo_fwdcolor="#999999"
                     /ApEinfo_revcolor="#999999"
                     /ApEinfo_graphicformat="arrow_data {{0 0.5 0 1 2 0 0 -1 0
                     -0.5} {0 .5 .1 .5 .1 -.5 0 -.5} 0} width 5 offset 0"
     gene            complement(4474..5334)
                     /note="label="Ampicillin" - gene="Ampicillin" -
                     translation="MTEYKLVVVGAVGVGKSALTIQLIQNHFVDEYDPTIEDSYRKQV
                     -
                     VIDGETCLLDILDTAGQEEYSAMRDQYMRTGEGFLCVFAINNTKSFEDIHHYREQIKR
                     -
                     VKDSEDVPMVLVGNKCDLPSRTVDTKQAQELARSYGIPFIETSAKTRQGVDDAFYTLV
                     - REIRKHKEKMSKDGKKKKKKSRTRCTVM*" -"
                     /locus_tag="AmpR"
                     /label="AmpR"
                     /ApEinfo_label="AmpR"
                     /ApEinfo_fwdcolor="#ff797d"
                     /ApEinfo_revcolor="#ff102f"
                     /ApEinfo_graphicformat="arrow_data {{0 0.5 0 1 2 0 0 -1 0
                     -0.5} {0 .5 .1 .5 .1 -.5 0 -.5} 0} width 5 offset 0"
     CDS             complement(4474..5334)
                     /note="label="ORF frame 2" -
                     translation="MSIQHFRVALIPFFAAFCLPVFAHPETLVKVKDAEDQLGARVGY
                     -
                     IELDLNSGKILESFRPEERFPMMSTFKVLLCGAVLSRIDAGQEQLGRRIHYSQNDLVE
                     -
                     YSPVTEKHLTDGMTVRELCSAAITMSDNTAANLLLTTIGGPKELTAFLHNMGDHVTRL
                     - DRWEPELNEAIPNDERDTTMPVAMATTLRKLLTGELLTLASRQQLIDWMEA"
                     /locus_tag="ORF frame 2"
                     /label="ORF frame 2"
                     /ApEinfo_label="ORF frame 2"
                     /ApEinfo_fwdcolor="#e9d024"
                     /ApEinfo_revcolor="#e9d024"
                     /ApEinfo_graphicformat="arrow_data {{0 0.5 0 1 2 0 0 -1 0
                     -0.5} {0 .5 .1 .5 .1 -.5 0 -.5} 0} width 5 offset 0"
     promoter        complement(5335..5439)
                     /gene="bla"
                     /locus_tag="AmpR promoter"
                     /label="AmpR promoter"
                     /ApEinfo_label="AmpR promoter"
                     /ApEinfo_fwdcolor="#346ee0"
                     /ApEinfo_revcolor="#346ee0"
                     /ApEinfo_graphicformat="arrow_data {{0 0.5 0 1 2 0 0 -1 0
                     -0.5} {0 .5 .1 .5 .1 -.5 0 -.5} 0} width 5 offset 0"
     promoter        complement(5376..5404)
                     /note="label="AmpR_promoter" -
                     translation="MSIQHFRVALIPFFAAFCLPVFAHPETLVKVKDAEDQLGARVGY
                     -
                     IELDLNSGKILESFRPEERFPMMSTFKVLLCGAVLSRIDAGQEQLGRRIHYSQNDLVE
                     -
                     YSPVTEKHLTDGMTVRELCSAAITMSDNTAANLLLTTIGGPKELTAFLHNMGDHVTRL
                     - DRWEPELNEAIPNDERDTTMPVAMATTLRKLLTGELLTLASRQQLIDWM"
                     /locus_tag="AmpR_promoter"
                     /label="AmpR_promoter"
                     /ApEinfo_label="AmpR_promoter"
                     /ApEinfo_fwdcolor="#346ee0"
                     /ApEinfo_revcolor="#346ee0"
                     /ApEinfo_graphicformat="arrow_data {{0 0.5 0 1 2 0 0 -1 0
                     -0.5} {0 .5 .1 .5 .1 -.5 0 -.5} 0} width 5 offset 0"
     misc_feature    complement(5563..5585)
                     /note="label="pGEX_3_primer" -
                     translation="MSIQHFRVALIPFFAAFCLPVFAHPETLVKVKDAEDQLGARVGY
                     -
                     IELDLNSGKILESFRPEERFPMMSTFKVLLCGAVLSRIDAGQEQLGRRIHYSQNDLVE
                     -
                     YSPVTEKHLTDGMTVRELCSAAITMSDNTAANLLLTTIGGPKELTAFLHNMGDHVTRL
                     - DRWEPELNEAIPNDERDTTMPVAMATTLRKLLTGELLTLASRQQLIDWM"
                     /locus_tag="pGEX_3_primer"
                     /label="pGEX_3_primer"
                     /ApEinfo_label="pGEX_3_primer"
                     /ApEinfo_fwdcolor="#7eff74"
                     /ApEinfo_revcolor="#7eff74"
                     /ApEinfo_graphicformat="arrow_data {{0 0.5 0 1 2 0 0 -1 0
                     -0.5} {0 .5 .1 .5 .1 -.5 0 -.5} 0} width 5 offset 0"
     misc_feature    2600..2824
                     /__SeqFeature__="True"
                     /__level__="0"
                     /application_notes="Allows transcription termination and
                     polyadenylation of mRNA transcribed by Pol II RNA
                     polymerase."
                     /full_name="Bovine growth hormone polyadenylation"
                     /name="BGH pA"
                     /note="color: #46c6ef; direction: LEFT"
                     /official_designation="BGH pA"
                     /uuid="439e92d8cb8e406dbfacd87ce28d14dc"
                     /vntifkey="21"
                     /locus_tag="BGH pA"
                     /label="BGH pA"
                     /ApEinfo_label="BGH pA"
                     /ApEinfo_fwdcolor="#7eff74"
                     /ApEinfo_revcolor="#7eff74"
                     /ApEinfo_graphicformat="arrow_data {{0 0.5 0 1 2 0 0 -1 0
                     -0.5} {0 .5 .1 .5 .1 -.5 0 -.5} 0} width 5 offset 0"
     misc_feature    5898..5920
                     /note="label="M13_pUC_fwd_primer" -
                     translation="MSIQHFRVALIPFFAAFCLPVFAHPETLVKVKDAEDQLGARVGY
                     -
                     IELDLNSGKILESFRPEERFPMMSTFKVLLCGAVLSRIDAGQEQLGRRIHYSQNDLVE
                     -
                     YSPVTEKHLTDGMTVRELCSAAITMSDNTAANLLLTTIGGPKELTAFLHNMGDHVTRL
                     - DRWEPELNEAIPNDERDTTMPVAMATTLRKLLTGELLTLASRQQ"
                     /locus_tag="M13_pUC_fwd_primer"
                     /label="M13_pUC_fwd_primer"
                     /ApEinfo_label="M13_pUC_fwd_primer"
                     /ApEinfo_fwdcolor="#7eff74"
                     /ApEinfo_revcolor="#7eff74"
                     /ApEinfo_graphicformat="arrow_data {{0 0.5 0 1 2 0 0 -1 0
                     -0.5} {0 .5 .1 .5 .1 -.5 0 -.5} 0} width 5 offset 0"
     promoter        5913..5929
                     /note="label="M13_forward20_primer" -
                     translation="MSIQHFRVALIPFFAAFCLPVFAHPETLVKVKDAEDQLGARVGY
                     -
                     IELDLNSGKILESFRPEERFPMMSTFKVLLCGAVLSRIDAGQEQLGRRIHYSQNDLVE
                     -
                     YSPVTEKHLTDGMTVRELCSAAITMSDNTAANLLLTTIGGPKELTAFLHNMGDHVTRL
                     - DRWEPELNEAIPNDERDTTMPVAMATTLRKLLTGELLTLASR"
                     /locus_tag="M13_forward20_primer"
                     /label="M13_forward20_primer"
                     /ApEinfo_label="M13_forward20_primer"
                     /ApEinfo_fwdcolor="#346ee0"
                     /ApEinfo_revcolor="#346ee0"
                     /ApEinfo_graphicformat="arrow_data {{0 0.5 0 1 2 0 0 -1 0
                     -0.5} {0 .5 .1 .5 .1 -.5 0 -.5} 0} width 5 offset 0"
     primer_bind     5913..5929
                     /note="common sequencing primer, one of multiple similar
                     variants"
                     /locus_tag="M13 fwd"
                     /label="M13 fwd"
                     /ApEinfo_label="M13 fwd"
                     /ApEinfo_fwdcolor="#14c0bd"
                     /ApEinfo_revcolor="#4ec02b"
                     /ApEinfo_graphicformat="arrow_data {{0 0.5 0 1 2 0 0 -1 0
                     -0.5} {0 .5 .1 .5 .1 -.5 0 -.5} 0} width 5 offset 0"
     misc_feature    complement(776..776)
                     /locus_tag="G31(1)"
                     /label="G31(1)"
                     /ApEinfo_label="G31"
                     /ApEinfo_fwdcolor="#ff7d78"
                     /ApEinfo_revcolor="green"
                     /ApEinfo_graphicformat="arrow_data {{0 1 2 0 0 -1} {} 0}
                     width 5 offset 0"
     variation       complement(776..776)
                     /locus_tag="G-to-A mutation (target A)"
                     /label="G-to-A mutation (target A)"
                     /ApEinfo_label="G-to-A mutation (target A)"
                     /ApEinfo_fwdcolor="#fe0300"
                     /ApEinfo_revcolor="#ff9e39"
                     /ApEinfo_graphicformat="arrow_data {{0 0.5 0 1 2 0 0 -1 0
                     -0.5} {0 .5 .1 .5 .1 -.5 0 -.5} 0} width 5 offset 0"
     terminator      793..798
                     /locus_tag="PolIII terminator"
                     /label="PolIII terminator"
                     /ApEinfo_label="PolIII terminator"
                     /ApEinfo_fwdcolor="#9d1b1c"
                     /ApEinfo_revcolor="#9d1b1c"
                     /ApEinfo_graphicformat="arrow_data {{0 1 2 0 0 -1} {} 0}
                     width 5 offset 0"
     misc_binding    739..742
                     /bound_moiety=""
                     /locus_tag="4nt mismatch (as in Uzonyi et al 2021)"
                     /label="4nt mismatch (as in Uzonyi et al 2021)"
                     /ApEinfo_label="4nt mismatch (as in Uzonyi et al 2021)"
                     /ApEinfo_fwdcolor="#ff4809"
                     /ApEinfo_revcolor="#ff4809"
                     /ApEinfo_graphicformat="arrow_data {{0 0.5 0 1 2 0 0 -1 0
                     -0.5} {0 .5 .1 .5 .1 -.5 0 -.5} 0} width 5 offset 0"
     misc_feature    complement(762..775)
                     /locus_tag="EAAGIGILTV(1)"
                     /label="EAAGIGILTV(1)"
                     /ApEinfo_label="EAAGIGILTV"
                     /ApEinfo_fwdcolor="#009192"
                     /ApEinfo_revcolor="#ff7d78"
                     /ApEinfo_graphicformat="arrow_data {{0 1 2 0 0 -1} {} 0}
                     width 5 offset 0"
ORIGIN
        1 gaattcgagc tcggtaccct acagttgaag tcggaagttt acatacactt aagttggagt
       61 cattaaaact cgtttttcaa ctactccaca aatttcttgt taacaaacaa tagttttggc
      121 aagtcagtta ggacatctac tttgtgcatg acacaagtca tttttccaac aattgtttac
      181 agacagatta tttcacttat aattcactgt atcacaattc cagtgggtca gaagtttaca
      241 tacactaagt tgactgtgcc tttaaacagc ttggaaaatt ccagaaaatg atgtcatggc
      301 tttagaagct tctgatagac taattgacat catttgagtc aattggaggt gtacctgtgg
      361 atgtatttca aggaattctg tggaatgtgt gtcagttagg gtgtggaaag tccccaggct
      421 ccccagcagg cagaagtatg caaagcatgc ataaggtcgg gcaggaagag ggcctatttc
      481 ccatgattcc ttcatatttg catatacgat acaaggctgt tagagagata attagaatta
      541 atttgactgt aaacacaaag atattagtac aaaatacgtg acgtagaaag taataatttc
      601 ttgggtagtt tgcagtttta aaattatgtt ttaaaatgga ctatcatatg cttaccgtaa
      661 cttgaaagta tttcgatttc ttggctttat atatcttgtg gaaaggacga aacaccGGtt
      721 ctacaatacc aacagccgTA CTgcagtaag actcccagga tcactgtcag gatgcCgatc
      781 ccagcggcct ctttttttct agacccagct ttcttgtaca aactcgagaa taaaggtacc
      841 ttattttcat tagatttgtg tgttggtttt ttgtgtgCTA GCgggttgcg ccttttccaa
      901 ggcagccctg ggtttgcgca gggacgcggc tgctctgggc gtggttccgg gaaacgcagc
      961 ggcgccgacc ctgggtctcg cacattcttc acgtccgttc gcagcgtcac ccggatcttc
     1021 gccgctaccc ttgtgggccc cccggcgacg cttcctgctc cgcccctaag tcgggaaggt
     1081 tccttgcggt tcgcggcgtg ccggacgtga caaacggaag ccgcacgtct cactagtacc
     1141 ctcgcagacg gacagcgcca gggagcaatg gcagcgcgcc gaccgcgatg ggctgtggcc
     1201 aatagcggct gctcagcagg gcgcgccgag agcagcggcc gggaaggggc ggtgcgggag
     1261 gcggggtgtg gggcggtagt gtgggccctg ttcctgcccg cgcggtgttc cgcattctgc
     1321 aagcctccgg agcgcacgtc ggcagtcggc tccctcgttg accgaatcac cgacctctct
     1381 ccccaggcaa gtttgtacaa aaaagcaggc tgccaccatg gtgagcaagg gcgaggagct
     1441 gttcaccggg gtggtgccca tcctggtcga gctggacggc gacgtaaacg gccacaagtt
     1501 cagcgtgtcc ggcgagggcg agggcgatgc cacctacggc aagctgaccc tgaagttcat
     1561 ctgcaccacc ggcaagctgc ccgtgccctg gcccaccctc gtgaccaccc tgacccacgg
     1621 cgtgcagtgc ttcagccgct accccgacca catgaagcag cacgacttct tcaagtccgc
     1681 catgcccgaa ggctacgtcc aggagcgcac catcttcttc aaggacgacg gcaactacaa
     1741 gacccgcgcc gaggtgaagt tcgagggcga caccctggtg aaccgcatcg agctgaaggg
     1801 catcgacttc aaggaggacg gcaacatcct ggggcacaag ctggagtaca acttcaacag
     1861 ccacaacgtc tatatcatgg ccgacaagca gaagaacggc atcaaggtga acttcaagat
     1921 ccgccacaac atcgaggacg gcagcgtgca gctcgccgac cactaccagc agaacacccc
     1981 catcggcgac ggccccgtgc tgctgcccga caaccactac ctgagcaccc agtccgccct
     2041 gagcaaagac cccaacgaga agcgcgatca catggtcctg ctggagttcg tgaccgccgc
     2101 cgggatcact ctcggcatgg acgagctgta caagggaagc ggagccacga acttctctct
     2161 gttaaagcaa gcaggagatg ttgaagaaaa ccccgggcct atggccaagc ctttgtctca
     2221 agaagaatcc accctcattg aaagagcaac ggctacaatc aacagcatcc ccatctctga
     2281 agactacagc gtcgccagcg cagctctctc tagcgacggc cgcatcttca ctggtgtcaa
     2341 tgtatatcat tttactgggg gaccttgtgc agaactcgtg gtgctgggca ctgctgctgc
     2401 tgcggcagct ggcaacctga cttgtatcgt cgcgatcgga aatgagaaca ggggcatctt
     2461 gagcccctgc ggacggtgcc gacaggtgct tctcgatctg catcctggga tcaaagccat
     2521 agtgaaggac agtgatggac agccgacggc agttgggatt cgtgaattgc tgccctctgg
     2581 ttatgtgtgg gagggctaac tgtgccttct agttgccagc catctgttgt ttgcccctcc
     2641 cccgtgcctt ccttgaccct ggaaggtgcc actcccactg tcctttccta ataaaatgag
     2701 gaaattgcat cgcattgtct gagtaggtgt cattctattc tggggggtgg ggtggggcag
     2761 gacagcaagg gggaggattg ggaagacaat agcaggcatg ctggggatgc ggtgggctct
     2821 atgggcggcc gctgcattct agttgtggtt tgtccaaact catcaatgta tcttatcatg
     2881 tctggatccc atcacaaagc tctgacctca atcctataga aaggaggaat gagccaaaat
     2941 tcacccaact tattgtggga agcttgtgga aggctactcg aaatgtttga cccaagttaa
     3001 acaatttaaa ggcaatgcta ccaaatacta attgagtgta tgttaacttc tgacccactg
     3061 ggaatgtgat gaaagaaata aaagctgaaa tgaatcattc tctctactat tattctgata
     3121 tttcacattc ttaaaataaa gtggtgatcc taactgacct taagacaggg aatctttact
     3181 cggattaaat gtcaggaatt gtgaaaaagt gagtttaaat gtatttggct aaggtgtatg
     3241 taaacttccg acttcaactg taggggatcc tctagagtcg acctgcaggc atgcaagctt
     3301 ggcgtaatca tggtcatagc tgtttcctgt gtgaaattgt tatccgctca caattccaca
     3361 caacatacga gccggaagca taaagtgtaa agcctggggt gcctaatgag tgagctaact
     3421 cacattaatt gcgttgcgct cactgcccgc tttccagtcg ggaaacctgt cgtgccagct
     3481 gcattaatga atcggccaac gcgcggggag aggcggtttg cgtattgggc gctcttccgc
     3541 ttcctcgctc actgactcgc tgcgctcggt cgttcggctg cggcgagcgg tatcagctca
     3601 ctcaaaggcg gtaatacggt tatccacaga atcaggggat aacgcaggaa agaacatgtg
     3661 agcaaaaggc cagcaaaagg ccaggaaccg taaaaaggcc gcgttgctgg cgtttttcca
     3721 taggctccgc ccccctgacg agcatcacaa aaatcgacgc tcaagtcaga ggtggcgaaa
     3781 cccgacagga ctataaagat accaggcgtt tccccctgga agctccctcg tgcgctctcc
     3841 tgttccgacc ctgccgctta ccggatacct gtccgccttt ctcccttcgg gaagcgtggc
     3901 gctttctcat agctcacgct gtaggtatct cagttcggtg taggtcgttc gctccaagct
     3961 gggctgtgtg cacgaacccc ccgttcagcc cgaccgctgc gccttatccg gtaactatcg
     4021 tcttgagtcc aacccggtaa gacacgactt atcgccactg gcagcagcca ctggtaacag
     4081 gattagcaga gcgaggtatg taggcggtgc tacagagttc ttgaagtggt ggcctaacta
     4141 cggctacact agaagaacag tatttggtat ctgcgctctg ctgaagccag ttaccttcgg
     4201 aaaaagagtt ggtagctctt gatccggcaa acaaaccacc gctggtagcg gtggtttttt
     4261 tgtttgcaag cagcagatta cgcgcagaaa aaaaggatct caagaagatc ctttgatctt
     4321 ttctacgggg tctgacgctc agtggaacga aaactcacgt taagggattt tggtcatgag
     4381 attatcaaaa aggatcttca cctagatcct tttaaattaa aaatgaagtt ttaaatcaat
     4441 ctaaagtata tatgagtaaa cttggtctga cagttaccaa tgcttaatca gtgaggcacc
     4501 tatctcagcg atctgtctat ttcgttcatc catagttgcc tgactccccg tcgtgtagat
     4561 aactacgata cgggagggct taccatctgg ccccagtgct gcaatgatac cgcgagaccc
     4621 acgctcaccg gctccagatt tatcagcaat aaaccagcca gccggaaggg ccgagcgcag
     4681 aagtggtcct gcaactttat ccgcctccat ccagtctatt aattgttgcc gggaagctag
     4741 agtaagtagt tcgccagtta atagtttgcg caacgttgtt gccattgcta caggcatcgt
     4801 ggtgtcacgc tcgtcgtttg gtatggcttc attcagctcc ggttcccaac gatcaaggcg
     4861 agttacatga tcccccatgt tgtgcaaaaa agcggttagc tccttcggtc ctccgatcgt
     4921 tgtcagaagt aagttggccg cagtgttatc actcatggtt atggcagcac tgcataattc
     4981 tcttactgtc atgccatccg taagatgctt ttctgtgact ggtgagtact caaccaagtc
     5041 attctgagaa tagtgtatgc ggcgaccgag ttgctcttgc ccggcgtcaa tacgggataa
     5101 taccgcgcca catagcagaa ctttaaaagt gctcatcatt ggaaaacgtt cttcggggcg
     5161 aaaactctca aggatcttac cgctgttgag atccagttcg atgtaaccca ctcgtgcacc
     5221 caactgatct tcagcatctt ttactttcac cagcgtttct gggtgagcaa aaacaggaag
     5281 gcaaaatgcc gcaaaaaagg gaataagggc gacacggaaa tgttgaatac tcatactctt
     5341 cctttttcaa tattattgaa gcatttatca gggttattgt ctcatgagcg gatacatatt
     5401 tgaatgtatt tagaaaaata aacaaatagg ggttccgcgc acatttcccc gaaaagtgcc
     5461 acctgacgtc taagaaacca ttattatcat gacattaacc tataaaaata ggcgtatcac
     5521 gaggcccttt cgtctcgcgc gtttcggtga tgacggtgaa aacctctgac acatgcagct
     5581 cccggagacg gtcacagctt gtctgtaagc ggatgccggg agcagacaag cccgtcaggg
     5641 cgcgtcagcg ggtgttggcg ggtgtcgggg ctggcttaac tatgcggcat cagagcagat
     5701 tgtactgaga gtgcaccata tgcggtgtga aataccgcac agatgcgtaa ggagaaaata
     5761 ccgcatcagg cgccattcgc cattcaggct gcgcaactgt tgggaagggc gatcggtgcg
     5821 ggcctcttcg ctattacgcc agctggcgaa agggggatgt gctgcaaggc gattaagttg
     5881 ggtaacgcca gggttttccc agtcacgacg ttgtaaaacg acggccagt
//
```

d. **References**

1.	Qu, L. *et al.* Programmable RNA editing by recruiting endogenous ADAR using engineered RNAs. *Nat Biotechnol* **37**, 1059–1069 (2019).

2.	Uzonyi, A. *et al.* Deciphering the principles of the RNA editing code via large-scale systematic probing. *Mol Cell* **81**, 2374-2387.e3 (2021).

3.	Doman, J. L., Raguram, A., Newby, G. A. & Liu, D. R. Evaluation and minimization of Cas9-independent off-target DNA editing by cytosine base editors. *Nat Biotechnol* **38**, 620–628 (2020).

4.	Kluesner, M. *et al.* MultiEditR: The first tool for detection and quantification of multiple RNA editing sites from Sanger sequencing demonstrates comparable fidelity to RNA-seq. *Molecular Therapy - Nucleic Acids* https://doi.org/10.1016/j.omtn.2021.07.008 (2021) doi:10.1016/j.omtn.2021.07.008.
